# Epistasis, inbreeding depression and the evolution of self-fertilization

**DOI:** 10.1101/809814

**Authors:** Diala Abu Awad, Denis Roze

**Author notes:** Address for correspondence: Denis Roze, Station Biologique de Roscoff, Place Georges Teissier, CS90074, 29688 Roscoff Cedex, France.

## Abstract

Inbreeding depression resulting from partially recessive deleterious alleles is thought to be the main genetic factor preventing self-fertilizing mutants from spreading in outcrossing hermaphroditic populations. However, deleterious alleles may also generate an advantage to selfers in terms of more efficient purging, while the effects of epistasis among those alleles on inbreeding depression and mating system evolution remain little explored. In this paper, we use a general model of selection to disentangle the effects of different forms of epistasis (additive-by-additive, additive-by-dominance and dominance-by-dominance) on inbreeding depression and on the strength of selection for selfing. Models with fixed epistasis across loci, and models of stabilizing selection acting on quantitative traits (generating distributions of epistasis) are considered as special cases. Besides its effects on inbreeding depression, epistasis may increase the purging advantage associated with selfing (when it is negative on average), while the variance in epistasis favors selfing through the generation of linkage disequilibria that increase mean fitness. Approximations for the strengths of these effects are derived, and compared with individual-based simulation results.

## INTRODUCTION

Self-fertilization is a widespread mating system found in hermaphroditic plants and animals (e.g., Jarne and Auld, 2006; Igic and Busch, 2013). In Angiosperms, the transition from outcrossing to selfing occurred multiple times, leading to approximately 10−15% of species self-fertilizing at very high rates (Barrett et al., 2014). Two possible benefits of selfing have been proposed to explain such transitions: the possibility for a single individual to generate offspring in the absence of mating partner or pollinator (“reproductive assurance”, Darwin, 1876; Stebbins, 1957; Porcher and Lande, 2005a; Busch and Delph, 2012), and the “automatic advantage” stemming from the fact that, in a population containing both selfers and outcrossers, selfers tend to transmit more copies of their genome to the next generation if they continue to export pollen — thus retaining the ability to sire outcrossed ovules (Fisher, 1941; Charlesworth, 1980; Stone et al., 2014). The main evolutionary force thought to oppose the spread of selfing is inbreeding depression, the decreased fitness of inbred offspring resulting from the expression of partially recessive deleterious alleles segregating within populations (Charlesworth and Charlesworth, 1987). When selfers export as much pollen as out-crossers (leading to a 50% transmission advantage for selfing), inbreeding depression must be 0.5 to compensate for the automatic advantage of selfing (Lande and Schemske, 1985). However, observations from natural populations indicate that self-fertilizing individuals do not always export as much pollen as their outcrossing counterparts, as some of their pollen production is used to fertilize their own ovules (see references in Porcher and Lande, 2005a). This phenomenon, known as pollen discounting, decreases the automatic advantage of selfing (Nagylaki, 1976; Charlesworth, 1980), thus reducing the threshold value of inbreeding depression above which outcrossing can be maintained (e.g., Holsinger et al., 1984). It may also lead to evolutionarily stable mixed mating systems (involving both selfing and outcrossing) under some models of discounting such as the mass-action pollination model (Holsinger, 1991; Porcher and Lande, 2005a).

Several models explored the evolution of mating systems while explicitly representing the genetic architecture of inbreeding depression (e.g., Charlesworth et al., 1990; Uyenoyama and Waller, 1991; Epinat and Lenormand, 2009; Porcher and Lande, 2005b; Gervais et al., 2014), and highlighted the importance of another genetic factor (besides the automatic advantage and inbreeding depression) affecting the evolution of selfing. This third factor stems from the fact that selection against deleterious alleles is more efficient among selfed offspring (due to their increased homozygosity) than among outcrossed offspring, generating positive linkage disequilibria between alleles increasing the selfing rate and the more advantageous alleles at selected loci. Alleles increasing selfing thus tend to be found on better purged genetic backgrounds, which may allow selfing to spread even when inbreeding depression is higher than 0.5 (Charlesworth et al., 1990). This effect becomes more important as the strength of selection against deleterious alleles increases (so that purging occurs more rapidly), recombination decreases, and as alleles increasing selfing have larger effects — so that linkage disequilibria can be maintained over larger numbers of generations (Charlesworth et al., 1990; Uyenoyama and Waller, 1991; Epinat and Lenormand, 2009). This corresponds to Lande and Schemske’s (1985) verbal prediction that a mutant allele coding for complete selfing may increase in frequency regardless of the amount of inbreeding depression.

Most genetic models on the evolution of selfing assume that deleterious alleles at different loci have multiplicative effects (no epistasis). Charlesworth et al. (1991) considered a deterministic model including synergistic epistasis between deleterious alleles, showing that this form of epistasis tends to flatten the relation between inbreeding depression and the population’s selfing rate, inbreeding depression sometimes increasing at high selfing rates. Concerning the spread of selfing modifier alleles, the results were qualitatively similar to the multiplicative model, except that, for parameter values where full outcrossing is not stable, the evolutionarily stable selfing rate tended to be slightly below 1 under synergistic epistasis (whereas it would have been at exactly 1 in the absence of epistasis). Other models explored the effect of partial selfing on inbreeding depression generated by polygenic quantitative traits under stabilizing selection (Lande and Porcher, 2015; Abu Awad and Roze, 2018). This type of model typically generates distributions of epistatic interactions across loci, including possible compensatory effects between mutations. When effective recombination is sufficiently weak, linkage disequilibria generated by epistasis may greatly reduce inbreeding depression, and even generate outbreeding depression between selfing lineages carrying different combinations of compensatory mutations. However, the evolution of the selfing rate was not considered by these models.

In this paper, we use a general model of epistasis between pairs of selected loci to explore the effects of epistasis on inbreeding depression and on the evolution of selfing. We derive analytical approximations showing that epistatic interactions affect the spread of selfing modifiers through various mechanisms: by affecting inbreeding depression, the purging advantage of selfers and also through linkage disequilibria between selected loci. Although the expressions obtained can become complicated for intermediate selfing rates, we will see that the condition determining whether selfing can spread in a fully outcrossing population often remains relatively simple. Notably, our model allows us to disentangle the effects of additive-by-additive, additive-by-dominance and dominance-by-dominance epistatic interactions on inbreeding depression and selection for selfing — while the models used by Charlesworth et al. (1991), Lande and Porcher (2015) and Abu Awad and Roze (2018) impose certain relations between these quantities. The cases of fixed, synergistic epistasis and of stabilizing selection acting on quantitative traits (Fisher’s geometric model) will be considered as special cases, for which we will also present individual-based simulation results. Overall, our results show that, for a given level of inbreeding depression and average strength of selection against deleterious alleles, epistatic interactions tend to facilitate the spread of selfing, due to the fact that selfing can maintain beneficial combinations of alleles.

## METHODS

### Life cycle

Our analytical model represents an infinite, hermaphroditic population with discrete generations. A proportion *σ* of ovules produced by a given individual are self-fertilized, while its remaining ovules are fertilized by pollen sampled from the population pollen pool (Table 1 provides a list of the symbols used throughout the paper). A parameter *κ* represents the rate of pollen discounting: an individual with selfing rate *σ* contributes to the pollen pool in proportion 1 − *κσ* (e.g., Charlesworth, 1980). Therefore, *κ* equals 0 in the absence of pollen discounting, while *κ* equals 1 under full discounting (in which case complete selfers do not contribute to the pollen pool). We assume that the selfing rate *σ* is genetically variable, and coded by *ℓ*_*σ*_ loci with additive effects:

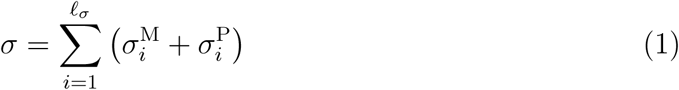

where the sum is over all loci affecting the selfing rate, and where 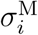 and 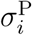 represent the effect of the alleles present respectively on the maternally and paternally inherited genes at locus *i* (note that the assumption of additivity within and between loci may not always hold, in particular when selfing rates are close to 0 or 1). The model does not make any assumption concerning the number of alleles segregating at loci affecting the selfing rate; however, our analysis will assume that the variance of *σ* in the population remains small and that linkage disequilibria between loci affecting the selfing rate may be neglected, effectively leading to the same expression for the selection gradient on the selfing rate as in a simpler model considering the spread of a mutant allele changing *σ* by a small amount. Although we assume that the selfing rate is purely genetically determined, our general results should still hold when *σ* is also affected by (uncorrelated) environmental effects, after multiplying expressions for the change in the average selfing rate over time by the heritability of *σ*.

**Table 1:** Parameters and variables of the model.

|  |  |
| --- | --- |
| $\sigma$ | Selfing rate |
| $\bar{\sigma}, V_{\sigma}$ | Mean and variance in the selfing rate in the population |
| $\kappa$ | Rate of pollen discounting |
| $\ell_{\sigma}$ | Number of loci affecting the selfing rate |
| $W, \bar{W}$ | Fitness of an individual, and average fitness |
| $\ell$ | Number of loci affecting fitness |
| $U$ | Overall (haploid) mutation rate at loci affecting fitness |
| $p_j, q_j$ | Frequencies of alleles 1 and 0 at loci affecting fitness |
| $\ell$ | Number of loci affecting selected traits |
| $n_d$ | Mean number of deleterious alleles per haploid genome |
| $s, h$ | Selection and dominance coefficients of deleterious alleles |
| $e_{\text{axa}}, e_{\text{axd}}, e_{\text{dxd}}$ | Additive-by-additive, additive-by-dominance and dominance-by-dominance epistasis between deleterious alleles |
| $\beta$ | Strength of synergistic epistasis in Charlesworth et al.'s (1991) model |
| $n$ | Number of quantitative traits under stabilizing selection |
| $V_s$ | Strength of stabilizing selection |
| $r_{\alpha j}$ | Effect of allele 1 at locus $j$ on trait $\alpha$ |
| $\varsigma_j$ | Fitness effect of a heterozygous mutation in an optimal genotype (stabilizing selection model) |
| $\bar{\varsigma}$ | Average fitness effect of a heterozygous mutation in an optimal genotype (stabilizing selection model) |
| $a^2$ | Variance of mutational effects on traits under stabilizing selection |
| $Q$ | Shape of the fitness peak (equation 15) |
| $a_{\mathbb{U},\mathbb{V}}$ | Effect of selection on the sets $\mathbb{U}$ and $\mathbb{V}$ of loci present on the maternally and paternally inherited haplotypes of an individual (equation 8) |
| $D_{\mathbb{U},\mathbb{V}}$ | Genetic association between the sets $\mathbb{U}$ and $\mathbb{V}$ of loci present on the maternally and paternally inherited haplotypes of an individual (equation 4) |
| $\rho_{jk}$ | Recombination rate between loci $j$ and $k$ |
| $U_{\text{self}}$ | Mutation rate at loci affecting the selfing rate |
| $\sigma_{\text{self}}^2$ | Variance of mutational effects at loci affecting the selfing rate |
| $\delta$ | Inbreeding depression |
| $\delta'$ | Inbreeding depression measured after selection |
| $F$ | Inbreeding coefficient |
| $G_{jk}$ | Identity disequilibrium between loci $j$ and $k$ |
| $G$ | Identity disequilibrium between freely recombining loci |

The fitness *W* of an organism is defined as its overall fecundity (that may depend on its survival), so that the expected number of seeds produced by an individual is proportional to *W*, while its contribution to the population pollen pool is proportional to *W* (1 − *κσ*). We assume that *W* is affected by a possibly large number *ℓ* of biallelic loci. Alleles at each of these loci are denoted 0 and 1; we assume an equal mutation rate *u* from 0 to 1 and from 1 to 0, assumed to be small relative to the strength of selection at each locus. The overall mutation rate (per haploid genome) at loci affecting fitness is denoted *U* = *uℓ*. The quantity 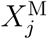 (resp. 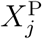) equals 0 if the individual carries allele 0 on its maternally (resp. paternally) inherited copy of locus *j*, and equals 1 otherwise. The frequencies of allele 1 at locus *j* on the maternally and paternally inherited genes (averages of 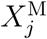 and 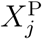 over the whole population) are denoted 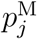 and 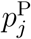. Finally, 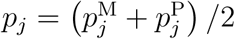 is the frequency of allele 1 at locus *j* in the whole population.

### Genetic associations

Throughout the paper, index *i* will denote a locus affecting the selfing rate of individuals, while indices *j* and *k* will denote loci affecting fitness. Following Barton and Turelli (1991) and Kirkpatrick et al. (2002), we define the centered variables:

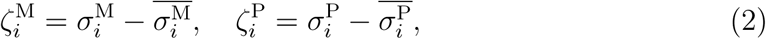

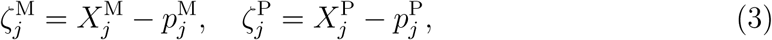

where 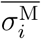 and 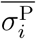 are the averages of 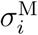 and 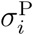 over the whole population. The genetic association between the sets 𝕌 and 𝕍 of loci present in the maternally and paternally derived genome of an individual is defined as:

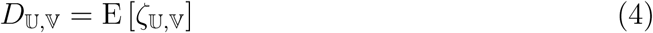

where E stands for the average over all individuals in the population, and with:

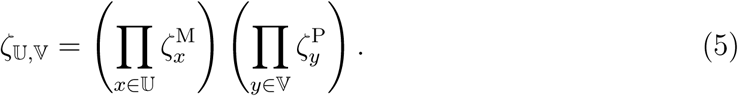

For example, 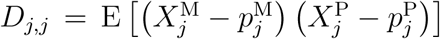 is a measure of the departure from Hardy-Weinberg equilibrium at locus *j*, while 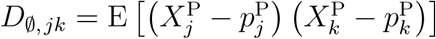 measures the linkage disequilibrium between loci *j* and *k* on paternally derived haplotypes. Finally, 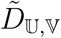 is defined as (*D*_𝕌,𝕍_ + *D*_𝕍,𝕌_) /2, and 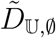 will be denoted 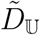.

Using these notations, the variance in selfing rate in the population can be written as:

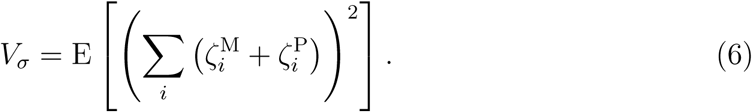

Ignoring genetic associations between different loci affecting the selfing rate, this becomes:

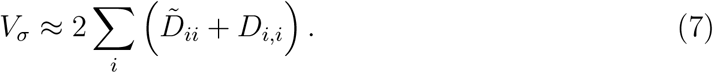

### General expression for fitness, and special cases

The fitness of an individual divided by the population mean fitness 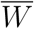 can be expressed in terms of “selection coefficients” *a*_𝕌,𝕍_ representing the effect of selection acting on the sets 𝕌 and 𝕍 of loci (Barton and Turelli, 1991; Kirkpatrick et al., 2002):

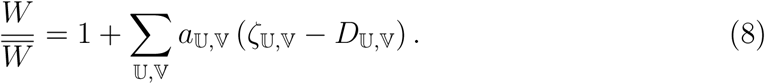

Throughout the paper, we assume no effect of the sex-of-origin of genes on fitness, so that *a*_𝕌,𝕍_ = *a*_𝕍,𝕌_. The coefficient *a*_*j*,ø_ = *a*_ø,*j*_ will be denoted *a*_*j*_ and represents selection for allele 1 at locus *j*. The coefficient *a*_*j,j*_ represents the effect of dominance at locus *j*, while *a*_*jk*,ø_ and *a*_*j,k*_ represent cis and trans epistasis between loci *j* and *k*. Coefficients *a*_*jk,j*_ and *a*_*jk,jk*_ respectively correspond to additive-by-dominance and dominance-by-dominance epistatic interactions between loci *j* and *k*, measured as deviations from additivity. Throughout the paper, we will assume that selection is weak, all *a*_𝕌,𝕍_ being of order *ϵ* (where *ϵ* is a small term), and derive general expressions for inbreeding depression and the strength of selection for selfing to leading order in *a*_𝕌,𝕍_ coefficients. Results for any particular fitness function can then be obtained by computing the corresponding expressions for *a*_𝕌,𝕍_ coefficients. We will consider three examples of fitness function that have been used in previous papers, and lead to different properties of the three forms of epistasis mentioned above. Approximate expressions for *a*_𝕌,𝕍_ coefficients under these fitness functions are computed in Supplementary File S1.

#### Uniformly deleterious alleles

Our first example corresponds to the case where allele 1 at each fitness locus *j* is deleterious, with selection and dominance coefficients *s* and *h*. Epistatic interactions occur between pairs of loci, and are decomposed into additive-by-additive (*e*_axa_), additive-by-dominance (*e*_axd_) and dominance-by-dominance (*e*_dxd_) epistasis (see Supplementary Figure S1 for an interpretation of these terms). We assume multiplicative effects of epistatic components on fitness *W* (*i.e.*, additive effects on log *W*), so that:

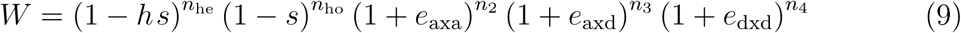

where *n*_he_ and *n*_ho_ are the numbers of loci at which a deleterious allele is present in the heterozygous (*n*_he_) or homozygous (*n*_ho_) state, while *n*_2_, *n*_3_ and *n*_4_ are the numbers of interactions between 2, 3 and 4 deleterious alleles at two different loci, given by:

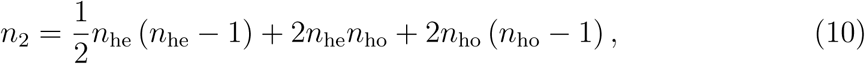

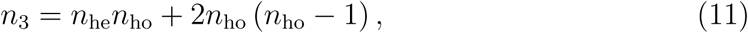

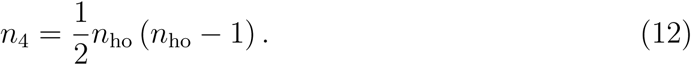

For example, *n*_2_ is given by the number of pairs of heterozygous loci in the genome (*n*_he_ (*n*_he_ − 1)/2), plus twice the number of pairs involving one heterozygous locus and one homozygous locus for the deleterious allele (*n*_he_*n*_ho_), plus four times the number of pairs of homozygous loci for the deleterious allele (*n*_ho_ (*n*_ho_ − 1)/2). In such models with fixed epistasis and possibly large numbers of loci, combinations of mutations quickly become advantageous when epistasis is positive, in which case they sweep through the population. We therefore focused on cases where *e*_axa_, *e*_axd_ and *e*_dxd_ are negative, and will assume throughout that deleterious alleles stay at low frequencies in the population (*p*_*j*_ remains small). As shown in Supplementary File S1, equation 9 leads to *a*_*jk*_ = *a*_*j,k*_ ≈ *e*_axa_, *a*_*jk,j*_ ≈ *e*_axd_ and *a*_*jk,jk*_ ≈ *e*_dxd_, while the strength of directional selection at each locus (*a*_*j*_) is affected by *e*_axa_ and the effective dominance (*a*_*j,j*_) is affected by *e*_axd_. Because epistatic coefficients are the same for all pairs of loci, equation 9 leads to a situation where the variances of *a*_*jk*_, *a*_*jk,j*_ and *a*_*jk,jk*_ over pairs of loci equal zero, while their mean values may depart from zero.

Charlesworth et al. (1991) explored the effect of synergistic epistasis (measured by a parameter *β*) on inbreeding depression, using a fitness function that imposes relations between *h, e*_axa_, *e*_axd_ and *e*_dxd_. As explained in Supplementary File S1, their fitness function (equation 2 in Charlesworth et al., 1991) is equivalent to setting *e*_axa_ = −*βh*^2^, *e*_axd_ = −*βh* (1 − 2*h*) and *e*_dxd_ = −*β* (1 − 2*h*)^2^ in our equation 9.

#### Gaussian stabilizing selection

Our second fitness function corresponds to stabilizing selection acting on an arbitrary number *n* of quantitative traits, with a symmetrical, Gaussian-shaped fitness function. The general model is the same as in Abu Awad and Roze (2018): *r*_*αj*_ denotes the effect of allele 1 at locus *j* on trait *α*, and we assume that the different loci have additive effects on traits:

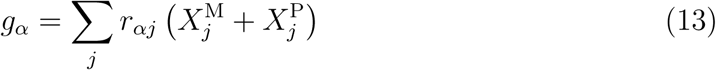

where *g*_*α*_ is the value of trait *α* in a given individual (note that *g*_*α*_ = 0 for all traits in an individual carrying allele 0 at all loci). We assume that the values of *r*_*αj*_ for all loci and traits are sampled from the same distribution with mean zero and variance *a*^2^. The fitness of individuals is given by:

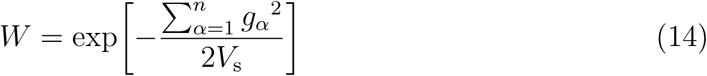

where *V*_s_ represents the strength of selection. According to equation 14, the optimal value of each trait is zero. We assume that 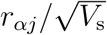 is small, so that selection is weak at each locus. This model generates distributions of fitness effects of mutations and of pairwise epistatic effects on fitness (the average value of epistasis being zero), while deleterious alleles have a dominance coefficient close to 1/4 in an optimal genotype (Martin and Lenormand, 2006b; Martin et al., 2007; Manna et al., 2011). In a population at equilibrium, equations 13 and 14 lead to 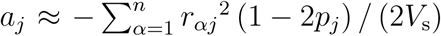 (*i.e.*, the rarer allele at locus *j* is disfavored), 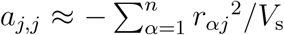 and 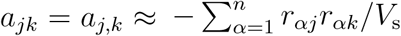, while *a*_*jk,j*_ and *a*_*jk,jk*_ are smaller in magnitude (see Supplementary File S1). This scenario thus generates a situation where additive-by-additive epistasis (*a*_*jk*_ = *a*_*j,k*_) is zero on average (because the average of *r*_*αj*_ is zero) but has a positive variance among pairs of loci, while additive-by-dominance and dominance-by-dominance epistasis are negligible. As in the previous example, we will generally assume that the deleterious allele at each locus *j* (allele 1 if *p*_*j*_ < 0.5, allele 0 if *p*_*j*_ > 0) stays rare in the population, by assuming that (1 − 2*p*_*j*_)^2^ is close to 1; this is also true in the next example.

#### Non-Gaussian stabilizing selection

The last example we examined is a generalization of the fitness function given by equation 14, in order to introduce a coefficient *Q* affecting the shape of the fitness peak (e.g., Martin and Lenormand, 2006a; Tenaillon et al., 2007; Gros et al., 2009; Roze and Blanckaert, 2014; Abu Awad and Roze, 2018):

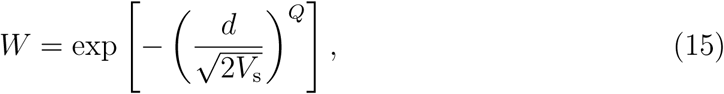

where 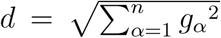 is the Euclidean distance from the optimum in phenotypic space. The fitness function is thus Gaussian when *Q* = 2, while *Q* > 2 leads to a flatter fitness peak around the optimum. The expressions for *a*_𝕌,𝕍_ coefficients derived in Supplementary File S1 show that the variances of *a*_*jk*_ = *a*_*j,k*_, *a*_*jk,j*_ and *a*_*jk,jk*_ over pairs of loci have the same order of magnitude, and that additive-by-additive epistasis (*a*_*jk*_ = *a*_*j,k*_) is zero on average, while additive-by-dominance and dominance-by-dominance epistasis (*a*_*jk,j*_, *a*_*jk,jk*_) are negative on average when *Q* > 2. Note that *Q* > 2 also generates higher-order epistatic interactions (involving more than two loci); however, we did not compute expressions for these terms.

### Quasi-linkage equilibrium (QLE) approximation

Using the general expression for fitness given by equation 8, the change in the mean selfing rate per generation can be expressed in terms of genetic associations between loci affecting the selfing rate and loci affecting fitness. Expressions for these associations can then be computed using general methods to derive recursions on allele frequencies and genetic associations (Barton and Turelli, 1991; Kirkpatrick et al., 2002). For this, we decompose the life cycle into two steps: selection corresponds to the differential contribution of individuals due to differences in overall fecundity and/or survival rates (*W*), while reproduction corresponds to gamete production and fertilization (involving either selfing or outcrossing). Associations measured after selection (that is, weighting each parent by its relative fitness) will be denoted 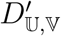, while associations after reproduction (among offspring) will be denoted 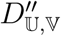. Assuming that “effective recombination rates” (that is, recombination rates multiplied by outcrossing rates) are sufficiently large relative to the strength of selection, genetic associations equilibrate rapidly relative to the change in allele frequencies due to selection. In that case, associations can be expressed in terms of allele frequencies by computing their values at equilibrium, for given allele frequencies (e.g., Barton and Turelli, 1991; Nagylaki, 1993). Note that when allele frequencies at fitness loci have reached an equilibrium (for example, at mutation-selection balance), one does not need to assume that the selection coefficients *a*_𝕌,𝕍_ are small relative to effective recombination rates for the QLE approximation to hold, but only that changes in allele frequencies due to the variation in the selfing rate between individuals are small. We will thus assume that the variance in the selfing rate in the population *V*_*σ*_ stays small (and therefore, the genetic variance contributed by each locus affecting the selfing rate is also small), and compute expressions to the first order in *V*_*σ*_. This is equivalent to the assumption that alleles at modifier loci have small effects, commonly done in modifier models.

### Individual-based simulations

In order to verify our analytical results, individual-based simulations were run using two C++ programs, one with uniformly deleterious alleles with fixed epistatic effects (equation 9) and the other with stabilizing selection on *n* quantitative traits (equation 14). Both are described in Supplementary File S5 and are available from Dryad. Both programs represent a population of *N* diploid individuals with discrete generations, the genome of each individual consisting of two copies of a linear chromosome with map length *R* Morgans. In the first program (fixed epistasis), deleterious alleles occur at rate *U* par haploid genome per generation at an infinite number of possible sites along the chromosome. A locus with an infinite number of possible alleles, located at the mid-point of the chromosome controls the selfing rate of the individual. In the program representing stabilizing selection, each chromosome carries *ℓ* equidistant biallelic loci affecting the *n* traits under selection (as in Abu Awad and Roze, 2018). The selfing rate is controlled by *ℓ*_*σ*_ = 10 additive loci evenly spaced over the chromosome, each with an infinite number of possible alleles (the selfing rate being set to zero if the sum of allelic values at these loci is negative, and one if the sum is larger than one). In both programs, mutations affecting the selfing rate occur at rate *U*_self_ = 10^−3^ per generation, the value of each mutant allele at a selfing modifier locus being drawn from a Gaussian distribution with standard deviation *σ*_self_ centered on the allele value before mutation. The selfing rate is set to zero during an initial burn-in period (set to 20,000 generations) after which mutations are introduced at selfing modifier loci.

## RESULTS

### Effects of epistasis on inbreeding depression

We first explore the effects of epistasis on inbreeding depression, assuming that the selfing rate is fixed. Throughout the paper, inbreeding depression *δ* is classically defined as:

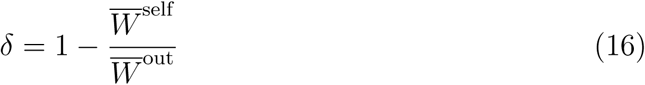

where 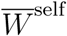 and 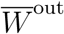 are the mean fitnesses of offspring produced by selfing and by outcrossing, respectively (e.g., Lande and Schemske, 1985). In Supplementary File S2, we show that a general expression for *δ* in terms of one- and two-locus selection coefficients, in a randomly mating population (*σ* = 0) is given by:

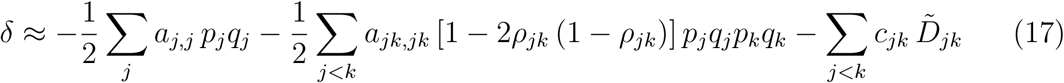

where the sums are over all loci affecting fitness, and with:

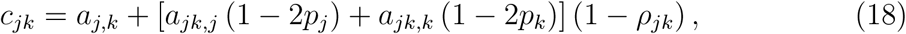

*ρ*_*jk*_ being the recombination rate between loci *j* and *k*. With arbitrary selfing, and assuming all *ρ*_*jk*_ ≈ 1/2, equation 17 generalizes to:

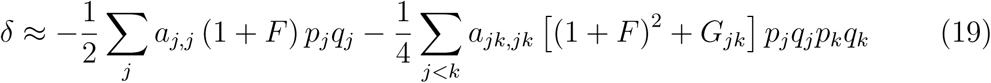

with several higher-order terms depending on genetic associations between loci generated by epistatic interactions (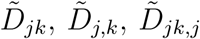, see equation B17 in Supplementary File S2 for the complete expression). The term *F* in equation 19 corresponds to the inbreeding coefficient (probability of identity by descent between the maternal and paternal copy of a gene), given by:

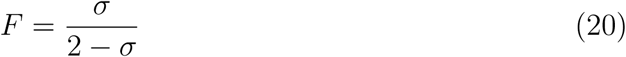

at equilibrium, while *G*_*jk*_ is the identity disequilibrium between loci *j* and *k* (Weir and Cockerham, 1973), given by:

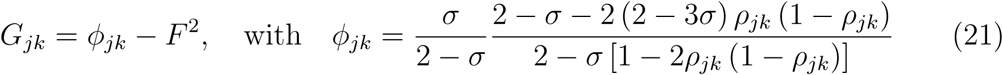

(*ϕ*_*jk*_ is the joint probability of identity by descent at loci *j* and *k*). Under free recombination (*ρ*_*jk*_ = 1/2), it simplifies to:

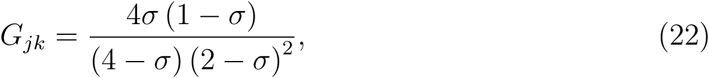

which will be denoted *G* hereafter. Given that *G*_*jk*_ is only weakly dependent on *ρ*_*jk*_, *G*_*jk*_ should be close to *G* for most pairs of loci when the genome map length is not too small.

#### Uniformly deleterious alleles

When fitness is given by equation 9, from equation 19 and using the expressions for the *a*_𝕌,𝕍_ coefficients given in Supplementary File S1 we find:

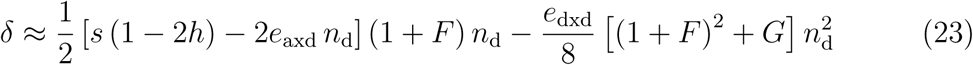

where *n*_d_ = ∑_*j*_ *p*_*j*_ is the average number of deleterious alleles per haploid genome. Equation 24 assumes that deleterious alleles stay rare in the population (so that terms in *p*_*j*_^2^ may be neglected). As explained in Supplementary File S2, the expression:

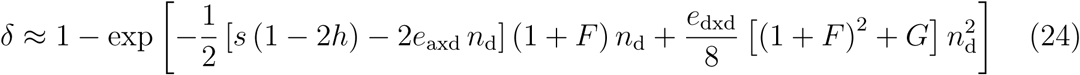

(obtained by assuming that the effects on inbreeding depression of individual loci and their interactions do multiply, rather than sum) provides more accurate results for parameter values leading to high inbreeding depression. Equation 24 yields the classical expression *δ* ≈ 1 − exp [−*U* (1 − 2*h*) / (2*h*)] in the absence of epistasis and under random mating (e.g., Charlesworth and Charlesworth, 1987).

Equations 23 and 24 only depend on the mean number of deleterious alleles *n*_d_ (and not on recombination rates between selected loci) because the effects of genetic associations between loci on *δ* have been neglected (as they generate higher-order terms, whose effects should remain small in most cases), and because *G*_*jk*_ was approximated by *G*. The equilibrium value of *n*_d_ can be obtained by solving

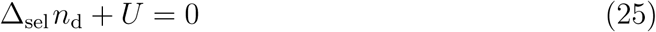

where Δ_sel_ *n*_d_ = _*j*_ Δ_sel_ *p*_*j*_ is the change in *n*_d_ due to selection and *U* is the deleterious mutation rate per haploid genome. From equation B26 in Supplementary File S2, we have to the first order in the selection coefficients (and assuming that deleterious alleles stay rare):

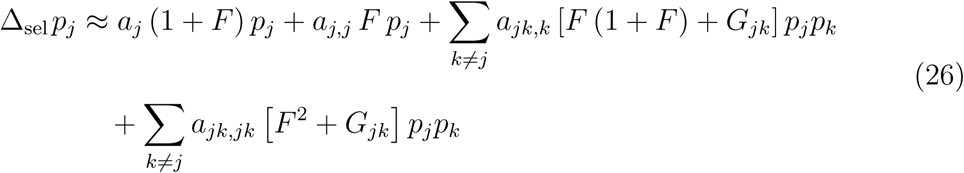

simplifying to *a*_*j*_ *p*_*j*_ under random mating. The first term of equation 26 represents the effect of directional selection against deleterious alleles (*a*_*j*_ < 0), which is increased by selfing due to the higher variance between individuals generated by homozygosity (by a factor 1 + *F*). The other terms represent additional effects of dominance and epistatic terms involving dominance arising when *σ* > 0 (that is, when the frequency of homozygous mutants is not negligible). Summing over loci and using the expressions for *a*_𝕌,𝕍_ coefficients given in Supplementary File S1, one obtains:

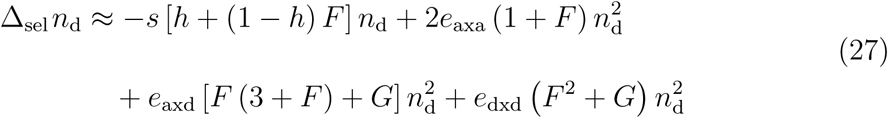

that can be used with equation 25 to obtain the equilibrium value of *n*_d_ (note that the term in *e*_axa_ in equation 27 stems from the term in *a*_*j*_ in equation 26, while part of the term in *e*_axd_ stems from the term in *a*_*j,j*_).

Equation 27 shows that, for non-random mating, negative values of *e*_axa_, *e*_axd_ or *e*_dxd_ reduce the mean number of deleterious alleles at equilibrium, thereby reducing inbreeding depression (the effects of *e*_axd_ and *e*_dxd_ on the equilibrium value of *n*_d_ disappear when mating is random, as *F* = *G* = 0 in this case). As shown by equation 24, negative values of *e*_axd_ and *e*_dxd_ also directly increase inbreeding depression (even under random mating), by decreasing the fitness of homozygous offspring. Figures 1A–C compare the predictions obtained from equations 24 and 27 with simulation results, testing the effect of each epistatic component separately. Negative *e*_axa_ reduces inbreeding depression by lowering the frequency of deleterious alleles in the population (equation 27, Figure 1A); furthermore, it reduces the purging effect of selfing, so that inbreeding depression may remain constant or even slightly increase as the selfing rate increases. When the selfing rate is low, *e*_axd_ and *e*_dxd_ have little effect on the mean number of deleterious alleles *n*_d_, and the main effect of negative *e*_axd_ and *e*_dxd_ is to increase inbreeding depression by decreasing the fitness of homozygous offspring (equation 24, Figures 1B–C). As selfing increases, this effect becomes compensated by the enhanced purging caused by negative *e*_axd_ and *e*_dxd_ (equation 27). Figure 1D shows the results obtained using Charlesworth et al.’s (1991) fitness function, yielding *e*_axa_ = −*βh*^2^, *e*_axd_ = −*βh* (1 − 2*h*) and *e*_dxd_ = −*β* (1 − 2*h*)^2^. Remarkably, the increased purging caused by negative epistasis almost exactly compensates the decreased fitness of homozygous offspring, so that inbreeding depression is only weakly affected by epistasis in this particular model; this result is also observed for different values of *s, U* and *h* (Supplementary Figures S2 – S4). The discrepancies between analytical and simulation results in Figure 1 likely stem from the effects of genetic associations, which are neglected in equations 24 and 27 (e.g., Roze, 2015). As shown by Supplementary Figures S2 – S4, these discrepancies become more important as the strength of epistasis increases (relative to *s*), as the mutation rate *U* increases and as dominance *h* decreases.

**Figure 1.**
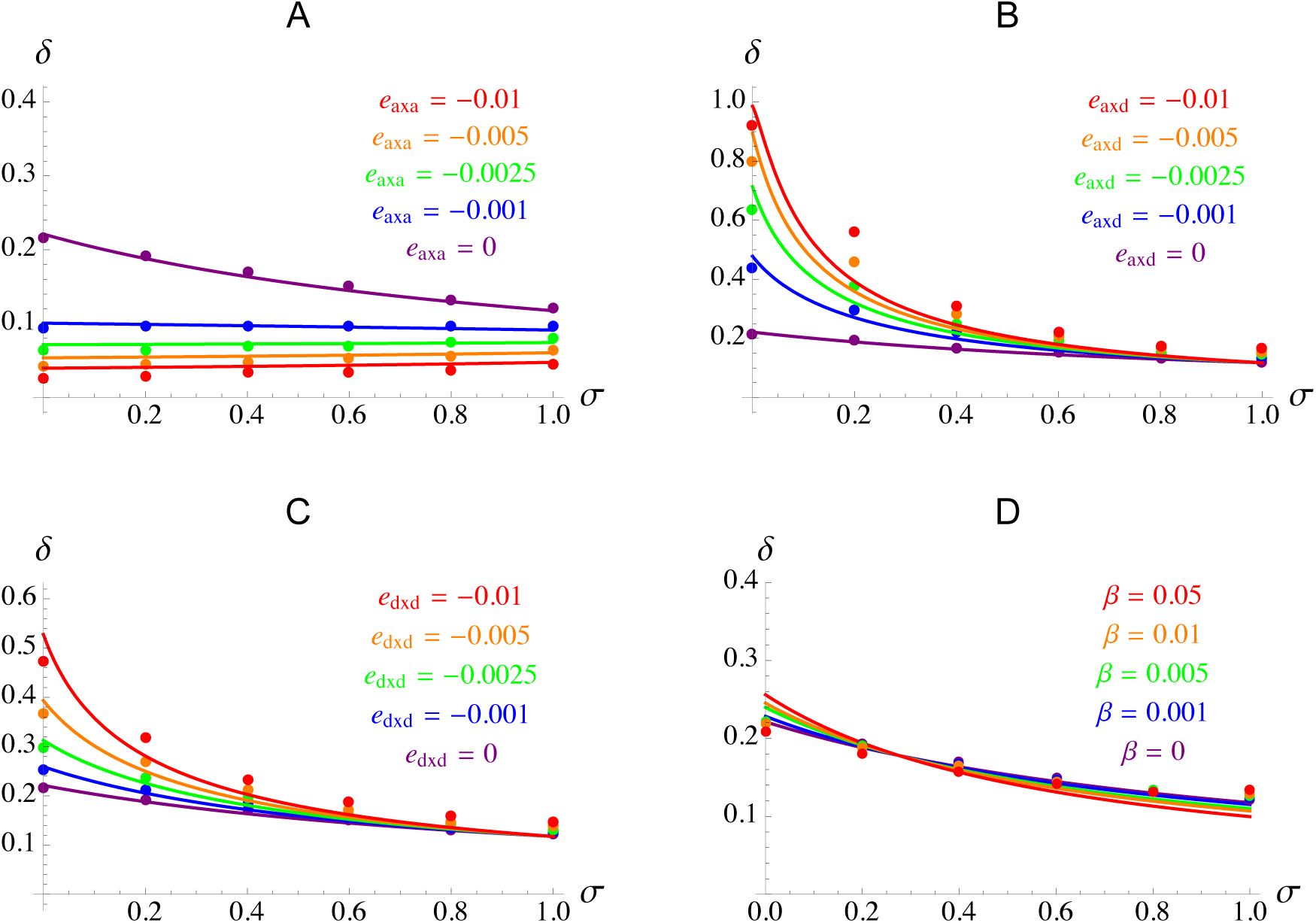
Inbreeding depression *δ* as a function of the selfing rate *σ*. A–C: effects of the different components of epistasis between deleterious alleles, additive-by-additive (*e*_axa_), additive-by-dominance (*e*_axd_) and dominance-by-dominance (*e*_dxd_) — in each plot, the other two components of epistasis are set to zero. D: results obtained using Charlesworth et al.’s (1991) fitness function, where *β* represents synergistic epistasis between deleterious alleles (slightly modified as explained in Supplementary File S1). Dots correspond to simulation results (error bars are smaller than the size of symbols), and curves to analytical predictions from equations 24 and 27. Parameter values: *U* = 0.25, *s* = 0.05, *h* = 0.25. In the simulations *N* = 20,000 (population size) and *R* = 10 (genome map length); simulations lasted 10^5^ generations and inbreeding depression was averaged over the last 5 × 10^4^ generations.

#### Gaussian stabilizing selection

An expression for inbreeding depression under Gaussian stabilizing selection (equation 14) is given in Abu Awad and Roze (2018). As shown in Supplementary File S2, this expression can be recovered from our general expression for *δ* in terms of *a*_𝕌,𝕍_ coefficients. Because the average epistasis is zero under Gaussian selection (e.g., Martin et al., 2007), inbreeding depression is only affected by the variance in epistasis, whose main effect is to generate linkage disequilibria that increase the frequency of deleterious alleles (see also Phillips et al., 2000) and thus increase *δ*. As shown by Abu Awad and Roze (2018), a different regime is entered above a threshold selfing rate when the mutation rate *U* is sufficiently large, in which epistatic interactions tend to lower inbreeding depression (see also Lande and Porcher, 2015).

#### Non-Gaussian stabilizing selection

Expressions for *a*_𝕌,𝕍_ coefficients under the more general fitness function given by equation 15 (Supplementary File S1) show that a “flatter-than-Gaussian”fitness peak (*Q* > 2) generates negative dominance-by-dominance epistasis (*a*_*jk,jk*_ < 0), increasing inbreeding depression (by contrast, the first term of equation 17 representing the effect of dominance is not affected by *Q*, as the effects of *Q* on *a*_*j,j*_ and on *p*_*j*_*q*_*j*_ cancel out). In the absence of selfing, and neglecting the effects of genetic associations among loci, one obtains (see Supplementary File S2 for derivation):

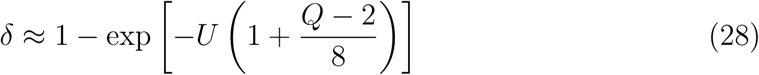

where the term in (*Q* − 2) /8 is generated by the term in *a*_*jk,jk*_ in equation 17. Although this expression differs from equation 29 in Abu Awad and Roze (2018) — that was obtained using a different method — both results are quantitatively very similar as long as *Q* is not too large (roughly, *Q* < 6). Generalizations of equation 28 to arbitrary *σ*, and including the effects of pairwise associations between loci (for *σ* = 0) are given in Supplementary File S2 (equations B40 and B54).

### Evolution of selfing in the absence of epistasis

Before exploring the effects of epistasis on selection for selfing, we first derive a general expression for the strength of indirect selection for selfing in the absence of epistasis (that is, ignoring the effects of coefficients *a*_*jk*_, *a*_*j,k*_, *a*_*jk,j*_ and *a*_*jk,jk*_ of the fitness function). In Supplementary File S3, we show that the change in the mean selfing rate 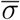 per generation can be decomposed into three terms:

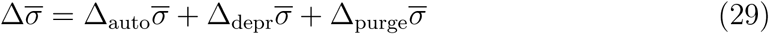

with:

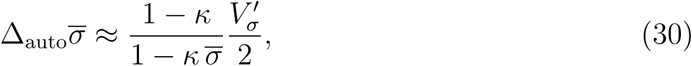

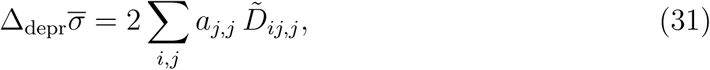

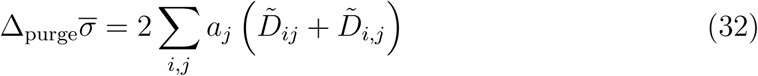

where the sums are over all loci *i* affecting the selfing rate and all loci *j* affecting fitness. The term 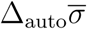 represents selection for increased selfing rates due to the automatic transmission advantage associated with selfing (Fisher, 1941). It is proportional to the variance in the selfing rate after selection 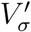, and vanishes when pollen discounting is complete (*κ* = 1). The second term corresponds to the effect of inbreeding depression. It depends on coefficients *a*_*j,j*_, representing the effect of dominance at loci affecting fitness; in particular, *a*_*j,j*_ < 0 when the average fitness of the two homozygotes at locus *j* is lower than the fitness of heterozygotes (which is the case when the deleterious allele at locus *j* is recessive or partially recessive). It also depends on associations 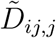 that are shown to be positive at QLE, reflecting the fact that alleles increasing the selfing rate tend to be present on more homozygous backgrounds. Finally, the last term depends on coefficients *a*_*j*_ representing directional selection for allele 1 at locus *j*, and associations 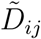 and 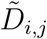 which are positive when alleles increasing the selfing rate at locus *i* tend to be associated with allele 1 at locus *j*, either on the same or on the other haplotype. This term is generally positive (favoring increased selfing rates), representing the fact that alleles coding for higher selfing increase the efficiency of selection at selected loci (by increasing homozygosity), and thus tend to be found on better purged genetic backgrounds, as explained in the Introduction (we show in Supplementary File S3 that 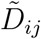 and 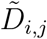 are also generated by other effects involving the identity disequilibrium between loci *i* and *j*, when 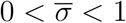).

The variance in the selfing rate after selection 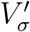, and the associations 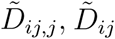 and 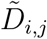 can be expressed in terms of *V*_*σ*_ and of allele frequencies using the QLE approximation described in the Methods. The derivations and expressions obtained for arbitrary values of 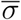 can be found in Supplementary File S3 (equations C31, C47, C48, C55 and C64), and generalize the results given by Epinat and Lenormand (2009) in the case of strong discounting (*κ* ≈ 1). When the mean selfing rate in the population approaches zero, one obtains:

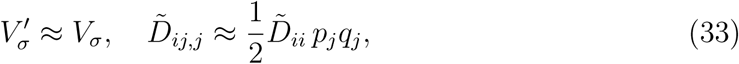

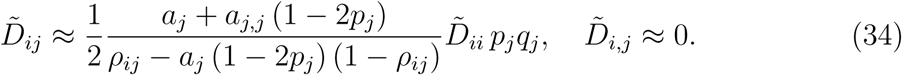

Using the fact that 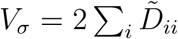 under random mating (equation 7), equations 30 – 34 yield, for 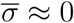:

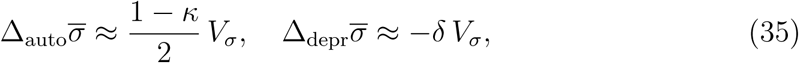

where *δ* = − (Σ_*j*_ *a*_*j,j*_ *p*_*j*_*q*_*j*_) /2 is inbreeding depression, neglecting the effect of interactions between selected loci (see equation 17), while

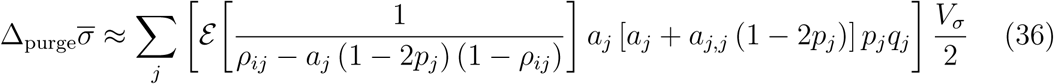

where the sum is over all loci *j* affecting fitness, and where *ε* is the average over all loci *i* affecting the selfing rate. Because 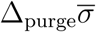 is of second order in the selection coefficients (*a*_*j*_, *a*_*j,j*_), it will generally be negligible relative to 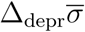 (which is of first order in *a*_*j,j*_), in which case selfing can increase if *δ* < (1 − *κ*) /2 (Charlesworth, 1980). When 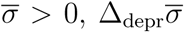 is not simply given by −*δ V*_*σ*_ (in particular, it also depends on the rate of pollen discounting and on identity disequilibria between loci affecting the selfing rate and loci affecting fitness, as shown by equation C31 in Supplementary File S3), but it is possible to show that 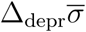 tends to decrease in magnitude as 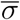 increases (while 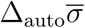 becomes stronger as 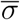 increases), leading to the prediction that 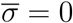 and 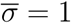 should be the only evolutionarily stable selfing rates (Lande and Schemske, 1985). As shown by equation 36, the relative importance of 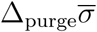 should increase when the strength of directional selection (*a*_*j*_) increases, when deviations from additivity (*a*_*j,j*_) are weaker and when linkage among loci is tighter.

In Supplementary File S3, we show that equation 36 can be expressed in terms of the increase in mean fitness caused by a single generation of selfing. In particular, if we imagine an experiment where a large pool of selfed offspring and a large pool of outcrossed offspring are produced from the same pool of parents (sampled from a randomly mating population), and if these offspring are allowed to reproduce (in proportion to their fitness, and by random mating within each pool), one can show that the mean fitness of the offspring of selfed individuals will be increased relative to the offspring of outcrossed individuals (due to purging), by an amount approximately equal to *P* = ∑_*j*_ *a*_*j*_ [*a*_*j*_ + *a*_*j,j*_ (1 − 2*p*_*j*_)] *p*_*j*_*q*_*j*_. Therefore, when linkage between loci affecting selfing and selected loci is not too tight (so that the term in *a*_*j*_ in the denominator of equation 36 may be neglected), 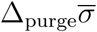 is approximately *P V*_*σ*_/ (2*ρ*_h_), where *ρ*_*h*_ is the harmonic mean recombination rate over all pairs of loci *i* and *j*, where *i* affects the selfing rate and *j* affects fitness (in the case of freely recombining loci, we thus have 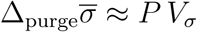).

#### Uniformly deleterious alleles

In the case where allele 1 at each fitness locus is deleterious with selection and dominance coefficients *s* and *h* (and assuming that *p*_*j*_ ≪ 1) we have *a*_*j*_ ≈ −*sh* and *a*_*j,j*_ ≈ −*s* (1 − 2*h*), while *p*_*j*_*q*_*j*_ ≈ *u/* (*sh*) at mutation-selection balance (where *u* is the mutation rate per locus). In that case, equation 36 simplifies to:

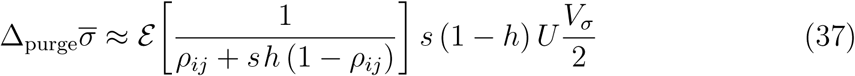

where *U* is the deleterious mutation rate per haploid genome and *ε* is now the average over all pairs of loci *i* and *j* (where locus *i* affects the selfing rate while locus *j* affects fitness). Figure 2A compares the predictions obtained from equations 35 and 37 with simulation results, in the absence of pollen discounting (*κ* = 0), and when alleles affecting the selfing rate have weak effects (*σ*_self_ = 0.01). Simulations confirm that selfing may evolve when inbreeding depression is higher than 0.5 (due to the effect of 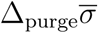), provided that the fitness effect of deleterious alleles is sufficiently strong. The prediction for the case of unlinked loci (obtained by setting *ρ*_*ij*_ = 0.5 in equation 37) actually gives a closer match to the simulation results than the result obtained by integrating equation 37 over the genetic map. This may stem from the fact that equation 37 overestimates the effect of tightly linked loci (possibly because the QLE approximation becomes inaccurate when *ρ*_*ij*_ is of the same order of magnitude as changes in allele frequencies due to selection for selfing). The effect of the size of mutational steps at the modifier locus does not affect the maximum value of inbreeding depression for which selfing can spread, as long as mutations tend to have small effects on the selfing rate (compare Figure 2A and 2B). However, the relative effect of purging (observed for high values of *s*) becomes more important when selfing can evolve by mutations of large size (*σ*_self_ = 0.3 in Figure 2C, while mutations directly lead to fully selfing individuals in Figure 2D), in agreement with the results obtained by Charlesworth et al. (1990) — note that our approximations break down when selfing evolves by large-effect mutations.

**Figure 2.**
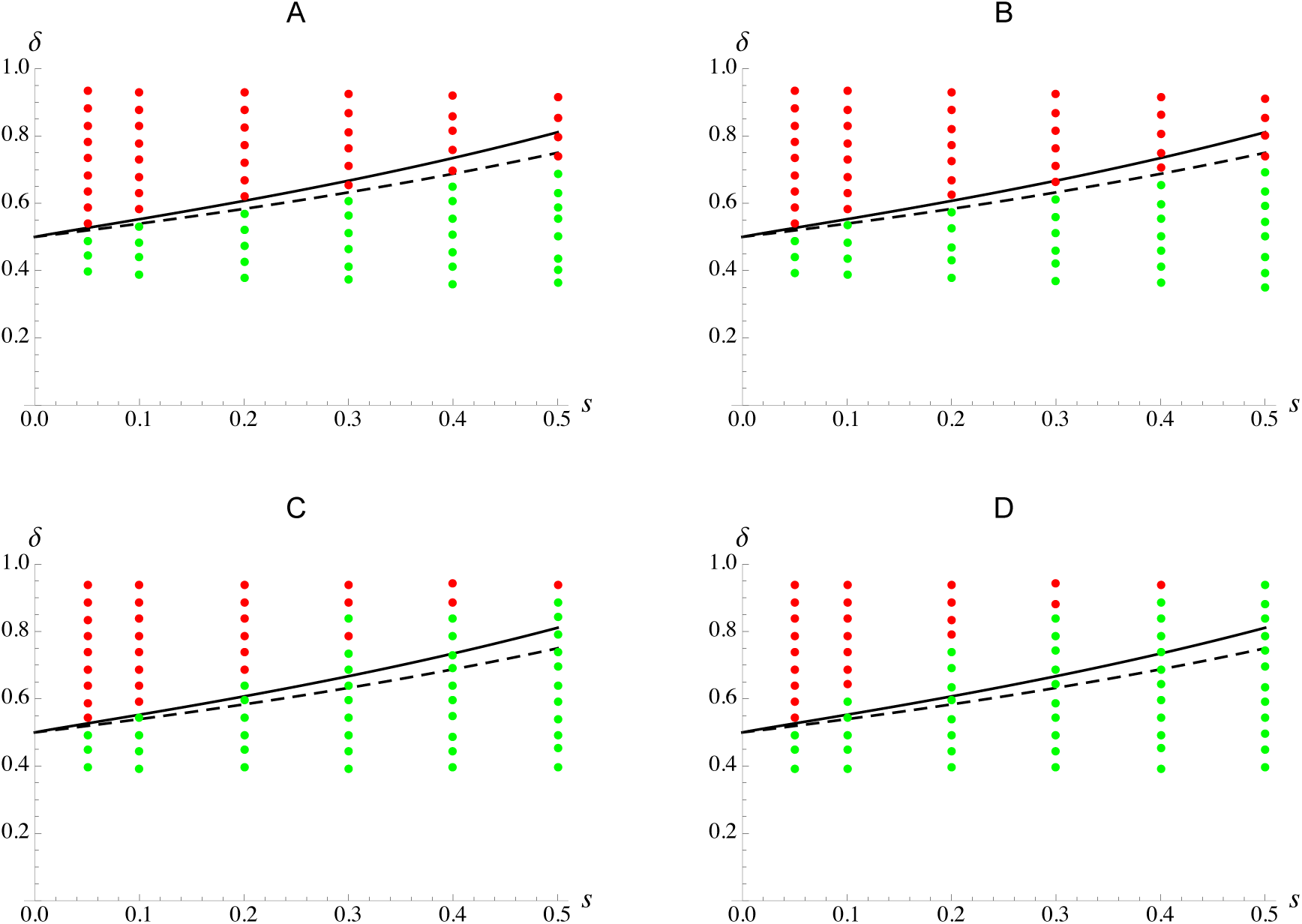
Evolution of selfing in the absence of epistasis. The solid curve shows the maximum value of inbreeding depression *δ* for selfing to spread in an initially outcrossing population, as a function of the strength of selection *s* against deleterious alleles (obtained from equations 35 and 37, after integrating equation 37 over the genetic map), while the dashed curve corresponds to the same prediction in the case of unlinked loci (obtained by setting *ρ*_*ij*_ = 1/2 in equation 37). Dots correspond to simulation results (using different values of *U* for each value of *s*, in order to generate a range of values of *δ*). In the simulations the population evolves under random mating during the first 20,000 generations (inbreeding depression is estimated by averaging over the last 10,000 generations); mutation is then introduced at the selfing modifier locus. A red dot means that the selfing rate stayed below 0.05 during the 2 × 10^5^generations of the simulation, while a green dot means that selfing increased (in which case the population always evolved towards nearly complete selfing). Parameter values: *κ* = 0, *h* = 0.25, *R* = 10; in the simulations *N* = 20,000, *U*_self_ = 0.001 (mutation rate at the selfing modifier locus). In A, the standard deviation of mutational effects at the modifier locus is set to *σ*_self_ = 0.01, while it is set to *σ*_self_ = 0.03 in B, and to *σ*_self_ = 0.3 in C. In D, only two alleles are possible at the modifier locus, coding for *σ* = 0 or 1, respectively.

#### Gaussian and non-Gaussian stabilizing selection

In the case of multivariate Gaussian stabilizing selection acting on *n* traits coded by biallelic loci with additive effects (equation 14) we have (to the first order in the strength of selection 1*/V*_s_): *a*_*j*_ = −*ς*_*j*_ (1 − 2*p*_*j*_) and *a*_*j,j*_ = −2*ς*_*j*_, where 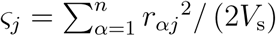 is the fitness effect of a heterozygous mutation at locus *j* in an optimal genotype. Assuming that polymorphism stays weak at loci coding for the traits under stabilizing selection, so that (1 − 2*p*_*j*_)^2^ ≈ 1, and using the fact that *p*_*j*_*q*_*j*_ ≈ *u/ς*_*j*_ under random mating (from equation B26, and neglecting interactions between loci), one obtains from equation 36:

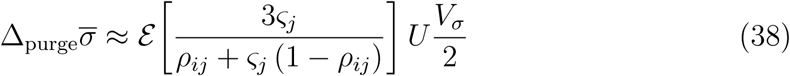

which is equivalent to equation 37 when introducing differences in *s* among loci, with *h* = 1/4 (note that the homozygous effect of mutations at locus *j* in an optimal genotype is ≈ 4*ς*_*j*_). When neglecting the term in *ς*_*j*_ in the denominator of equation 38, this simplifies to:

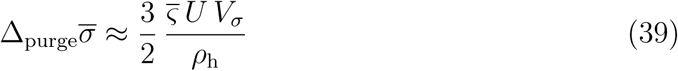

where 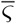 is the average heterozygous effect of mutations on fitness in an optimal genotype, and where *ρ*_h_ is the harmonic mean recombination rate over all pairs of loci *i* and *j*, where *i* affects the selfing rate and *j* affects the traits under stabilizing selection. Using the fitness function given by equation 15 (where *Q* describes the shape of the fitness peak), equation 39 generalizes to:

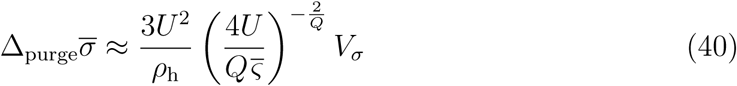

(see Supplementary File S1), which increases as *Q* increases in most cases (the derivative of equation 40 with respect to *Q* is positive as long as 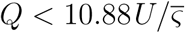). Therefore, for a given value of inbreeding depression and a fixed 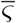, a flatter fitness peak tends to increase the relative importance of purging on the spread of selfing mutants in an outcrossing population.

### Effects of epistasis on the evolution of selfing

We now extend the previous expressions to include the effect of epistasis between pairs of selected loci. For this, we assume that all selection coefficients *a*_𝕌,𝕍_ are of the same order of magnitude (of order *ϵ*), and derive expressions for the effects of epistatic coefficients *a*_*jk*_, *a*_*j,k*_, *a*_*jk,j*_ and *a*_*jk,jk*_ on the change in mean selfing rate 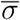 to leading order in *ϵ*. Because the expressions quickly become cumbersome under partial selfing, we restrict our analysis to the initial spread of selfing in an outcrossing population 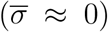. Figure 3 summarizes the different effects of epistasis, that are detailed below.

**Figure 3.**
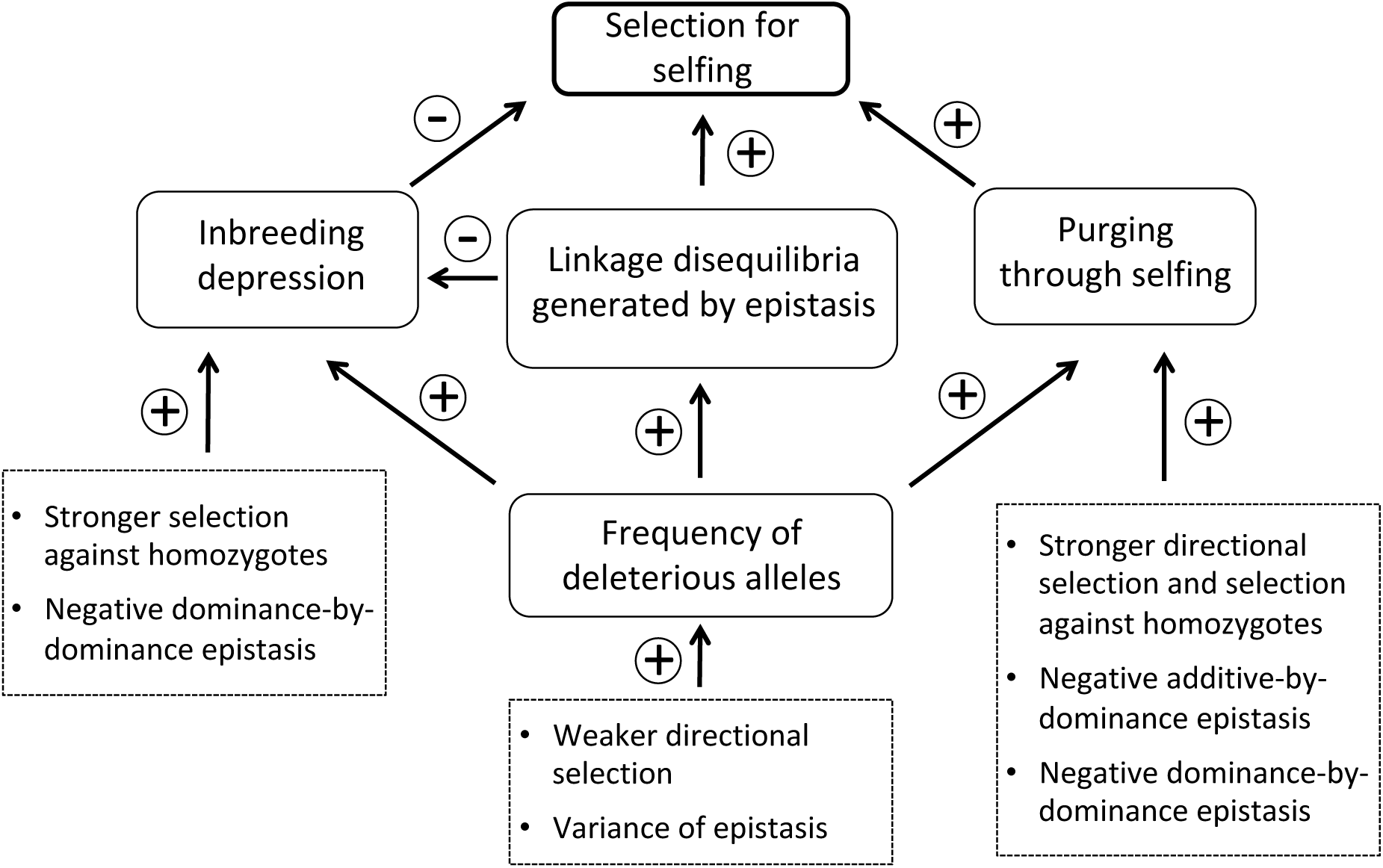
Summary of the effects of the strength of directional selection (*a*_*j*_), the strength of selection against homozygotes (*a*_*j,j*_) and epistasis (*a*_*jk*_, *a*_*j,k*_, *a*_*jk,j*_, *a*_*jk,jk*_) on the three components of indirect selection for selfing, in a randomly mating population. Note that *a*_*j*_ and *a*_*j,j*_ may be affected by epistatic interactions among loci (e.g., equations A10, A11 in Supplementary File S1).

As shown in Supplementary File S4, the change in mean selfing rate per generation now writes:

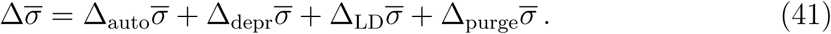

As above, 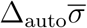 represents the direct transmission advantage of selfing and is still given by equation 35 as 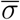 tends to zero. The term 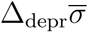 corresponds to the effect of inbreeding depression; taking into account epistasis between selected loci, it writes:

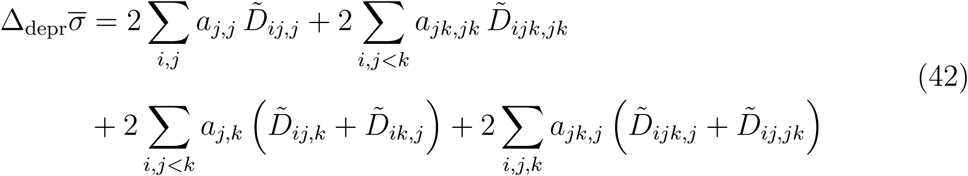

As shown in Supplementary File S4, expressing the different associations that appear in equation 42 at QLE, to leading order (and when 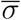 tends to zero) yields 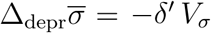, where *δ*′ is inbreeding depression measured after selection, that is, when the parents used to produced selfed and outcrossed offspring contribute in proportion to their fitness (an expression for *δ*′ in terms of allele frequencies and associations between pairs of loci is given by equation B9 in Supplementary File S2). Indeed, what matters for the spread of selfing is the ratio between the mean fitnesses of selfed and outcrossed offspring, taking into account the differential contributions of parents due to their different fitnesses. With epistasis, inbreeding depression is affected by genetic associations between selected loci, and *δ*′ thus depends on the magnitude of those associations after selection. Note that epistasis may also affect inbreeding depression through the effective dominance *a*_*j,j*_ and the equilibrium frequency *p*_*j*_ of deleterious alleles (as described earlier), and these effects are often stronger than effects involving genetic associations when epistasis differs from zero on average.

The new term 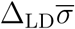 appearing in equation 41 represents an additional effect of epistasis (besides its effects on inbreeding depression *δ*′), and is given by:

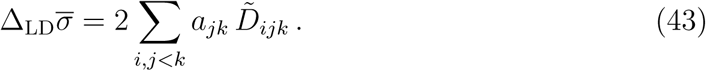

The association 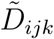 represents the fact that the linkage disequilibrium *D*_*jk*_ between loci *j* and *k* (generated by epistasis among those loci) tends to be stronger on haplotypes that also carry an allele increasing the selfing rate at locus *i*. Indeed, the magnitude of *D*_*jk*_ depends on the relative forces of selection generating *D*_*jk*_ and recombination breaking it, and selfing affects both processes: by increasing homozygosity, selfing reduces the effect of recombination (e.g., Nordborg, 1997), but it also increases “effective” epistasis, given that when a beneficial combination of alleles is present on one haplotype of an individual, it also tends to be present on the other haplotype due to homozygosity, enhancing the effect of fitness differences between haplotypes. An expression for 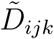 at QLE is given in Supplementary File S4, showing that 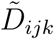 is generated by all epistatic components (*a*_*jk*_, *a*_*j,k*_, *a*_*jk,j*_, and *a*_*jk,jk*_). However, in the case of stabilizing selection the terms in *a*_*jk,j*_, and *a*_*jk,jk*_ should vanish when summed over sufficiently large numbers of loci (as explained in Supplementary File S4).

As in the previous section, the term 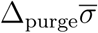 equals 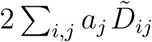 under random mating and represents indirect selection for selfing due to the fact that selfing increases the efficiency of selection against deleterious alleles. At QLE and to the first order in *a*_𝕌,𝕍_ coefficients, the linkage disequilibrium 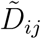 is given by (see Supplementary File S4 for derivation):

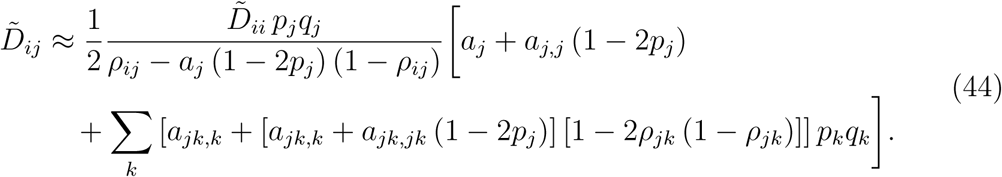

The term on the first line of equation 44 is the same as in equation 34, representing the fact that increased homozygosity at locus *j* improves the efficiency of selection acting at this locus. Note that epistatic interactions may affect this term (in particular when the average epistasis between selected loci differs from zero) through the selection coefficients *a*_*j*_ and *a*_*j,j*_ as well as equilibrium allele frequencies *p*_*j*_. The term on the second line of equation 44 shows that negative additive-by-dominance (*a*_*jk,j*_) or dominance-by-dominance epistasis (*a*_*jk,jk*_) between deleterious alleles increase the benefit of selfing: indeed, negative values of *a*_*jk,j*_ and *a*_*jk,jk*_ increase the magnitude of the negative linkage disequilibrium between alleles increasing selfing and disfavored alleles at loci affecting fitness (allele 1 in the case of uniformly deleterious alleles, or the rarer allele in the case of stabilizing selection). This effect stems from the increased homozygosity of offspring produced by selfing, negative values of *a*_*jk,j*_ and *a*_*jk,jk*_ increasing the efficiency of selection against deleterious alleles in homozygous individuals.

#### Uniformly deleterious alleles

Under fixed selection and epistatic coefficients across loci (fitness given by equation 9) and assuming that deleterious alleles stay rare in the population, one obtains for 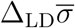:

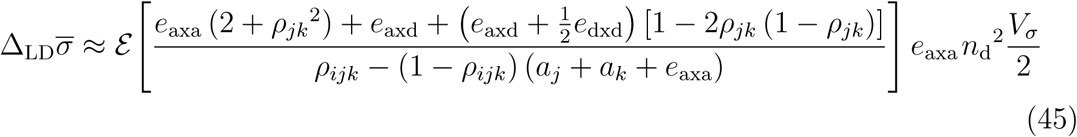

where *ε* is the average over all triplets of loci *i, j* and *k, ρ*_*ijk*_ is the probability that at least one recombination event occurs between the three loci *i, j* and *k* during meiosis (note that the denominator is approximately *ρ*_*ijk*_ when recombination rates are large relative to selection coefficients), and where *n*_d_ is the mean number of deleterious alleles per haploid genome. Assuming free recombination among all loci (*ρ*_*jk*_ = 1/2, *ρ*_*ijk*_ = 3/4), equation 45 simplifies to:

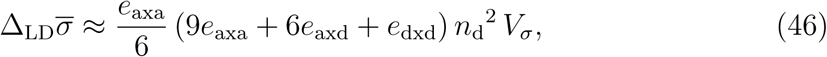

or, using Charlesworth et al.’s (1991) fitness function:

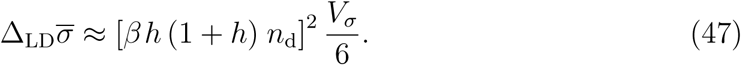

Furthermore, 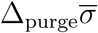 is given by:

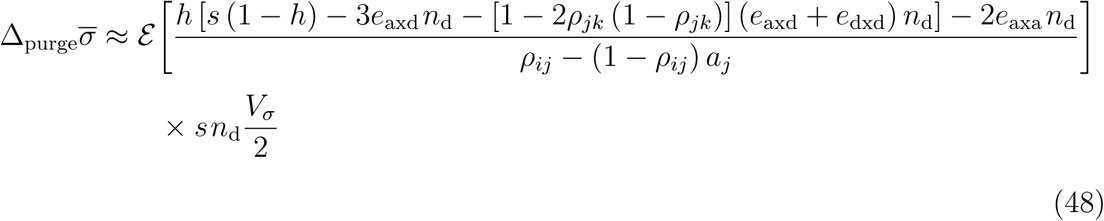

simplifying to:

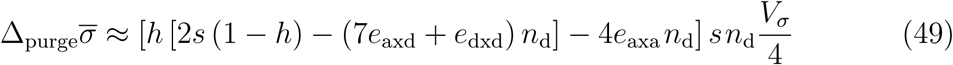

under free recombination.

Figure 4 shows the parameter space (in the *κ* – *δ*′ plane) in which an initially outcrossing population 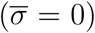 evolves towards selfing, in the case of uniformly deleterious alleles. Note that when selfing increased in the simulations (green dots), we always observed that the population evolved towards selfing rates close to 1. Figures 4A–C show that negative *e*_axd_ or *e*_dxd_ (the other epistatic components being set to zero) slightly increase the parameter range under which selfing evolves: in particular, selfing can invade for values of inbreeding depression *δ*′ slightly higher than 0.5 in the absence of pollen discounting (*κ* = 0). Epistasis has stronger effects when negative *e*_axd_ and/or *e*_dxd_ are combined with negative *e*_axa_, as shown by Figures 4D–F (we did not test the effect of negative *e*_axa_ alone, as *δ*′ is greatly reduced in this case unless *e*_axa_ is extremely weak). The QLE model (dashed and solid curves) correctly predicts the maximum inbreeding depression *δ*′ for selfing to evolve, as long as this maximum is not too large: high values of *δ*′ indeed imply high values of *U*, for which the QLE model overestimates the strength of indirect effects (in particular, the model predicts that selfing may evolve under high depression, above the upper parts of the curves in Figures 4D–F, but this was never observed in the simulations). This discrepancy may stem from higher-order associations between selected loci (associations involving 3 or more selected loci), that are neglected in this analysis and may become important when large numbers of mutations are segregating.

**Figure 4.**
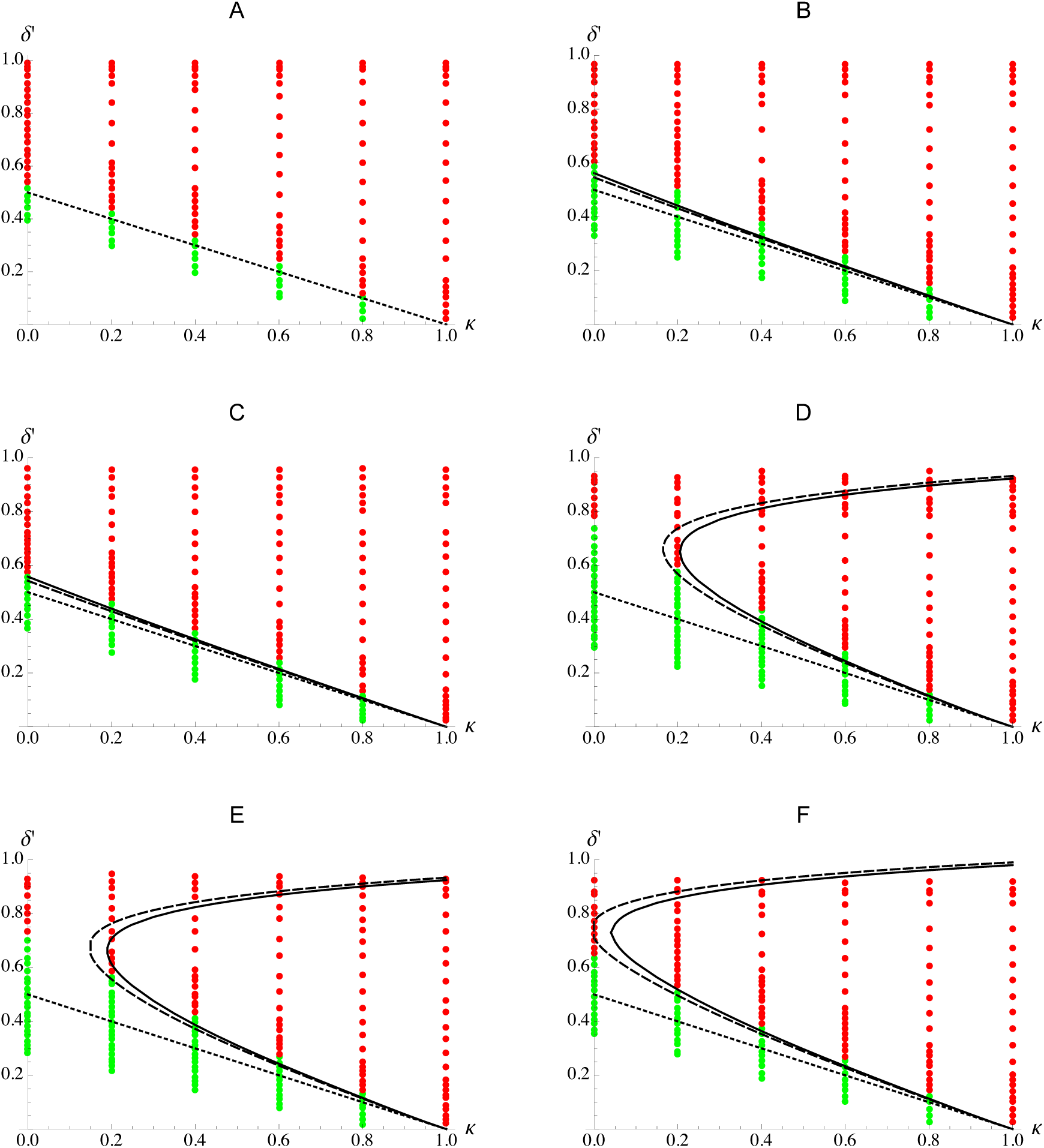
Evolution of selfing with fixed, negative epistasis. The different plots show the maximum value of inbreeding depression *δ*′ (measured after selection) for selfing to spread in an initially outcrossing population, as a function of the rate of pollen discounting *κ*. Green and red dots correspond to simulation results and have the same meaning as in Figure 2 (*δ*′ was estimated by averaging over the last 10,000 generations of the 20,000 preliminary generations without selfing, simulations lasted 2×10^5^ generations). The dotted lines correspond to the predicted maximum inbreeding depression for selfing to increase obtained when neglecting 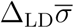 and 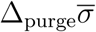 (that is, *δ*′ = (1 − *κ*) /2), the dashed curves correspond to the prediction obtained using the expressions for 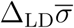 and 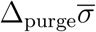 under free recombination (equations 46 and 49), while the solid curves correspond to the predictions obtained by integrating equations 45 and 48 over the genetic map (the effect of 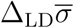 is predicted to be negligible relative to the effect of 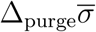 in all cases). To obtain these predictions, the relation between the mean number of deleterious alleles per haplotype *n*_d_ (that appears in equations 45–46 and 48–49) and *δ*′ was obtained from a fit of the simulation results. A: *e*_axa_ = *e*_axd_ = *e*_dxd_ = 0; B: *e*_axa_ = *e*_dxd_ = 0, *e*_axd_ = −0.01; C: *e*_axa_ = *e*_axd_ = 0, *e*_dxd_ = −0.01; D: *e*_axa_ = −0.005, *e*_axd_ = −0.01, *e*_dxd_ = 0; E: *e*_axa_ = −0.005, *e*_axd_ = *e*_dxd_ = −0.01; F: Charlesworth et al.’s (1991) model with *β* = 0.05. Other parameter values: *s* = 0.05, *h* = 0.25, *R* = 20; in the simulations *N* = 20,000, *U*_self_ = 0.001 (mutation rate at the selfing modifier locus), *σ*_self_ = 0.03 (standard deviation of mutational effects at the modifier locus).

In all cases shown in Figure 4, the increased parameter range under which selfing can evolve is predicted to be mostly due to the effect of negative epistasis on 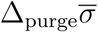, the effect of 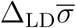 remaining negligible. Finally, one can note that the maximum *δ*′ for selfing to evolve is lower with *e*_axa_ = −0.005, *e*_axd_ = *e*_dxd_ = −0.01 (Figure 4E) than with *e*_axa_ = −0.005, *e*_axd_ = −0.01, *e*_dxd_ = 0 (Figure 4D). This is due to the fact that negative *e*_axd_ and *e*_dxd_ have two opposite effects: they increase the effect of selection against homozygous mutations (which increases 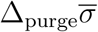), but they also increase the strength of inbreeding depression for a given mutation rate *U* (see Figure 1), decreasing the mean number of deleterious alleles per haplotype *n*_d_ associated with a given value of *δ*′ (which decreases 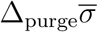).

Supplementary Figure S5 shows the effect of the size of mutational steps at the selfing modifier locus, in the absence of epistasis (corresponding to Figure 4A), and with all three components of epistasis being negative (corresponding to Figure 4E). Increasing the size of mutational steps has more effect in the presence of negative epistasis, since negative epistasis increases the purging advantage of alleles coding for more selfing 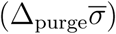, whose effect becomes stronger relative to 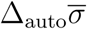 and 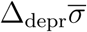 when modifier alleles have larger effects (as previously shown in Figure 2).

#### Gaussian and non-Gaussian stabilizing selection

Under stabilizing selection acting on quantitative traits (and assuming that recombination rates are not too small), one obtains:

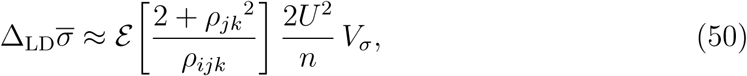

(where *n* is the number of selected traits) independently of the shape of the fitness peak *Q*, simplifying to (6*U*^2^*/n*) *V*_*σ*_ under free recombination (see Supplementary File S4). Independence from *Q* stems from the fact that 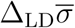 is proportional to ∑_*j,k*_ *a*_*jk*_^2^ *p*_*j*_*q*_*j*_*p*_*k*_*q*_*k*_, while *Q* has opposite effects on *a*_*jk*_^2^ and on *p*_*j*_*q*_*j*_*p*_*k*_*q*_*k*_ (*a*_*jk*_^2^ decreases, while *p*_*j*_*q*_*j*_*p*_*k*_*q*_*k*_increases as *Q* increases), which compensate each other exactly in this sum.

Under Gaussian stabilizing selection (*Q* = 2), the coefficients *a*_*jk,j*_ and *a*_*jk,jk*_ are small relative to the other selection coefficients (as shown in Supplementary File S1), and their effect on 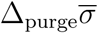 may thus be neglected (in which case 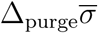 is still given by equation 39). With a flatter fitness peak (*Q* > 2), using the expressions for *a*_*jk,j*_ and *a*_*jk,jk*_ given by equations A54 and A55 in Supplementary File S1 yields:

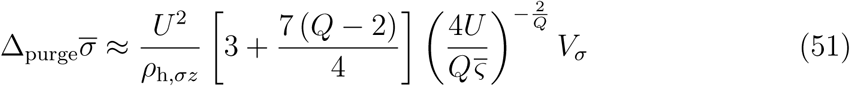

where the term in *Q*−2 between brackets corresponds to the term on the second line of equation 44 (effects of additive-by-dominance and dominance-by-dominance epistasis).

Figure 5 shows simulation results obtained under Gaussian stabilizing selection acting on different numbers of traits *n* (the mean deleterious effect of mutations 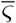 being kept constant by adjusting the variance of mutational effects *a*^2^). Under stabilizing selection, inbreeding depression reaches an upper limit as the mutation rate *U* increases (this upper limit being lower for smaller values of *n*), explaining why high values of *δ*′ could not be explored in Figure 5. Again, epistasis increases the parameter range under which selfing can invade (the effect of epistasis being stronger when the number of selected traits *n* is lower), and the QLE model yields correct predictions as long as inbreeding depression (and thus *U*) is not too large. In contrast with the previous example (uniformly deleterious alleles), the model predicts that 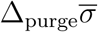 stays negligible, the difference between the dotted and solid/dashed curves in Figure 5 being mostly due to 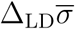: selfers thus benefit from the fact that they can maintain beneficial combinations of alleles (mutations with compensatory effects) at different loci. Interestingly, for *n* = 5 and sufficiently high rates of pollen discounting *κ*, selfing can invade if inbreeding depression is lower than a given threshold, or is very high. The latter case corresponds to a situation where polymorphism is important (high *U*) and where large numbers of compensatory combinations of alleles are possible. Although the model predicts that the same phenomenon should occur for higher values of *n*, it was not observed in simulations with *n* = 15 and *n* = 30, except for *n* = 15 and *κ* = 0.4. However, Supplementary Figures S6 and S7 show that the evolution of selfing above a threshold value of *δ*′ occurs more frequently when the fitness peak is flatter (*Q* > 2), and when mutations affecting the selfing rate have larger effects.

**Figure 5.**
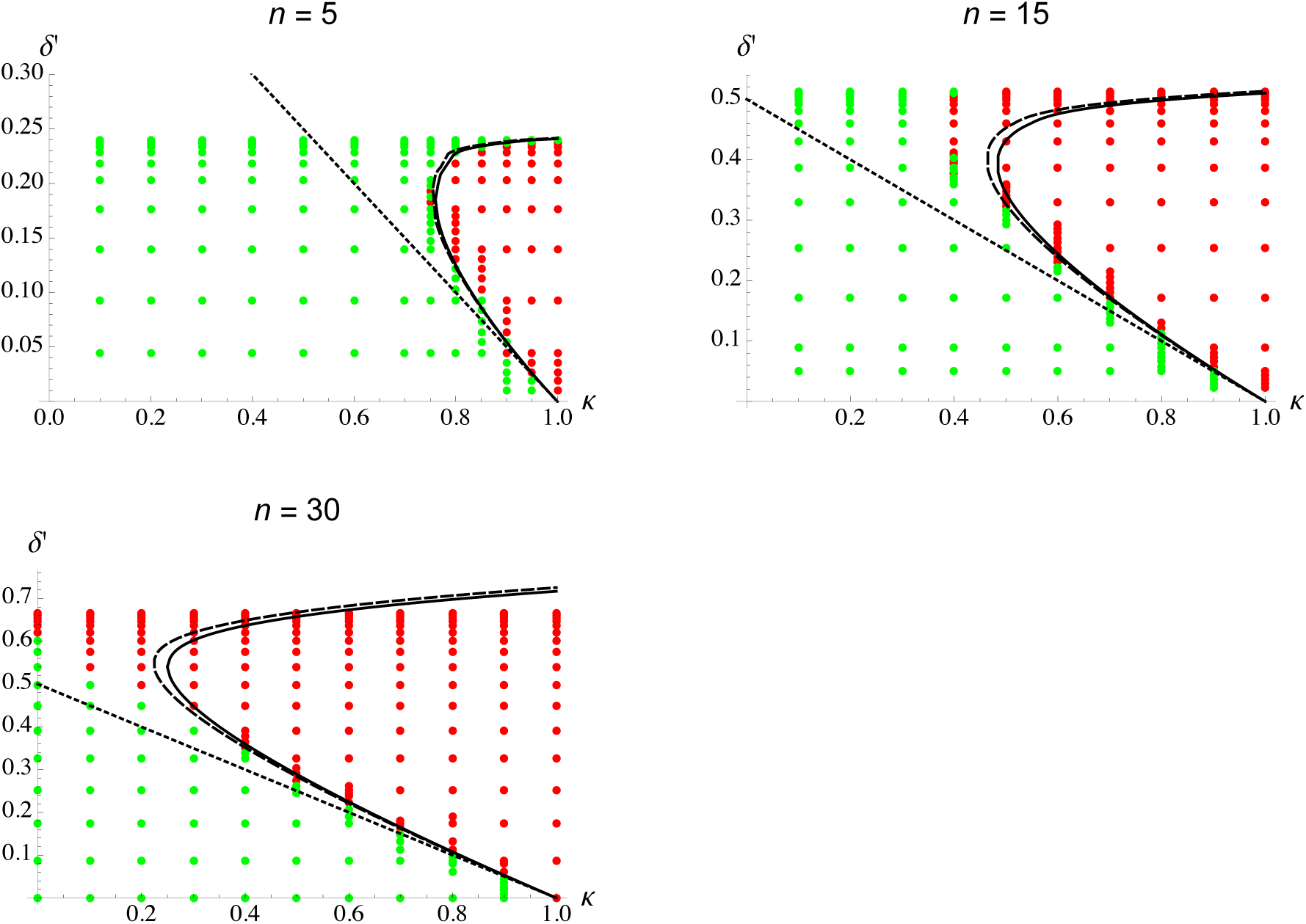
Evolution of self-fertilization under Gaussian stabilizing selection. The three plots show the effects of inbreeding depression *δ*′ (measured after selection) and pollen discounting (parameter *κ*) on the evolution of self-fertilization, for different numbers of traits under selection (*n* = 5, 15 and 30). Green and red dots correspond to simulation results and have the same meaning as in Figures 2 and 4 (*δ*′ was estimated by averaging over the last 10,000 generations of the 20,000 preliminary generations without selfing, simulations lasted 5 × 10^4^ generations). The fact that inbreeding depression reaches a plateau as *U* increases (at lower values of *δ*′ for lower values of *n*) sets an upper limit to the values of *δ*′ that can be obtained in the simulations. The dotted lines correspond to the predicted maximum inbreeding depression for selfing to increase obtained when neglecting 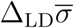 and 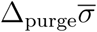 (that is, *δ*′ = (1 − *κ*) /2), the dashed curves correspond to the prediction obtained using the expression for 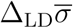 under free recombination (that is, 6*U* ^2^*V*_*σ*_*/n*, see equation 50), while the solid curves correspond to the predictions obtained by integrating equation 50 over the genetic map (the effect of 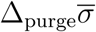 is predicted to be negligible relative to the effect of 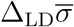). To obtain these predictions, the relation between *U* and *δ*′ was obtained from a fit of the simulation results. Other parameter values: 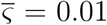, *R* = 20; in the simulations *N* = 5,000, *U*_self_ = 0.001 (overall mutation rate at selfing modifier loci), *σ*_self_ = 0.01 (standard deviation of mutational effects on selfing).

Finally, Figure 6 provides additional results on the effect of the number of selected traits *n*, for fixed values of the overall mutation rate *U*. Inbreeding depression is little affected by epistatic interactions when *n* is large, while low values of *n* tend to decrease inbreeding depression, explaining the shapes of the dotted curves showing the maximum level of pollen discounting for selfing to spread, when only taking into account the effects of the automatic advantage and inbreeding depression. The difference between the dotted and solid/dashed curves shows the additional effect of linkage disequilibria generated by epistasis 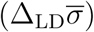, whose relative importance increases as the number of traits *n* decreases, and as the mutation rate *U* increases. Because *U* stays moderate (*U* = 0.2 or 0.5), the analytical model provides accurate predictions of the parameter range over which selfing is favored.

**Figure 6.**
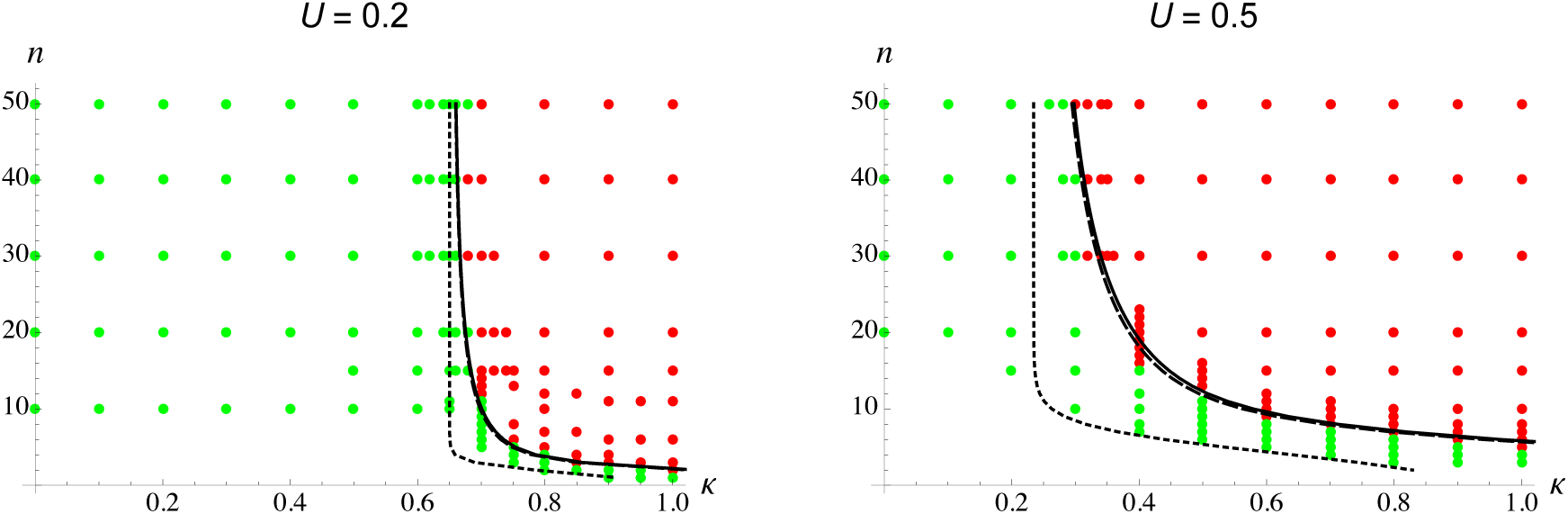
Evolution of self-fertilization under Gaussian stabilizing selection. The two plots show the effect of the number of traits under selection *n* and pollen discounting (parameter *κ*) on the evolution of self-fertilization for two values of the mutation rate on traits under stabilizing selection (*U* = 0.2 and 0.5). Green and red dots correspond to simulation results and have the same meaning as in the previous figures. The dotted curves show the maximum value of pollen discounting *κ* for selfing to increase obtained when neglecting 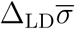 and 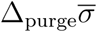 (that is, *δ*′ = (1 − *κ*) /2), while the dashed and solid curves correspond to the predictions including the term 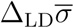 (from equation 50) under free recombination (dashed) or integrated over the genetic map (solid). To obtain these predictions, the relation between *n* and *δ*′ was obtained from a fit of the simulation results. Other parameter values: 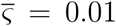, *R* = 20; in the simulations *N* = 5,000, *U*_self_ = 0.001 (overall mutation rate at selfing modifier loci), *σ*_self_ = 0.01 (standard deviation of mutational effects on selfing).

## DISCUSSION

The automatic transmission advantage associated with selfing and inbreeding depression are the two most commonly discussed genetic mechanisms affecting the evolution of self-fertilization. When these are the only forces at play, a selfing mutant arising in an outcrossing population is expected to increase in frequency as long as inbreeding depression is weaker than the automatic advantage, whose magnitude depends on the level of pollen discounting (Lande and Schemske, 1985; Holsinger et al., 1984). However, because selfers also tend to carry better purged genomes due to their increased homozygosity, several models showed that selfing mutants may invade under wider conditions than those predicted solely based on these two aforementioned forces (Charlesworth et al., 1990; Uyenoyama and Waller, 1991; Epinat and Lenormand, 2009; Porcher and Lande, 2005b; Gervais et al., 2014). Our analytical and simulation results confirm that the advantage procured through purging increases with the strength of selection against deleterious alleles and with the degree of linkage within the genome. The simulation results also indicate that the verbal prediction, according to which mutations causing complete selfing may invade a population independently of its level of inbreeding depression (Lande and Schemske, 1985, p. 33), only holds when deleterious alleles have strong fitness effects, so that purging occurs rapidly (Figure 2D).

Whether purging efficiency should significantly contribute to the spread of selfing mutants depends on the genetic architecture of inbreeding depression. To date, experimental data point to a small contribution of strongly deleterious alleles to inbreeding depression: for example, Baldwin and Schoen (2019) recently showed that in the self-incompatible species *Leavenworthia alabamica*, inbreeding depression is not affected by three generations of enforced selfing (which should have lead to the elimination of deleterious alleles with strong fitness effects). Previous experiments on different plant species also indicate that inbreeding depression is probably generated mostly by weakly deleterious alleles (Dudash et al., 1997; Willis, 1999; Carr and Dudash, 2003; Charlesworth and Willis, 2009). Data on the additive variance in fitness within populations are also informative regarding the possible effect of purging: indeed, using our general expression for fitness (equation 8) and neglecting linkage disequilibria, one can show that the additive component of the variance in fitness in a randomly mating population (more precisely, the variance in 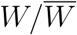) is given by the sum over selected loci of 2*a*_*j*_^2^*p*_*j*_*q*_*j*_ (see also eq. A3b in Charlesworth and Barton, 1996), a term which also appears in the effect of purging on the strength of selection for selfing (equation 36). Although estimates of the additive variance in fitness in wild populations remain scarce, the few estimates of the “evolvability” parameter (corresponding to the additive component of the variance in 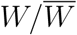) available from plant species are small, of the order of a few percent (Hendry et al., 2018). Note that strictly, the effect of purging on the strength of selection for selfing is proportional to the quantity ∑_*j*_ *a*_*j*_ [*a*_*j*_ + *a*_*j,j*_ (1 − 2*p*_*j*_)] *p*_*j*_*q*_*j*_ (equation 36), which may be larger than ∑ _*j*_ *a*_*j*_^2^*p*_*j*_*q*_*j*_(for example, in the case of deleterious alleles with fixed *s* and *h*, the first quantity is approximately *s* (1 − *h*) *U* and the second *shU*). As explained in the Results section (and in Supplementary File S3), the strength of selection for selfing through purging may, in principle, be estimated from the increase in mean fitness following a single generation of selfing. However, the small values of the available estimates of ∑_*j*_ *a*_*j*_^2^*p*_*j*_*q*_*j*_, together with the experimental evidence mentioned above on the genetics of inbreeding depression, indicate that selfing mutants probably do not benefit greatly from purging. Nevertheless, it remains possible that the strength of selection against deleterious alleles (*a*_*j*_) increases in harsher environments (Cheptou et al., 2000; Agrawal and Whitlock, 2010), leading to stronger purging effects in such environments.

The effects of epistasis between deleterious alleles on inbreeding depression and on the evolution of mating systems have been little explored (but see Charlesworth et al., 1991). In this paper, we derived general expressions for the effect of epistasis between pairs of loci on inbreeding depression and on the strength of selection for selfing, that can be applied to more specific models. Our results show that different components of epistasis have different effects on inbreeding depression: negative dominance-by-dominance epistasis directly increases inbreeding depression due to covariances in homozygosity across loci among selfed offspring, while additive-by-additive and additive-by-dominance epistasis may indirectly affect inbreeding depression by changing the effective strength of selection (*a*_*j*_) or effective dominance (*a*_*j,j*_) of deleterious alleles. Very little is known on the average sign and relative magnitude of these different forms of epistasis. In principle, the overall sign of dominance-by-dominance effects can be deduced from the shape of the relation between the inbreeding coefficient of individuals (*F*) and their fitness (Crow and Kimura, 1970, p. 80), an accelerating decline in fitness as *F* increases indicating negative *e*_dxd_. The relation between *F* and fitness-related traits was measured in several plant species; the results often showed little departure from linearity (e.g., Willis, 1993; Kelly, 2005), but the experimental protocols used may have generated biases against finding negative *e*_dxd_ (Falconer and Mackay, 1996; Lynch and Walsh, 1998; Sharp and Agrawal, 2016).

Most empirical distributions of epistasis between pairs of mutations affecting fitness have been obtained from viruses, bacteria and unicellular eukaryotes (e.g., Martin et al., 2007; Kouyos et al., 2007; de Visser and Elena, 2007). While no clear conclusion emerges regarding the average coefficient of epistasis (some studies find that it is negative, other positive and other close to zero), a general observation is that epistasis is quite variable across pairs of loci. This variance of epistasis may slightly increase inbreeding depression when it remains small (by reducing the efficiency of selection against deleterious alleles, Phillips et al., 2000; Abu Awad and Roze, 2018), or decrease inbreeding depression when it is larger and/or effective recombination is sufficiently weak, so that selfing can maintain beneficial multilocus genotypes (Lande and Porcher, 2015; Abu Awad and Roze, 2018). Besides this “short-term” effect on inbreeding depression, the variance of epistasis also favors selfing through the progressive buildup of linkage disequilibria that increase mean fitness (associations between alleles with compensatory effects at different loci): this effect is equivalent to selection for reduced recombination rates caused by the variance of epistasis among loci, previously described by Otto and Feldman (1997). Interestingly, this effect may become stronger than inbreeding depression above a threshold value of the rate of mutation on traits under stabilizing selection (Figures 4, S7). Is the variance of epistasis typically large enough, so that the benefit of maintaining beneficial combinations of alleles may significantly help selfing mutants to spread? Answering this question is difficult without better knowledge of the importance of epistatic interactions on fitness in natural environments. Nevertheless, some insights can be gained from our analytical results: for example, neglecting additive-by-dominance and dominance-by-dominance effects, equations 43 and D7 indicate that the effect of linkage disequilibria on the strength of selection for selfing should scale with the sum over pairs of selected loci of *a*_*jk*_^2^*p*_*j*_*q*_*j*_*p*_*k*_*q*_*k*_, which also corresponds to the epistatic component of the variance in fitness in randomly mating populations. Although estimates of epistatic components of variance remain scarce, they are typically not larger than additive components (e.g., Hill et al., 2008), suggesting that the benefit of maintaining beneficial multilocus genotypes may be generally limited (given that the additive variance in fitness seems typically small, as discussed previously).

Previous models on the evolution of recombination showed that increased recombination rates may be favored when epistasis is negative and sufficiently weak relative to the strength of directional selection (e.g., Barton, 1995). Similarly, weakly negative epistasis may favour the maintenance of outcrossing (through selection for recombination): this effect does not appear in our model, because it involves higher order terms (proportional to *a*_*j*_ *a*_*k*_ *a*_*jk*_) that were neglected in our analysis. When epistasis is weak (*a*_*jk*_ ≪ *a*_*j*_, *a*_*k*_), these terms may become of the same order as the term in *a*_*jk*_^2^ arising in 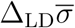, leading to a net effect of epistasis favoring outcrossing; however, the overall effect of epistasis should be negligible (relative to the effects of inbreeding depression and purging) when *a*_*jk*_ ≪ *a*_*j*_, *a*_*k*_. Selection for recombination (and outcrossing) due to negative epistasis may become stronger when effective rates of recombination between loci become smaller (in particular, in highly selfing populations), which may prevent evolution towards complete selfing. Indeed, Charlesworth et al. (1991) showed that in the presence of negative epistasis between deleterious alleles, and when outcrossing is not stable, a selfing rate slightly below one corresponds to the evolutionarily stable strategy (ESS). In finite populations, selection for recombination is also driven by the Hill-Robertson effect (through a term proportional to *a*_*j*_^2^*a*_*k*_^2^*/N*_e_, e.g., Barton and Otto, 2005) even in the absence of epistasis. Again, while this term should generally stay negligible (relative to inbreeding depression and purging) in an outcrossing population, it may become more important in highly selfing populations, due to their reduced effective size. Accordingly, Kamran-Disfani and Agrawal (2014) showed that selfing rates slightly below one are selectively favoured over complete selfing in finite populations under multiplicative selection (no epistasis). Similar effects must have occurred in our simulations, although we did not check that selfing rates slightly below one resulted from selection to maintain low rates of outcrossing, rather than from the constant input of mutations at selfing modifier loci (this could be done by comparing the probabilities of fixation of alleles coding for different selfing rates, as in Kamran-Disfani and Agrawal, 2014). Together with the results of Charlesworth et al. (1991) and Kamran-Disfani and Agrawal (2014), our simulation results indicate that, while selection for recombination may favour the maintenance of low rates of outcrossing in highly selfing populations, it cannot explain the maintenance of mixed mating systems (involving higher rates of outcrossing) under constant environmental conditions (the selfing rate always evolved towards values either close to zero or one in our simulations). It is possible that mixed mating systems may be more easily maintained under changing environmental conditions, however (for example, under directional selection acting on quantitative traits); this represents an interesting avenue for future research.

## Supporting information

Mathematica notebook

## Acknowledgements

We thank the bioinformatics and computing service of Roscoff’s Biological Station (Abims platform) for computing time, and Nick Barton, Sylvain Gandon and an anonymous reviewer for useful comments. This work was supported by the French Agence Nationale de la Recherche (project SEAD, ANR-13-ADAP-0011 and project SexChange, ANR-14-CE02-0001). Diala Abu Awad was partly funded by the TUM University Foundation Fellowship. Version 4 of this preprint has been peer-reviewed and recommended by Peer Community in Evolutionary Biology (https://doi.org/10.24072/pci.evolbiol.100093).

## Conflict of interest disclosure

The authors of this preprint declare that they have no financial conflict of interest with the content of this article.

Denis Roze is one of the PCI Evolutionary Biology recommenders.

## Data archiving

https://datadryad.org/stash/share/taHadrCrm9YXp0bmm21ILG4z5284tXziyAqVzXX-JrI

