## Supplementary File S1 for "Epistasis, inbreeding depression and the evolution of self-fertilization"

### FILE S1: DERIVATION OF SELECTION COEFFICIENTS

Taking only one and two-locus selection coefficients into account and assuming no sex-of-origin effect of genes on fitness, equation 8 in the main text writes:

$$\begin{aligned}\frac{W}{\overline{\overline{W}}} = 1 &+ \sum_j a_j \left(\zeta_{j,\emptyset} + \zeta_{\emptyset,j}\right) + \sum_j a_{j,j}\left(\zeta_{j,j} - D_{j,j}\right) \\ &+ \sum_{j<k} a_{jk}\left(\zeta_{jk,\emptyset} + \zeta_{\emptyset,jk} - 2\tilde{D}_{jk}\right) + \sum_{j<k} a_{j,k}\left(\zeta_{j,k} + \zeta_{k,j} - 2\tilde{D}_{j,k}\right) \\ &+ \sum_{j<k} a_{jk,j}\left(\zeta_{jk,j} + \zeta_{j,jk} - 2\tilde{D}_{jk,j}\right) + \sum_{j<k} a_{jk,k}\left(\zeta_{jk,k} + \zeta_{k,jk} - 2\tilde{D}_{jk,k}\right) \\ &+ \sum_{j<k} a_{jk,jk}\left(\zeta_{jk,jk} - D_{jk,jk}\right).\end{aligned} \tag{A1}$$

We derive here expressions for these different $a_{\mathbb{U},\mathbb{V}}$ coefficients under our three fitness functions: uniformly deleterious alleles with fixed epistasis, Gaussian stabilizing selection acting on $n$ quantitative traits, and stabilizing selection with non-Gaussian fitness function (equations 9, 14 and 15 in the main text, respectively).

**Uniformly deleterious alleles.** Expressions for $a_{\mathbb{U},\mathbb{V}}$ coefficients to the second order in $s$ and to the first order in epistatic coefficients ($e_{\text{axa}}$, $e_{\text{axd}}$, $e_{\text{dxd}}$) are given in Appendix A of Roze and Lenormand (2005), and can be obtained as follows. Assuming that $s$ is of order $\epsilon$ (where $\epsilon$ is a small term) and $e_{\text{axa}}$, $e_{\text{axd}}$, $e_{\text{dxd}}$ are of order $\epsilon^2$, equation 9 may be written as:

$$\begin{aligned}W = 1 - \sum_j S_j + \sum_{j<k}\Big[&S_j S_k + e_{\text{axa}}\left(X_j^{\text{M}} + X_j^{\text{P}}\right)\left(X_k^{\text{M}} + X_k^{\text{P}}\right) \\ &+ e_{\text{axd}}\left[\left(X_j^{\text{M}} + X_j^{\text{P}}\right)X_k^{\text{M}}X_k^{\text{P}} + \left(X_k^{\text{M}} + X_k^{\text{P}}\right)X_j^{\text{M}}X_j^{\text{P}}\right] \\ &+ e_{\text{dxd}}\, X_j^{\text{M}}X_j^{\text{P}}X_k^{\text{M}}X_k^{\text{P}}\Big] + o\!\left(\epsilon^2\right).\end{aligned} \tag{A2}$$

with

$$S_j = s\left[h\left(X_j^{\text{M}} + X_j^{\text{P}}\right) + (1-2h)\,X_j^{\text{M}}X_j^{\text{P}}\right] \tag{A3}$$

and where the last sum is over all pairs of loci $j$ and $k$ (counting each pair once). After replacing $X_j^{\mathrm{M}}$, $X_j^{\mathrm{P}}$ by $\zeta_j^{\mathrm{M}} + p_j$, $\zeta_j^{\mathrm{P}} + p_j$ (and similarly for$X_k^{\mathrm{M}}$, $X_k^{\mathrm{P}}$), expanding equation A2 and using equation 5 in the main text, one obtains an expression for $W$ in terms of $\zeta_{\mathbb{U},\mathbb{V}}$ variables, where $\mathbb{U}$ and $\mathbb{V}$ take values in $\{\emptyset, j, k, jk\}$. Mean fitness $\overline{W}$ is given by the same expression, replacing $\zeta_{\mathbb{U},\mathbb{V}}$ by $D_{\mathbb{U},\mathbb{V}}$. Expressing $W/\overline{W}$ then yields equation 8 in the main text, where $a_{\mathbb{U},\mathbb{V}}$ coefficients are given by (to the order $\epsilon^2$):

$$a_{j,\emptyset} = -s\left[h + p_j\left(1-2h\right)\right]\left(1+T_j\right) + \sum_k p_k\left[2e_{\mathrm{axa}} + e_{\mathrm{axd}}\left(2p_j + p_k\right) + e_{\mathrm{dxd}}\, p_j p_k\right] \quad \text{(A4)}$$

$$a_{j,j} = -s\left(1-2h\right)\left(1+T_j\right) + \sum_k p_k\left(2e_{\mathrm{axd}} + e_{\mathrm{dxd}}\, p_k\right) \quad \text{(A5)}$$

$$\begin{aligned} a_{jk,\emptyset} = a_{j,k} = {} & e_{\mathrm{axa}} + e_{\mathrm{axd}}\left(p_j + p_k\right) + e_{\mathrm{dxd}}\, p_j p_k \\ & + s^2\left[h + p_j\left(1-2h\right)\right]\left[h + p_k\left(1-2h\right)\right] \end{aligned} \quad \text{(A6)}$$

$$a_{jk,j} = e_{\mathrm{axd}} + e_{\mathrm{dxd}}\, p_k + s^2\left(1-2h\right)\left[h + p_k\left(1-2h\right)\right] \quad \text{(A7)}$$

$$a_{jk,jk} = e_{\mathrm{dxd}} + s^2\left(1-2h\right)^2 \quad \text{(A8)}$$

with:

$$T_j = s\, p_j\left[2h + p_j\left(1-2h\right)\right] + s\left(1-2h\right)\left(D_{j,j} + \sum_k D_{k,k}\right) \quad \text{(A9)}$$

and where the sums are over all loci $k$ different from $j$. Selection acting at a locus will only have an effect when this locus is polymorphic, and $a_{\mathbb{U},\mathbb{V}}$ coefficients with a '$j$' index will thus be multiplied by $p_j q_j$ in expressions describing changes in allele frequencies and genetic associations due to selection. Assuming that deleterious alleles stay at low frequency, we may neglect terms in ${p_j}^2$, and thus neglect the terms in $p_j$ appearing in expressions for $a_{\mathbb{U},\mathbb{V}}$ coefficients with a '$j$' index. Furthermore, one can show that $D_{j,j} = F\, p_j q_j$ to leading order (e.g., Roze, 2015), and equations A4 – A9 can thus be simplified to:

$$a_{j,\emptyset} \approx -sh + \left[2e_{\mathrm{axa}} - s^2 h\left(1-2h\right)F\right]\sum_k p_k \quad \text{(A10)}$$

$$a_{j,j} \approx -s\left(1-2h\right) + \left[2e_{\text{axd}} - s^2\left(1-2h\right)^2 F\right] \sum_k p_k \tag{A11}$$

$$a_{jk,\emptyset} = a_{j,k} \approx e_{\text{axa}} + s^2 h^2 \tag{A12}$$

$$a_{jk,j} \approx e_{\text{axd}} + s^2 h\left(1-2h\right) \tag{A13}$$

$$a_{jk,jk} \approx e_{\text{dxd}} + s^2\left(1-2h\right)^2 . \tag{A14}$$

These approximations also hold when $s$, $e_{\text{axa}}$, $e_{\text{axd}}$ and $e_{\text{dxd}}$ are all of order $\epsilon$, in which case the terms in $s^2$ may be neglected in the expressions above.

Charlesworth et al. (1991) used the following fitness function to explore the effects of synergistic epistasis on the mutation load and inbreeding depression in diploids:

$$W = e^{-\alpha n - \frac{\beta}{2} n^2} \tag{A15}$$

where $n = h n_{\text{he}} + n_{\text{ho}}$, $n_{\text{he}}$ and $n_{\text{ho}}$ being the number of heterozygous and homozygous mutations in the genome of the individual, respectively. Here we use a slightly modified version ensuring that the $\beta$ term (representing epistasis) vanishes when only one locus carries a deleterious allele:

$$W = \left(1-hs\right)^{n_{\text{he}}}\left(1-s\right)^{n_{\text{ho}}}\left(1-\beta h^2\right)^{\frac{1}{2}n_{\text{he}}\left(n_{\text{he}}-1\right)}\left(1-\beta h\right)^{n_{\text{he}}n_{\text{ho}}}\left(1-\beta\right)^{\frac{1}{2}n_{\text{ho}}\left(n_{\text{ho}}-1\right)} . \tag{A16}$$

Equation A16 becomes equivalent to equation A15 (with $\alpha = s$) when $n_{\text{he}}$ and $n_{\text{ho}}$ are large (so that $n_{\text{he}} - 1 \approx n_{\text{he}}$ and $n_{\text{ho}} - 1 \approx n_{\text{ho}}$), and when $s$ and $\beta$ are small. Assuming that $s$ is of order $\epsilon$ while $\beta$ is of order $\epsilon^2$, equation A16 may be written as:

$$W = 1 - s\sum_j H_j + \left(s^2 - \beta\right)\sum_{j<k} H_j H_k + o\!\left(\epsilon^2\right) \tag{A17}$$

with $H_j = h\left(X_j^{\text{M}} + X_j^{\text{P}}\right) + \left(1-2h\right) X_j^{\text{M}} X_j^{\text{P}}$. This yields the same expressions for $a_{\mathbb{U},\mathbb{V}}$ coefficients as above, replacing $e_{\text{axa}}$, $e_{\text{axd}}$ and $e_{\text{dxd}}$ by $-\beta h^2$, $-\beta h\left(1-2h\right)$ and

$-\beta\,(1-2h)^2$, respectively. When using Charlesworth et al.'s (1991) fitness function (equation A15), it can be shown that extra terms $-\frac{\beta}{2}h^2$ and $-\left(\frac{1}{2}-h^2\right)\beta$ appear in the expressions of $a_j$ and $a_{j,j}$, respectively. Figure 1D in the main text shows simulation results obtained when fitness is given by equation A16, but simulations using equation A15 gave very similar results (not shown).

**Gaussian stabilizing selection.** We now derive expression for $a_{\mathbb{U},\mathbb{V}}$ selection coefficients in the case of the multivariate, Gaussian fitness function given by equation 14 in the main text. In the following we assume weak selection, that is, ${g_\alpha}^2/V_{\mathrm{s}}$ is small, of order $\epsilon$. A Taylor series of equation 14 to the second order in $\epsilon$ yields:

$$W = 1 - \frac{\sum_\alpha {g_\alpha}^2}{2V_{\mathrm{s}}} + \frac{1}{2}\left(\frac{\sum_\alpha {g_\alpha}^2}{2V_{\mathrm{s}}}\right)^2 + o\big(\epsilon^2\big)\,. \tag{A18}$$

using equations 3 and 13 in the main text and assuming that mean trait values are at the optimum ($\overline{g_\alpha} = 2\sum_j r_{\alpha j}\, p_j = 0$), we have $g_\alpha = \sum_j r_{\alpha j}\left(\zeta_j^{\mathrm{M}} + \zeta_j^{\mathrm{P}}\right)$, and equation A18 can be written as:

$$\begin{aligned} W =& 1 + \sum_{j,k} b_{jk}\left(\zeta_{jk,\emptyset} + \zeta_{j,k} + \zeta_{k,j} + \zeta_{\emptyset,jk}\right) \\ &+ \frac{1}{2}\sum_{j,k,l,m} b_{jk}b_{lm}\Big(\zeta_{jklm,\emptyset} + \zeta_{jkl,m} + \zeta_{jkm,l} + \zeta_{jk,lm} \\ &\qquad + \zeta_{jlm,k} + \zeta_{jl,km} + \zeta_{jm,kl} + \zeta_{j,klm} \\ &\qquad + \zeta_{klm,j} + \zeta_{kl,jm} + \zeta_{km,jl} + \zeta_{k,jlm} \\ &\qquad + \zeta_{lm,jk} + \zeta_{l,jkm} + \zeta_{m,jkl} + \zeta_{\emptyset,jklm}\Big)\Bigg] + o\big(\epsilon^2\big) \end{aligned} \tag{A19}$$

where $b_{jk} = -\sum_\alpha r_{\alpha j} r_{\alpha k}/\left(2V_{\mathrm{s}}\right)$, of order $\epsilon$. The first sum of equation A19 is over all pairs of loci $j$ and $k$ (including $j = k$), each pair with $j \neq k$ being counted twice, while the second sum is over all possible quadruplets of loci $j$, $k$, $l$, $m$. Mean fitness is given by the same expression, replacing $\zeta_{\mathbb{U},\mathbb{V}}$ by $D_{\mathbb{U},\mathbb{V}}$. To the second order in $\epsilon$, this yields:

$$\begin{aligned}
\frac{W}{\overline{W}} = 1 + \sum_{j,k} b_{jk} &\left[\left(\zeta_{jk,\emptyset} + \zeta_{\emptyset,jk} - 2\tilde{D}_{jk}\right) + \left(\zeta_{j,k} + \zeta_{k,j} - 2\tilde{D}_{j,k}\right)\right] \\
&\times \left[1 - 2\sum_{l,m} b_{lm}\left(\tilde{D}_{lm} + \tilde{D}_{l,m}\right)\right] \\
+ \frac{1}{2}\sum_{j,k,l,m} b_{jk}b_{lm} &\Bigg[\left(\zeta_{jklm,\emptyset} + \zeta_{\emptyset,jklm} - 2\tilde{D}_{jklm}\right) + \left(\zeta_{jkl,m} + \zeta_{m,jkl} - 2\tilde{D}_{jkl,m}\right) \\
&+ \left(\zeta_{jkm,l} + \zeta_{l,jkm} - 2\tilde{D}_{jkm,l}\right) + \left(\zeta_{jlm,k} + \zeta_{k,jlm} - 2\tilde{D}_{jlm,k}\right) \\
&+ \left(\zeta_{klm,j} + \zeta_{j,klm} - 2\tilde{D}_{klm,j}\right) + \left(\zeta_{jk,lm} + \zeta_{lm,jk} - 2\tilde{D}_{jk,lm}\right) \\
&+ \left(\zeta_{jl,km} + \zeta_{km,jl} - 2\tilde{D}_{jl,km}\right) + \left(\zeta_{jm,kl} + \zeta_{kl,jm} - 2\tilde{D}_{jm,kl}\right)\Bigg] + o\left(\epsilon^2\right).
\end{aligned} \tag{A20}$$

Expressions for $a_{\mathbb{U},\mathbb{V}}$ coefficients can be obtained by eliminating repeated indices in $\zeta_{\mathbb{U},\mathbb{V}}$ variables using the relation (e.g., eq. 5 in Kirkpatrick et al., 2002):

$$\zeta_{\mathbb{U}jj,\mathbb{V}} = p_j q_j \, \zeta_{\mathbb{U},\mathbb{V}} + (1 - 2p_j)\, \zeta_{\mathbb{U}j,\mathbb{V}} \tag{A21}$$

and finding the coefficient of $\zeta_{\mathbb{U},\mathbb{V}}$ in equation A20 for each $\mathbb{U}$, $\mathbb{V}$ (neglecting terms involving more than 2 loci). Using the fact that $\tilde{D}_{jk}$ and $\tilde{D}_{j,k}$ are of order $\epsilon$ at equilibrium for $j \neq k$, one obtains:

$$a_{j,\emptyset} = \frac{a_{j,j}}{2}\,(1 - 2p_j) + o\left(\epsilon^2\right) \tag{A22}$$

$$a_{j,j} = 2b_{jj}\left[1 - 2\sum_k b_{kk}\,(1 + F)\, p_k q_k\right] + \sum_k a_{jk,jk}\, p_k q_k + o\left(\epsilon^2\right) \tag{A23}$$

$$a_{jk,\emptyset} = a_{j,k} = 2b_{jk} + o(\epsilon) \tag{A24}$$

$$a_{jk,j} = \frac{a_{jk,jk}}{2}\,(1 - 2p_k) + o\left(\epsilon^2\right) \tag{A25}$$

$$a_{jk,jk} = 4\left(b_{jj}b_{kk} + 2{b_{jk}}^2\right) + o\left(\epsilon^2\right). \tag{A26}$$

Note that the coefficient $a_{jk,j}$ also includes a term $12 b_{jj} b_{jk}\,(1 - 2p_j)$, but this term will vanish when summed over all $j$, $k$ (and when computations are performed to the order

$\epsilon^2$), since the average of $r_{\alpha k}$ over all loci is zero (no mutational bias).

**Non-Gaussian stabilizing selection.** We now derive two-locus $a_{\mathbb{U},\mathbb{V}}$ coefficients in the more general case where fitness declines as $d^Q$ with the distance $d$ from the phenotypic optimum (equation 15 in the main text). Weak selection implies that $({g_\alpha}^2/V_\mathrm{s})^{Q/2}$ is small, of order $\epsilon$. We will first derive expressions to the first order in $\epsilon$, and then to the second order. The derivations below assume that $Q$ is an even integer; however, simulations indicate that the approximations obtained may also hold when $Q$ is odd (not shown). We have:

$$\begin{aligned} W &= 1 - \left(\frac{\sum_\alpha {g_\alpha}^2}{2V_\mathrm{s}}\right)^{\frac{Q}{2}} + o(\epsilon) \\ &= 1 - \frac{1}{(2V_\mathrm{s})^{\frac{Q}{2}}} \sum_{\alpha,\beta,\gamma,\ldots} \underbrace{{g_\alpha}^2 {g_\beta}^2 {g_\gamma}^2 \ldots}_{\frac{Q}{2}\text{ elements}} + o(\epsilon)\,. \end{aligned} \tag{A27}$$

In the following we assume that average trait values are at the phenotypic optimum ($\overline{g_\alpha} = 0$), so that (from equation 13 in the main text) $g_\alpha = \sum_i r_{\alpha i}\left(\zeta_j^\mathrm{M} + \zeta_j^\mathrm{P}\right)$. We thus have (to the first order in $\epsilon$):

$$W = 1 - \frac{1}{(2V_\mathrm{s})^{\frac{Q}{2}}} \sum_{j,k,l,m\ldots} \left( \sum_{\alpha,\beta,\gamma,\ldots} r_{\alpha j} r_{\alpha k} r_{\beta l} r_{\beta m} \ldots \right) \left(\zeta_j^\mathrm{M} + \zeta_j^\mathrm{P}\right)\left(\zeta_k^\mathrm{M} + \zeta_k^\mathrm{P}\right)\left(\zeta_l^\mathrm{M} + \zeta_l^\mathrm{P}\right)\ldots \tag{A28}$$

where the first sum (over loci) comprises $Q$ elements. From this, we obtain:

$$\frac{W}{\overline{W}} = 1 - \frac{1}{(2V_\mathrm{s})^{\frac{Q}{2}}} \sum_{j,k,l,m\ldots} \left( \sum_{\alpha,\beta,\gamma,\ldots} r_{\alpha j} r_{\alpha k} r_{\beta l} r_{\beta m} \ldots \right) \sum_{\mathbb{S}+\mathbb{T}=jklm\ldots} \left(\zeta_{\mathbb{S},\mathbb{T}} - D_{\mathbb{S},\mathbb{T}}\right) + o(\epsilon) \tag{A29}$$

where the last sum is over all possible partitions of the set of $Q$ loci ($jklm\ldots$) into two sets $\mathbb{S}$ and $\mathbb{T}$. Using equation A21, one can see that the terms $\zeta_{j,\emptyset}$, $\zeta_{\emptyset,j}$ will necessarily arise from terms in which all indices in $\mathbb{S}$ and $\mathbb{T}$ are repeated. For example for $Q = 4$, we have

$$\zeta_{jjkk,\emptyset} - D_{jjkk,\emptyset} = (1-2p_j)\, p_k q_k\, \zeta_{j,\emptyset} + (1-2p_k)\, p_j q_j\, \zeta_{k,\emptyset} + (1-2p_j)(1-2p_k)(\zeta_{jk,\emptyset} - D_{jk,\emptyset}), \tag{A30}$$

$$\zeta_{jj,kk} - D_{jj,kk} = (1-2p_j)\, p_k q_k\, \zeta_{j,\emptyset} + (1-2p_k)\, p_j q_j\, \zeta_{\emptyset,k} + (1-2p_j)(1-2p_k)(\zeta_{j,k} - D_{j,k}). \tag{A31}$$

Note that the terms $\zeta_{jjjj,\emptyset} - D_{jjjj,\emptyset}$, $\zeta_{jj,jj} - D_{jj,jj}$ will also generate terms in $\zeta_{j,\emptyset}$, but we will neglect those based on the fact that they will be summed over a lower number of loci (assuming that the number of loci is large). From this, the coefficient $a_{j,\emptyset}$ is given by (still for $Q = 4$):

$$a_{j,\emptyset} = -\frac{2(1-2p_j)}{(2V_s)^2} \sum_k \sum_{\alpha,\beta} \left( r_{\alpha j}^2 r_{\beta k}^2 + 2 r_{\alpha j} r_{\beta j} r_{\alpha k} r_{\beta k} \right) \times 2 p_k q_k. \tag{A32}$$

We can then note that $2\sum_k \sum_\beta r_{\beta k}^2\, p_k q_k = n V_g^0$ (where $V_g^0$ is the genic variance, the same for all traits at equilibrium), while $2\sum_k r_{\alpha k} r_{\beta k}\, p_k q_k$ differs from zero only for $\alpha = \beta$ (due to the fact that the average of $r_{\alpha k}$ over all loci is zero) and equals $V_g^0$ in this case. Therefore,

$$a_{j,\emptyset} = -\frac{2(n+2)\, V_g^0}{(2V_s)^2} \sum_\alpha r_{\alpha j}^2 (1-2p_j). \tag{A33}$$

The same reasoning extends to other values of $Q$ (as long as $Q$ is even). In particular, the factor 2 in equations A32 and A33 (that stemmed from the fact that both the terms $\zeta_{j,\emptyset}$ and $\zeta_{k,\emptyset}$ of equation A30 contribute to $a_{j,\emptyset}$) becomes $Q/2$, while $V_g^0$ in the same equations becomes $\left(V_g^0\right)^{\frac{Q}{2}-1}$. The factor $n+2$ can be extended as follows. Consider the term $r_{\alpha j} r_{\alpha k} r_{\beta l} r_{\beta m} \ldots$ in equation A29. Indices $j$, $k$, $l \ldots$ correspond to loci, that have to be paired (in order to obtain repeated indices). A pair may occur "within the same trait" (for example, the pair $j = k$ is within the trait $\alpha$) or "connect" two traits (for example, $j = l$ connects traits $\alpha$ and $\beta$), in which case these traits will have to be the same when summing over loci (otherwise the sum equals zero). Two traits can be connected, and disconnected from the other traits in two ways (for example, $\alpha$ and $\beta$ can be connected by $j = l$, $k = m$ or by $j = m$, $k = l$). Similarly, 3 traits can be connected (and disconnected from the other traits) in 8 possible ways. For $Q = 6$, $\left(V_g^0\right)^2$ will thus be multiplied by a factor $n^2 + 3 \times 2n + 8 = (n+2)(n+4)$ (the factor 3 stems from the fact that trait $\alpha$ can be paired with $\beta$, or $\alpha$ with $\gamma$, or $\beta$ with $\gamma$. It is possible to show by recursion that for higher values of $Q$, this factor becomes:

$$(n+2)(n+4)\ldots(n+Q-2) = \frac{2^{\frac{Q}{2}}}{n} \frac{\Gamma\left(\frac{Q+n}{2}\right)}{\Gamma\left(\frac{n}{2}\right)}. \tag{A34}$$

Therefore, we finally obtain:

$$a_j = -\frac{Q}{2nV_g^0} \left(\frac{V_g^0}{V_\mathrm{s}}\right)^{\frac{Q}{2}} \frac{\Gamma\left(\frac{Q+n}{2}\right)}{\Gamma\left(\frac{n}{2}\right)} \sum_\alpha r_{\alpha j}^2 \left(1 - 2p_j\right). \tag{A35}$$

Using a similar reasoning as above, one arrives at:

$$a_{j,j} = -\frac{Q}{nV_g^0} \left(\frac{V_g^0}{V_\mathrm{s}}\right)^{\frac{Q}{2}} \frac{\Gamma\left(\frac{Q+n}{2}\right)}{\Gamma\left(\frac{n}{2}\right)} \sum_\alpha r_{\alpha j}^2. \tag{A36}$$

Equations A35 can be used to compute the genic variance $V_g^0$ at equilibrium under random mating ($\sigma = 0$), neglecting linkage disequilibria and other associations between loci (see Supplementary File S2 for an expression when $\sigma > 0$). Indeed, the change in $p_j q_j = \tilde{D}_{jj}$ during selection (to the first order in $\epsilon$) is given by:

$$\Delta_\mathrm{sel} \tilde{D}_{jj} = a_j \left(1 - 2p_j\right) p_j q_j + o(\epsilon) \tag{A37}$$

(see Supplementary File S2), while the change due to mutation is (to the first order in $u$):

$$\Delta_\mathrm{mut} \tilde{D}_{jj} = u \left(1 - 2p_i\right)^2. \tag{A38}$$

At mutation-selection balance, we thus have either $p_j = 1/2$ (if selection is weak relative to mutation), or

$$\frac{Q}{2nV_g^0}\left(\frac{V_g^0}{V_{\mathrm{s}}}\right)^{\frac{Q}{2}}\frac{\Gamma\left(\frac{Q+n}{2}\right)}{\Gamma\left(\frac{n}{2}\right)}\sum_{\alpha} r_{\alpha j}^2\, p_j q_j = u\,. \tag{A39}$$

Summing the last expression over all loci yields:

$$\frac{Q}{2nV_g^0}\left(\frac{V_g^0}{V_{\mathrm{s}}}\right)^{\frac{Q}{2}}\frac{\Gamma\left(\frac{Q+n}{2}\right)}{\Gamma\left(\frac{n}{2}\right)}\frac{nV_g^0}{2} = U \tag{A40}$$

and thus:

$$\left(\frac{V_g^0}{V_{\mathrm{s}}}\right)^{\frac{Q}{2}}\frac{\Gamma\left(\frac{Q+n}{2}\right)}{\Gamma\left(\frac{n}{2}\right)} = \frac{4U}{Q}. \tag{A41}$$

From equations A35 – A36 and A41, $a_j$ and $a_{j,j}$ may be written as:

$$a_j = -\frac{2U}{nV_g^0}\sum_{\alpha} r_{\alpha j}^2\left(1-2p_j\right) \tag{A42}$$

$$a_{j,j} = -\frac{4U}{nV_g^0}\sum_{\alpha} r_{\alpha j}^2 \tag{A43}$$

when mating is random. From equation 36 in the main text, and assuming that most recombination rates between loci are sufficiently large relative to the strength of selection $a_j$, the effect of purging on the spread of a selfing modifier (when $\sigma = 0$) depends on the sum $\sum_j a_j\left[a_j + a_{j,j}\left(1-2p_j\right)\right] p_j q_j$. From equations A42 and A43, this term equals $6U^2 a^2/V_g^0$, where $a^2$ is the variance of $r_{\alpha j}$ across loci. Using the fact that the average deleterious effect of a heterozygous mutation (in an optimal genotype) is given by

$$\overline{\varsigma} = \left(\frac{a^2}{V_{\mathrm{s}}}\right)^{\frac{Q}{2}}\frac{\Gamma\left(\frac{Q+n}{2}\right)}{\Gamma\left(\frac{n}{2}\right)} \tag{A44}$$

(e.g., Gros et al., 2009) and using equation A41 yields equation 40 in the main text.

Coefficients $a_{jk,\emptyset}$ and $a_{j,k}$ are obtained as above. For $Q = 4$, terms in $\zeta_{jk,\emptyset}$, $\zeta_{j,k}$ arise in equations A30 – A31, but also from terms such as $\zeta_{jkll,\emptyset} - D_{jkll,\emptyset}$, $\zeta_{jk,ll} - D_{jk,ll}$,

$\zeta_{jll,k} - D_{jll,k}$, $\zeta_{j,kll} - D_{j,kll}$ (with $j \neq k \neq l$). Here again, we will neglect terms that are summed over lower numbers of loci, and thus neglect the terms arising from equations A30 – A31. Using the same reasoning as for $a_j$, $a_{j,j}$, this yields:

$$a_{jk} = a_{j,k} = -\frac{Q}{nV_g^0}\left(\frac{V_g^0}{V_{\mathrm{s}}}\right)^{\frac{Q}{2}} \frac{\Gamma\left(\frac{Q+n}{2}\right)}{\Gamma\left(\frac{n}{2}\right)} \sum_{\alpha} r_{\alpha j} r_{\alpha k} \tag{A45}$$

and thus, using equation A41:

$$a_{jk} = a_{j,k} = -\frac{4U}{nV_g^0} \sum_{\alpha} r_{\alpha j} r_{\alpha k} \tag{A46}$$

at mutation-selection balance when $\sigma = 0$, to leading order. Under random mating, the linkage disequilibrium $D_{jk}$ is given by:

$$D_{jk} \approx \frac{a_{jk}}{\rho_{jk}} p_j q_j p_k q_k \tag{A47}$$

to leading order (see Supplementary File S2), and the effect of linkage disequilibria on the genetic variance is thus:

$$2 \sum_{j \neq k} r_{\alpha j} r_{\alpha k} D_{jk} \approx -2U V_g^0 / \left(n \rho_{\mathrm{H}}\right) \tag{A48}$$

where $\rho_{\mathrm{H}}$ is the harmonic mean recombination rate between loci affecting the traits. Furthermore, one obtains from equation A46

$$\sum_{j,k} {a_{jk}}^2 p_j q_j p_k q_k \approx \frac{4U^2}{n}, \tag{A49}$$

independent of $Q$.

Coefficients $a_{jk,j}$ and $a_{jk,jk}$ are also derived using the same method. For $Q = 4$, the term $\zeta_{jkk,j} - D_{jkk,j}$ appears twice in equation A29 and generate terms in $(1 - 2p_k)\left(\zeta_{jk,j} - D_{jk,j}\right)$, while the term $\zeta_{jk,jk} - D_{jk,jk}$ appears 4 times, yielding:

$$a_{jk,j} = -\frac{4}{\left(2V_{\mathrm{s}}\right)^2}\left(1 - 2p_k\right)\left(S_{jj}S_{kk} + 2{S_{jk}}^2\right) \tag{A50}$$

$$a_{jk,jk} = -\frac{8}{\left(2V_{\mathrm{s}}\right)^2}\left(S_{jj}S_{kk} + 2{S_{jk}}^2\right) \tag{A51}$$

with $S_{jk} = \sum_\alpha r_{\alpha j} r_{\alpha k}$ (the extra factor 2 arises from the fact that each pair of loci $j$, $k$ is counted only once in equation A1, but twice in equation A29). Generalizing to other (even) values of $Q$, one obtains:

$$a_{jk,j} = -\frac{Q\left(Q-2\right)}{2n\left(n+2\right)\left(V_g^0\right)^2}\left(\frac{V_g^0}{V_{\mathrm{s}}}\right)^{\frac{Q}{2}}\frac{\Gamma\left(\frac{Q+n}{2}\right)}{\Gamma\left(\frac{n}{2}\right)}\left(1-2p_k\right)\left(S_{jj}S_{kk} + 2{S_{jk}}^2\right) \tag{A52}$$

$$a_{jk,jk} = -\frac{Q\left(Q-2\right)}{n\left(n+2\right)\left(V_g^0\right)^2}\left(\frac{V_g^0}{V_{\mathrm{s}}}\right)^{\frac{Q}{2}}\frac{\Gamma\left(\frac{Q+n}{2}\right)}{\Gamma\left(\frac{n}{2}\right)}\left(S_{jj}S_{kk} + 2{S_{jk}}^2\right). \tag{A53}$$

Using equation A41, we thus have at mutation-selection balance and when $\sigma = 0$:

$$a_{jk,j} = -\frac{2U\left(Q-2\right)}{n\left(n+2\right)\left(V_g^0\right)^2}\left(1-2p_k\right)\left(S_{jj}S_{kk} + 2{S_{jk}}^2\right) \tag{A54}$$

$$a_{jk,jk} = -\frac{4U\left(Q-2\right)}{n\left(n+2\right)\left(V_g^0\right)^2}\left(S_{jj}S_{kk} + 2{S_{jk}}^2\right). \tag{A55}$$

In Supplementary File S2, we derive an expression for inbreeding depression in a randomly mating population to the second order in $U$, under the fitness function given by equation 15 in the main text. For this, the coefficients $a_j$, $a_{j,j}$ and $a_{jk,jk}$ need to be expressed to the second order in $\epsilon$. This can be achieved using the same method as above, starting from:

$$W = 1 - \left(\frac{\sum_\alpha {g_\alpha}^2}{2V_{\mathrm{s}}}\right)^{\frac{Q}{2}} + \frac{1}{2}\left(\frac{\sum_\alpha {g_\alpha}^2}{2V_{\mathrm{s}}}\right)^{Q} + o\!\left(\epsilon^2\right). \tag{A56}$$

Assuming random mating, this finally yields:

$$a_j = -\frac{Q}{2nV_g^0}\left[Z\left(Q,n\right)\left[1 + Z\left(Q,n\right)\right] - Z\left(2Q,n\right)\right]\sum_\alpha r_{\alpha j}^2\left(1-2p_j\right) \tag{A57}$$

$$a_{j,j} = -\frac{Q}{nV_g^0}\left[Z\left(Q,n\right)\left[1 + Z\left(Q,n\right)\right] - Z\left(2Q,n\right)\right]\sum_\alpha r_{\alpha j}^2 \tag{A58}$$

$$a_{jk,jk} = -\frac{Q}{n\left(n+2\right)\left(V_g^0\right)^2}\left[\left(Q-2\right)Z\left(Q,n\right)\left[1+Z\left(Q,n\right)\right]-\left(2Q-2\right)Z\left(2Q,n\right)\right]$$
$$\times\left(S_{jj}S_{kk}+2{S_{jk}}^2\right) \tag{A59}$$

with:

$$Z\left(Q,n\right) = \left(\frac{V_g^0}{V_\mathrm{s}}\right)^{\frac{Q}{2}}\frac{\Gamma\left(\frac{Q+n}{2}\right)}{\Gamma\left(\frac{n}{2}\right)}. \tag{A60}$$

It can be verified that equations A57 – A59 are equivalent to equations A22, A23 and A26 in the case of a Gaussian fitness function ($Q = 2$) and under random mating ($\sigma = 0$).
