## Supplementary File S2 for "Epistasis, inbreeding depression and the evolution of self-fertilization"

### FILE S2: GENERAL EXPRESSIONS FOR INBREEDING DEPRESSION

**Outcrossing population.** A general expression for inbreeding depression, including the effects of interactions between pairs of loci can be obtained from equation A1 in Supplementary File S1. We will first consider the case of a fully outcrossing population ($\sigma = 0$). Using the fact that associations $\tilde{D}_{j,j}$, $\tilde{D}_{j,k}$ and $\tilde{D}_{jk,j}$ equal zero among offspring produced by random mating, the mean fitness of offspring produced by mating between randomly sampled parents divided by the mean fitness of the parental population is given by:

$$\frac{\overline{W}^{\text{out}}}{\overline{W}} = 1 + 2\sum_{j<k} a_{jk}\left(\tilde{D}_{jk}^{\text{out}} - \tilde{D}_{jk}\right) + \sum_{j<k} a_{jk,jk}\left(D_{jk,jk}^{\text{out}} - D_{jk,jk}\right) \tag{B1}$$

where $\tilde{D}_{jk}^{\text{out}}$ and $D_{jk,jk}^{\text{out}}$ are measured among offspring, and $\tilde{D}_{jk}$, $D_{jk,jk}$ among parents. The association $D_{jk,jk}$ equals $\tilde{D}_{jk}{}^2$ under random mating, while $\tilde{D}_{jk}$ is reduced by a factor $1 - \rho_{jk}$ by recombination (assuming no selection among parents), giving:

$$\frac{\overline{W}^{\text{out}}}{\overline{W}} = 1 - 2\sum_{j<k} \rho_{jk}\, a_{jk}\, \tilde{D}_{jk} - \sum_{j<k} \rho_{jk}\left(2 - \rho_{jk}\right) a_{jk,jk}\, \tilde{D}_{jk}{}^2 \,. \tag{B2}$$

The mean fitness of offspring produced by selfing from randomly sampled parents, divided by the mean fitness of the parental population is given by (from equation A1):

$$\begin{aligned}\frac{\overline{W}^{\text{self}}}{\overline{W}} = 1 &+ \sum_{j} a_{j,j}\, D_{j,j}^{\text{self}} + 2\sum_{j<k} a_{jk}\left(\tilde{D}_{jk}^{\text{self}} - \tilde{D}_{jk}\right) \\ &+ 2\sum_{j<k} a_{j,k}\, \tilde{D}_{j,k}^{\text{self}} + 2\sum_{j,k} a_{jk,j}\, \tilde{D}_{jk,j}^{\text{self}} + 2\sum_{j<k} a_{jk,k}\, \tilde{D}_{jk,k}^{\text{self}} \\ &+ \sum_{j<k} a_{jk,jk}\left(D_{jk,jk}^{\text{self}} - \tilde{D}_{jk}{}^2\right).\end{aligned} \tag{B3}$$

Associations $D_{\mathbb{U},\mathbb{V}}^{\text{self}}$ in the expression above are measured among selfed offspring. Taking into account the possible configurations of genes within parental individuals, and using equation A21 in Supplementary File S1 to eliminate repeated indices, one obtains:

$$D^{\text{self}}_{j,j} = \frac{1}{2} p_j q_j, \quad \tilde{D}^{\text{self}}_{jk} = (1 - \rho_{jk})\, \tilde{D}_{jk}, \quad \tilde{D}^{\text{self}}_{j,k} = \frac{1}{2} \tilde{D}_{jk}, \tag{B4}$$

$$D^{\text{self}}_{jk,j} = \frac{1}{2} (1 - 2p_j)(1 - \rho_{jk})\, \tilde{D}_{jk}, \quad \tilde{D}^{\text{self}}_{jk,k} = \frac{1}{2} (1 - 2p_k)(1 - \rho_{jk})\, \tilde{D}_{jk}, \tag{B5}$$

$$D^{\text{self}}_{jk,jk} = \frac{1}{2} \left[1 - 2\rho_{jk}(1 - \rho_{jk})\right] \left(p_j q_j p_k q_k + \tilde{D}_{jk}{}^2\right). \tag{B6}$$

Inbreeding depression is defined as

$$\delta = 1 - \frac{\overline{W}^{\text{self}}}{\overline{W}^{\text{out}}} = 1 - \frac{\overline{W}^{\text{self}} / \overline{W}}{\overline{W}^{\text{out}} / \overline{W}}. \tag{B7}$$

From the expressions above, we have to the second order in $a_{\mathbb{U},\mathbb{V}}$ coefficients (and thus neglecting the terms in $\tilde{D}_{jk}{}^2$):

$$\begin{aligned} \delta \approx & -\frac{1}{2} \sum_j a_{j,j}\, p_j q_j - \frac{1}{2} \sum_{j<k} a_{jk,jk} \left[1 - 2\rho_{jk}(1 - \rho_{jk})\right] p_j q_j p_k q_k - \sum_{j<k} a_{j,k}\, \tilde{D}_{jk} \\ & - \sum_{j<k} a_{jk,j} (1 - 2p_j)(1 - \rho_{jk})\, \tilde{D}_{jk} - \sum_{j<k} a_{jk,k} (1 - 2p_k)(1 - \rho_{jk})\, \tilde{D}_{jk} \end{aligned} \tag{B8}$$

which corresponds to equation 17 in the main text.

When the deleterious mutation rate is high, equation B8 may be higher than 1 (while from equation B7, inbreeding depression cannot be higher than 1). In general, more accurate expressions for parameter values leading to high inbreeding depression can be obtained by assuming that the effects of individual loci (and their interactions) on $\delta$ do multiply (rather than sum). Using the present framework, and to the first order in $a_{\mathbb{U},\mathbb{V}}$ coefficients, this can be achieved by assuming that the variance in log-fitness among individuals stays small, so that $\overline{W} \approx e^{\overline{\ln W}}$, leading to $\delta \approx 1 - e^{\overline{\ln W}_{\text{self}} - \overline{\ln W}_{\text{out}}}$. To the first order in $a_{\mathbb{U},\mathbb{V}}$ coefficients, equation 8 in the main text yields $\ln W - \ln \overline{W} \approx \sum_{\mathbb{U},\mathbb{V}} a_{\mathbb{U},\mathbb{V}} \left(\zeta_{\mathbb{U},\mathbb{V}} - D_{\mathbb{U},\mathbb{V}}\right)$. Using the same reasoning as above, one obtains (still to the first order in $a_{\mathbb{U},\mathbb{V}}$ coefficients) $\overline{\ln W}_{\text{out}} - \ln \overline{W} \approx 0$, while

$\overline{\ln W}_{\text{self}} - \ln \overline{W} \approx \frac{1}{2}\sum_j a_{j,j}\, p_j q_j + \frac{1}{2}\sum_{j<k} a_{jk,jk}\left[1 - 2\rho_{jk}\left(1-\rho_{jk}\right)\right] p_j q_j p_k q_k$, leading to $\delta \approx 1 - \exp\left[\frac{1}{2}\sum_j a_{j,j}\, p_j q_j + \frac{1}{2}\sum_{j<k} a_{jk,jk}\left[1 - 2\rho_{jk}\left(1-\rho_{jk}\right)\right] p_j q_j p_k q_k\right]$ (which cannot be greater than 1). In a population undergoing partial selfing, the same reasoning leads to equation 24 in the main text.

Equation B8 is modified when inbreeding depression is measured after selection, that is, when the contribution of parents to the pools of selfed and outcrossed offspring is proportional to their fitness. In that case, we have $\tilde{D}_{jk}^{\text{out}} = \tilde{D}_{jk}^{\text{self}} = \left(1-\rho_{jk}\right)\tilde{D}'_{jk} + \rho_{jk}\,\tilde{D}'_{j,k}$ where $\tilde{D}'_{jk}$ and $\tilde{D}'_{j,k}$ are measured after selection, while $\tilde{D}_{j,k}^{\text{self}} = \frac{1}{2}\left(\tilde{D}'_{jk} + \tilde{D}'_{j,k}\right)$ and $\tilde{D}_{jk,j}^{\text{self}} = \frac{1}{2}\left(1-2p_j\right)\left[\left(1-\rho_{jk}\right)\tilde{D}'_{jk} + \rho_{jk}\,\tilde{D}'_{j,k}\right]$. This yields:

$$\begin{aligned}\delta' \approx & -\frac{1}{2}\sum_j a_{j,j}\, p_j q_j - \frac{1}{2}\sum_{j<k} a_{jk,jk}\left[1 - 2\rho_{jk}\left(1-\rho_{jk}\right)\right] p_j q_j p_k q_k \\ & - \sum_{j<k} a_{j,k}\left(\tilde{D}'_{jk} + \tilde{D}'_{j,k}\right) - \sum_{j,k} a_{jk,j}\left(1-2p_j\right)\left[\left(1-\rho_{jk}\right)\tilde{D}'_{jk} + \rho_{jk}\,\tilde{D}'_{j,k}\right]\end{aligned} \tag{B9}$$

where $\delta'$ stands for inbreeding depression measured after selection.

**Partially selfing population.** Under partial selfing, equation B1 becomes:

$$\begin{aligned}\frac{\overline{W}^{\text{out}}}{\overline{W}} = & \, 1 - \sum_j a_{j,j}\, D_{j,j} + 2\sum_{j<k} a_{jk}\left(\tilde{D}_{jk}^{\text{out}} - \tilde{D}_{jk}\right) - 2\sum_{j<k} a_{j,k}\, D_{j,k} \\ & - 2\sum_{j<k} a_{jk,j}\, D_{jk,j} - 2\sum_{j<k} a_{jk,k}\, D_{jk,k} + \sum_{j<k} a_{jk,jk}\left(D_{jk,jk}^{\text{out}} - D_{jk,jk}\right)\end{aligned} \tag{B10}$$

while equation B3 becomes:

$$\begin{aligned}\frac{\overline{W}^{\text{self}}}{\overline{W}} = & \, 1 + \sum_j a_{j,j}\left(D_{j,j}^{\text{self}} - D_{j,j}\right) + 2\sum_{j<k} a_{jk}\left(\tilde{D}_{jk}^{\text{self}} - \tilde{D}_{jk}\right) \\ & + 2\sum_{j<k} a_{j,k}\left(\tilde{D}_{j,k}^{\text{self}} - \tilde{D}_{j,k}\right) + 2\sum_{j<k} a_{jk,j}\left(\tilde{D}_{jk,j}^{\text{self}} - \tilde{D}_{jk,j}\right) \\ & + 2\sum_{j<k} a_{jk,k}\left(\tilde{D}_{jk,k}^{\text{self}} - \tilde{D}_{jk,k}\right) + \sum_{j<k} a_{jk,jk}\left(D_{jk,jk}^{\text{self}} - D_{jk,jk}\right).\end{aligned} \tag{B11}$$

Associations among selfed and outcrossed offspring are given by:

$$D^{\text{self}}_{j,j} = \frac{1}{2}\left(p_j q_j + D_{j,j}\right), \tag{B12}$$

$$\tilde{D}^{\text{self}}_{jk} = \tilde{D}^{\text{out}}_{jk} = (1-\rho_{jk})\,\tilde{D}_{jk} + \rho_{jk}\,\tilde{D}_{j,k}, \quad D^{\text{out}}_{jk,jk} = \left(\tilde{D}^{\text{out}}_{jk}\right)^2, \tag{B13}$$

$$\tilde{D}^{\text{self}}_{j,k} = \frac{1}{2}\left(\tilde{D}_{jk} + \tilde{D}_{j,k}\right), \tag{B14}$$

$$\tilde{D}^{\text{self}}_{jk,j} = \frac{1}{2}\left[\tilde{D}_{jk,j} + (1-2p_j)\left[(1-\rho_{jk})\,\tilde{D}_{jk} + \rho_{jk}\,\tilde{D}_{j,k}\right]\right], \tag{B15}$$

$$\begin{aligned}\tilde{D}^{\text{self}}_{jk,jk} = \frac{1}{2}\Big[&(1-\rho_{jk})^2\left[p_j q_j p_k q_k + D_{jk,jk} + (1-2p_j)(1-2p_k)\,\tilde{D}_{jk}\right] \\ &+ 2\rho_{jk}(1-\rho_{jk})\left[p_j q_j\, D_{k,k} + p_k q_k\, D_{j,j} + (1-2p_j)\,\tilde{D}_{jk,k} + (1-2p_k)\,\tilde{D}_{jk,j}\right] \\ &+ {\rho_{jk}}^2\left[p_j q_j p_k q_k + D_{jk,jk} + (1-2p_j)(1-2p_k)\,\tilde{D}_{j,k}\right]\Big].\end{aligned} \tag{B16}$$

Throughout the following, $\epsilon$ stands for the order of magnitude of the largest of $a_{\mathbb{U},\mathbb{V}}$ coefficients in absolute value. Associations $\tilde{D}_{jk}$, $\tilde{D}_{j,k}$, $\tilde{D}_{jk,j}$, $\tilde{D}_{jk,k}$ are generated by selection and are thus of order $\epsilon$. By contrast, associations $D_{j,j}$ and $D_{jk,jk}$ are generated by partial selfing even in the absence of selection, in which case they equal $F\,p_j q_j$ and $(G_{jk} + F^2)\,p_j q_j p_k q_k$ at equilibrium (respectively), where $F$ is the inbreeding coefficient and $G_{jk}$ the identity disequilibrium between loci $j$ and $k$ (e.g., Roze, 2015). From this, one obtains to the second order in $\epsilon$:

$$\begin{aligned}\delta = &-\frac{1}{2}\sum_{j} a_{j,j}\left(p_j q_j + D_{j,j}\right)\left(1 + \sum_{k} a_{k,k}\, D_{k,k}\right) - \sum_{j<k} a_{j,k}\left(\tilde{D}_{jk} + \tilde{D}_{j,k}\right)\\
&-\sum_{j,k} a_{jk,j}\left[\tilde{D}_{jk,j} + \left(1-2p_j\right)\left[\left(1-\rho_{jk}\right)\tilde{D}_{jk} + \rho_{jk}\,\tilde{D}_{j,k}\right]\right]\\
&-\frac{1}{2}\sum_{j<k} a_{jk,jk}\Bigg[\left(1-\rho_{jk}\right)^2\left[p_j q_j p_k q_k + D_{jk,jk} + \left(1-2p_j\right)\left(1-2p_k\right)\tilde{D}_{jk}\right]\\
&\qquad + 2\rho_{jk}\left(1-\rho_{jk}\right)\Big[p_j q_j\, D_{k,k} + p_k q_k\, D_{j,j}\\
&\qquad\qquad + \left(1-2p_j\right)\tilde{D}_{jk,k} + \left(1-2p_k\right)\tilde{D}_{jk,j}\Big]\\
&\qquad + {\rho_{jk}}^2\left[p_j q_j p_k q_k + D_{jk,jk} + \left(1-2p_j\right)\left(1-2p_k\right)\tilde{D}_{j,k}\right]\Bigg].\end{aligned} \tag{B17}$$

To the first order in $\epsilon$ and when $\rho_{jk} = 1/2$, equation B17 is equivalent to equation 19 in the main text. Expressions for genetic associations to the first order in $\epsilon$ can be obtained using standard multilocus techniques (e.g., Kirkpatrick et al., 2002). Using a rare allele approximation — that is, neglecting terms in $\left(p_j q_j\right)^2$ — and assuming $a_{jk} = a_{j,k}$, one obtains:

$$D_{j,j} = F\left(p_j q_j + \Delta_{\text{sel}} D_{j,j}\right), \tag{B18}$$

$$\tilde{D}_{jk} = \frac{2 - \sigma\left(1 - 2\rho_{jk}\right)}{2\rho_{jk}\left(1-\sigma\right)}\Delta_{\text{sel}}\tilde{D}_{jk}, \quad \tilde{D}_{j,k} = \frac{\sigma\left(1 + 2\rho_{jk}\right)}{2\rho_{jk}\left(1-\sigma\right)}\Delta_{\text{sel}}\tilde{D}_{jk}, \tag{B19}$$

$$\tilde{D}_{jk,j} = F\left[\Delta_{\text{sel}}\tilde{D}_{jk,j} + \left(1-2p_j\right)\tilde{D}_{jk}\right], \tag{B20}$$

with

$$\Delta_{\text{sel}} D_{j,j} = \sum_{k}\Big(a_{k,k}\, G_{jk} + \left[a_{jk,k}\left(1-2p_j\right) + a_{jk,jk}\right]\left(G_{jk} + F^2\right)\Big) p_j q_j p_k q_k, \tag{B21}$$

$$\begin{aligned}\Delta_{\text{sel}}\tilde{D}_{jk} = &\Big[a_{jk}\left[\left(1+F\right)^2 + G_{jk}\right] + a_{jk,jk}\left(1-2p_j\right)\left(1-2p_k\right)\left(G_{jk} + F^2\right)\\
&+ \left[a_{jk,j}\left(1-2p_j\right) + a_{jk,k}\left(1-2p_k\right)\right]\left[F\left(1+F\right) + G_{jk}\right]\Big] p_j q_j p_k q_k,\end{aligned} \tag{B22}$$

$$\Delta_{\text{sel}}\tilde{D}_{jk,j} = \Big[\left[a_k + a_{k,k}\left(1-2p_k\right)\right]G_{jk} + \left[2a_{jk}\left(1-2p_j\right) + a_{jk,j}\right]\left[F\left(1+F\right) + G_{jk}\right] + \left[2a_{jk,k}\left(1-2p_j\right) + a_{jk,jk}\right]\left(1-2p_k\right)\left(G_{jk} + F^2\right)\Big]p_j q_j p_k q_k\,. \tag{B23}$$

Finally, $D_{jk,jk}$ is obtained by solving:

$$\tilde{D}_{jk,jk} = \frac{\sigma}{2}\Big[\left(1-\rho_{jk}\right)^2\left[p_j q_j p_k q_k + D_{jk,jk}{}' + \left(1-2p_j\right)\left(1-2p_k\right)\tilde{D}_{jk}{}'\right] + 2\rho_{jk}\left(1-\rho_{jk}\right)\left[p_j q_j\, D_{k,k}{}' + p_k q_k\, D_{j,j}{}' + \left(1-2p_j\right)\tilde{D}_{jk,k}{}' + \left(1-2p_k\right)\tilde{D}_{jk,j}{}'\right] + \rho_{jk}{}^2\left[p_j q_j p_k q_k + D_{jk,jk}{}' + \left(1-2p_j\right)\left(1-2p_k\right)\tilde{D}_{j,k}{}'\right]\Big], \tag{B24}$$

where associations $D_{\mathbb{U},\mathbb{V}}{}'$ are measured after selection and are given by $D_{\mathbb{U},\mathbb{V}}{}' = D_{\mathbb{U},\mathbb{V}} + \Delta_{\text{sel}}D_{\mathbb{U},\mathbb{V}}$, with:

$$\Delta_{\text{sel}}\tilde{D}_{jk,jk} = \Big[2a_j\left(1-2p_j\right) + 2a_k\left(1-2p_k\right) + a_{j,j} + a_{k,k} + 4a_{jk}\left(1-2p_j\right)\left(1-2p_k\right) + 2a_{jk,j}\left(1-2p_k\right) + 2a_{jk,k}\left(1-2p_j\right) + a_{jk,jk}\Big]\left(G_{jk} + F^2\right)p_j q_j p_k q_k\,. \tag{B25}$$

Computing $\delta$ also requires an expression for $p_j q_j$ at equilibrium. Assuming the same mutation rate $u$ between both alleles at the same locus, this can be obtained from $-\Delta_{\text{sel}}p_j q_j = u$, with:

$$\begin{aligned}\Delta_{\text{sel}}p_j q_j = {} & a_j\left(1-2p_j\right)\left(p_j q_j + D_{j,j}\right) + \sum_{k\neq j} a_k\left(1-2p_j\right)\left(\tilde{D}_{jk} + \tilde{D}_{j,k}\right) \\ & + a_{j,j}\, D_{j,j} + \sum_{k\neq j} a_{k,k}\left(1-2p_j\right)\tilde{D}_{jk,k} + \sum_{k\neq j} a_{jk,jk}\, D_{jk,jk} \\ & + \sum_{k\neq j} a_{jk}\left[\tilde{D}_{jk} + \tilde{D}_{j,k} + 2\left(1-2p_j\right)\tilde{D}_{jk,j}\right] + 2\sum_{k\neq j} a_{jk,j}\,\tilde{D}_{jk,j} \\ & + \sum_{k\neq j} a_{jk,k}\left[\tilde{D}_{jk,k} + \left(1-2p_j\right)\left(p_j q_j D_{k,k} + D_{jk,jk}\right)\right].\end{aligned} \tag{B26}$$

**Relation to previous results.** Previous expressions for inbreeding depression including the effect of identity disequilibria between pairs of loci (Roze, 2015) and of epistasis generated by Gaussian stabilizing selection (Abu Awad and Roze, 2018) can be recovered as special cases of the general equations given above. In the case of uniformly deleterious alleles without epistasis (Roze, 2015), selection coefficients are given by equations A10 – A14 in Supplementary File S1, with $e_{\mathrm{axa}} = e_{\mathrm{axd}} = e_{\mathrm{dxd}} = 0$. To the second order in $s$ and assuming free recombination among loci, the equations above simplify to:

$$\begin{aligned} \delta \approx & -\frac{1}{2} \sum_j a_{j,j} \left(p_j q_j + D_{j,j}\right) \left(1 + \sum_k a_{k,k} \, D_{k,k}\right) \\ & - \frac{1}{4} \sum_{j<k} a_{jk,jk} \left[\left(1+F\right)^2 + G\right] p_j q_j p_k q_k \end{aligned} \tag{B27}$$

where $G$ is the identity disequilibrium,

$$\begin{aligned} \Delta_{\mathrm{sel}} p_j q_j = & \; a_j \left(p_j q_j + D_{j,j}\right) + a_{j,j} \, D_{j,j} + \sum_{k \neq j} a_{k,k} \, \tilde{D}_{jk,k} \\ & + \sum_{k \neq j} a_{jk,k} \left(p_j q_j D_{k,k} + D_{jk,jk}\right) + \sum_{k \neq j} a_{jk,jk} \, D_{jk,jk} \end{aligned} \tag{B28}$$

with:

$$D_{j,j} = F \left[1 + \sum_{k \neq j} a_{k,k} \, G \, p_k q_k\right] p_j q_j, \tag{B29}$$

$$\tilde{D}_{jk,k} = F \left(a_j + a_{j,j}\right) G \, p_j q_j p_k q_k, \quad D_{jk,jk} = \left(G + F^2\right) p_j q_j p_k q_k \, . \tag{B30}$$

Equations B27 – B30 yield equation 14 in Roze (2015), expressed to the second order in $U$.

Selection coefficients under Gaussian stabilizing selection are given by equations A22 – A26 in Supplementary File S1. Approximating the identity disequilibrium $G_{jk}$

by its expression under free recombination ($G$), we have:

$$\delta = -\frac{1}{2}\sum_j a_{j,j}\left(p_j q_j + D_{j,j}\right)\left(1 + \sum_k a_{k,k}\, D_{k,k}\right) - \sum_{j<k} a_{jk}\left(\tilde{D}_{jk} + \tilde{D}_{j,k}\right) \\ - \frac{1}{4}\sum_{j<k} a_{jk,jk}\left[(1+F)^2 + G\right] p_j q_j p_k q_k \tag{B31}$$

$$\Delta_{\text{sel}} p_j q_j = a_j\left(1-2p_j\right)\left(p_j q_j + D_{j,j}\right) + a_{j,j}\, D_{j,j} + \sum_{k\neq j} a_{k,k}\left(1-2p_j\right)\tilde{D}_{jk,k} \\ + \sum_{k\neq j} a_{jk}\left[\tilde{D}_{jk} + \tilde{D}_{j,k} + 2\left(1-2p_j\right)\tilde{D}_{jk,j}\right] \\ + \sum_{k\neq j} a_{jk,k}\left(1-2p_j\right)\left(p_j q_j D_{k,k} + D_{jk,jk}\right) + \sum_{k\neq j} a_{jk,jk}\, D_{jk,jk} \tag{B32}$$

with:

$$D_{j,j} = F\left[1 + \sum_{k\neq j} a_{k,k}\, G\, p_k q_k\right] p_j q_j, \tag{B33}$$

$$\tilde{D}_{jk} + \tilde{D}_{j,k} = a_{jk}\left[(1+F)^2 + G\right]\frac{1 + 2\rho_{jk}\sigma}{\rho_{jk}\left(1-\sigma\right)}\, p_j q_j p_k q_k, \tag{B34}$$

$$\tilde{D}_{jk,j} = F\left[a_k + a_{k,k}\left(1-2p_k\right)\right] G\, p_j q_j p_k q_k \\ + F\left(1-2p_j\right)\left[\tilde{D}_{jk} + 2a_{jk}\left[F\left(1+F\right) + G\right] p_j q_j p_k q_k\right], \tag{B35}$$

$$D_{jk,jk} = \left(G + F^2\right) p_j q_j p_k q_k\,. \tag{B36}$$

Equations B31 – B36 yield equations 37 and 41 – 42 in Abu Awad and Roze (2018), expressed to the second order in $U$ (and neglecting the term in $\overline{s}U$ in equation 42).

**Non-Gaussian stabilizing selection.** Expressions for inbreeding depression under the fitness function given by equation 15 in the main text can be obtained from the equations given above, and using the expressions for $a_{\mathbb{U},\mathbb{V}}$ coefficients derived in Supplementary File S1. These expressions show that while the coefficient $a_{jk,jk}$ (representing dominance-by-dominance epistasis on an additive scale) is an order of magnitude smaller than $a_{j,j}$ and $a_{jk}$ under Gaussian stabilizing selection ($Q = 2$), it becomes of the same order of magnitude as $a_{j,j}$, $a_{jk}$ when $Q > 2$: therefore, dominance-by-dominance epistasis is expected to have stronger effects on inbreeding depression when $Q > 2$. To the first order in the strength of selection, and approximating the identity disequilibrium $G_{jk}$ by its expression under free recombination $G$ (see Figure S5 in Abu Awad and Roze, 2018 for justification), we have:

$$\delta \approx -\frac{1}{2}\sum_{j} a_{j,j}\left(1+F\right)p_j q_j - \frac{1}{4}\sum_{j<k} a_{jk,jk}\left[\left(1+F\right)^2 + G\right] p_j q_j p_k q_k \,. \tag{B37}$$

From equation B26, one obtains to the first order in selection coefficients:

$$\begin{aligned}\Delta_{\text{sel}} p_j q_j = \; & a_j\left(1-2p_j\right)\left(1+F\right)p_j q_j + a_{j,j}\left(1-2p_j\right)^2 F\, p_j q_j \\ & + \sum_{k\neq j} a_{jk,k}\left(1-2p_j\right)\left[F\left(1+F\right)+G\right] p_j q_j p_k q_k \\ & + \sum_{k\neq j} a_{jk,jk}\left(1-2p_j\right)^2\left(F^2+G\right) p_j q_j p_k q_k\end{aligned} \tag{B38}$$

which must equal $-u\left(1-2p_j\right)^2$ at mutation-selection balance. Summing equation B38 over $j$, using expressions for selection coefficients derived in Supplementary File S1 (equations A35, A36, A52 and A53), and the fact that $\sum_j r_{\alpha j}^2 p_j q_j = n V_g^0/2$, while $\sum_{j,k}\left(\sum_\alpha r_{\alpha j} r_{\alpha k}\right)^2 p_j q_j p_k q_k = n\left(V_g^0/2\right)^2$, one obtains after simplification (and assuming that the equilibrium at which $p_j = 1/2$ is not stable):

$$\left(\frac{V_g^0}{V_{\text{s}}}\right)^{\frac{Q}{2}} \frac{\Gamma\left(\frac{Q+n}{2}\right)}{\Gamma\left(\frac{n}{2}\right)} = \frac{4U}{Q}\frac{1}{1+3F+\frac{Q-2}{2}\left[F\left(1+3F\right)+3G\right]}. \tag{B39}$$

Using equation B37 and A36, A53 in Supplementary File S1, this yields:

$$\delta = U\frac{1+F+\frac{Q-2}{8}\left[\left(1+F\right)^2+G\right]}{1+3F+\frac{Q-2}{2}\left[F\left(1+3F\right)+3G\right]}\,. \tag{B40}$$

Equation B40 differs from equation 29 in Abu Awad and Roze (2018), that was derived assuming a Gaussian distribution of phenotypic traits at equilibrium; however one can show that both expressions often give very similar quantitative results. Under random mating ($F = G = 0$), equation B40 yields equation 28 in the main text, while when $Q = 2$ (Gaussian fitness function), one recovers the classical expression for inbreeding depression when the dominance coefficient of deleterious alleles is $h = 1/4$, *i.e.* $\delta \approx U/\left(1+\sigma\right)$ (Abu Awad and Roze, 2018).

Although the general expressions given above may be used to compute an expression for inbreeding depression to the second order in $U$ (including the effects of genetic associations between pairs of loci), we only performed this calculation for the case of a randomly mating population. When $\sigma = 0$, equation B26 simplifies to:

$$\Delta_{\text{sel}} p_j q_j \approx a_j \left(1 - 2p_j\right) p_j q_j + \sum_{k \neq j} \left[a_k \left(1 - 2p_j\right) + a_{jk}\right] \tilde{D}_{jk} \,. \tag{B41}$$

When summed over a sufficiently large number of loci, the term in $a_k \left(1 - 2p_j\right)$ should vanish, due to the fact that about half the loci will be at the equilibrium where $p_j < 1/2$, while the other half will be at the symmetric equilibrium with $p_j > 1/2$, so that:

$$\Delta_{\text{sel}} p_j q_j \approx a_j \left(1 - 2p_j\right) p_j q_j + \sum_{k \neq j} a_{jk} \tilde{D}_{jk} \,. \tag{B42}$$

From equation A57 in Supplementary File S1, we have:

$$\sum_j a_j \left(1 - 2p_j\right) p_j q_j \approx -\frac{Q}{4} \left[Z\left(Q,n\right)\left[1 + Z\left(Q,n\right)\right] - Z\left(2Q,n\right)\right] \tag{B43}$$

with:

$$Z\left(Q,n\right) = \left(\frac{V_g^0}{V_\text{s}}\right)^{\frac{Q}{2}} \frac{\Gamma\left(\frac{Q+n}{2}\right)}{\Gamma\left(\frac{n}{2}\right)}, \tag{B44}$$

while from equations A47 and A49 in Supplementary File S1:

$$\sum_{j,k} a_{jk} \tilde{D}_{jk} \approx \frac{4U^2}{n \rho_\text{H}} \tag{B45}$$

at equilibrium, where $\rho_\text{H}$ is the harmonic mean recombination rate between pairs of loci controlling the selected traits. Furthermore, $Z\left(Q,n\right) \approx 4U/Q$ to leading order (equation A41 in Supplementary File S1), so that $\left(V_g^0\right)^{Q/2}$ is of order $U$, and the terms $\left[Z\left(Q,n\right)\right]^2$ and $Z\left(2Q,n\right)$ (proportional to $\left(V_g^0\right)^Q$) are of order $U^2$:

$$\left[Z\left(Q,n\right)\right]^2 \approx \left(\frac{4U}{Q}\right)^2, \quad Z\left(2Q,n\right) \approx \left(\frac{4U}{Q}\right)^2 \left(\frac{\Gamma\left(\frac{n}{2}\right)}{\Gamma\left(\frac{Q+n}{2}\right)}\right)^2 \frac{\Gamma\left(Q+\frac{n}{2}\right)}{\Gamma\left(\frac{n}{2}\right)}. \tag{B46}$$

Using equations B42 – B46, and $\sum_j \left(\Delta_{\text{sel}} p_j q_j\right) = -U$ at equilibrium yields the following expression:

$$\frac{Q}{4}\left[Z\left(Q,n\right) + \left(\frac{4U}{Q}\right)^2 \left[1 - \frac{\Gamma\left(\frac{n}{2}\right)\Gamma\left(Q+\frac{n}{2}\right)}{\left[\Gamma\left(\frac{Q+n}{2}\right)\right]^2}\right]\right] = U\left(1 + \frac{4U}{n\,\rho_{\mathrm{H}}}\right) \tag{B47}$$

that can be solved to express $Z\left(Q,n\right)$ to the second order in $U$:

$$Z\left(Q,n\right) = \frac{4U}{Q}\left[1 + \frac{4U}{n\,\rho_{\mathrm{H}}} + \frac{4U}{Q}\left[\frac{\Gamma\left(\frac{n}{2}\right)\Gamma\left(Q+\frac{n}{2}\right)}{\left[\Gamma\left(\frac{Q+n}{2}\right)\right]^2} - 1\right]\right]. \tag{B48}$$

Equation B48 is equivalent to equation A66 in Abu Awad and Roze (2018) when $Q = 2$. Neglecting linkage in the expression for inbreeding depression under random mating (equation B8), we have:

$$\delta \approx -\frac{1}{2}\sum_j a_{j,j}\, p_j q_j - \frac{1}{4}\sum_{j<k} a_{jk,jk}\, p_j q_j p_k q_k - \sum_{j<k} a_{jk}\, \tilde{D}_{jk}\,. \tag{B49}$$

Using equation A58 in Supplementary File S1 yields:

$$-\frac{1}{2}\sum_j a_{j,j}\, p_j q_j = \frac{Q}{4}\left[Z\left(Q,n\right)\left[1 + Z\left(Q,n\right)\right] - Z\left(2Q,n\right)\right]. \tag{B50}$$

Using B46 and B48, this simplifies to:

$$-\frac{1}{2}\sum_j a_{j,j}\, p_j q_j = U\left(1 + \frac{4U}{n\,\rho_{\mathrm{H}}}\right) \tag{B51}$$

independent of Q. From equation A59 in Supplementary File S1, we have:

$$-\frac{1}{4}\sum_{j<k} a_{jk,jk}\, p_j q_j p_k q_k = \frac{Q}{32}\Big[\left(Q-2\right) Z\left(Q,n\right)\left[1 + Z\left(Q,n\right)\right] - \left(2Q-2\right) Z\left(2Q,n\right)\Big] \tag{B52}$$

which gives, using B46 and B48:

$$-\frac{1}{4}\sum_{j<k} a_{jk,jk}\, p_j q_j p_k q_k = \frac{(Q-2)\,U}{8}\left(1+\frac{4U}{n\,\rho_{\mathrm{H}}}\right) - \frac{U^2}{2}\,\frac{\Gamma\left(\frac{n}{2}\right)\Gamma\left(Q+\frac{n}{2}\right)}{\left[\Gamma\left(\frac{Q+n}{2}\right)\right]^2}. \tag{B53}$$

Altogether, equations B45, B49, B51 and B53 yield the following expression for $\delta$ to the second order in $U$:

$$\delta \approx U\left(1+\frac{2U}{n\,\rho_{\mathrm{H}}}\right) + \frac{(Q-2)\,U}{8}\left(1+\frac{4U}{n\,\rho_{\mathrm{H}}}\right) - \frac{U^2}{2}\,\frac{\Gamma\left(\frac{n}{2}\right)\Gamma\left(Q+\frac{n}{2}\right)}{\left[\Gamma\left(\frac{Q+n}{2}\right)\right]^2}. \tag{B54}$$

It is possible to show that for $Q=2$, equation B54 is equivalent to equations 41 – 42 in Abu Awad and Roze (2018), expressed to the second order in $U$.
