## Supplementary File S3 for "Epistasis, inbreeding depression and the evolution of self-fertilization"

### FILE S3: QLE EXPRESSIONS, NEGLECTING ASSOCIATIONS BETWEEN LOCI AFFECTING FITNESS

We derive here approximations for the change in mean selfing rate during selection and reproduction, neglecting epistasis and genetic associations between loci affecting fitness. As explained in the main text, we assume that loci affecting the selfing rate have additive effects:

$$\sigma = \sum_{i=1}^{\ell_\sigma} \left( \sigma_i^{\mathrm{M}} + \sigma_i^{\mathrm{P}} \right) . \tag{C1}$$

Neglecting associations between loci affecting the selfing rate, the variance in selfing rate in the population can be expressed as:

$$V_\sigma \approx 2 \sum_i \left( \tilde{D}_{ii} + D_{i,i} \right) . \tag{C2}$$

We first express the change in mean selfing rate in terms of genetic associations, and then compute expressions for these associations at QLE.

**Change in $\overline{\sigma}$ in terms of genetic associations.** The change in $\overline{\sigma}$ during reproduction, $\Delta_{\mathrm{r}}\overline{\sigma}$, can be obtained as follows. The change in $\overline{\sigma_i^{\mathrm{M}}}$ is given by:

$$\Delta_{\mathrm{r}} \, \overline{\sigma_i^{\mathrm{M}}} = \overline{\sigma}' \, \mathrm{E}' \left[ \frac{\sigma}{\overline{\sigma}'} \, \frac{\sigma_i^{\mathrm{M}} + \sigma_i^{\mathrm{P}}}{2} \right] + \left( 1 - \overline{\sigma}' \right) \mathrm{E}' \left[ \frac{1 - \sigma}{1 - \overline{\sigma}'} \, \frac{\sigma_i^{\mathrm{M}} + \sigma_i^{\mathrm{P}}}{2} \right] - \overline{\sigma_i^{\mathrm{M}}}' \tag{C3}$$

where $\mathrm{E}'$ stands for the average over all individuals after selection, and $\overline{\sigma}'$, $\overline{\sigma_i^{\mathrm{M}}}'$ the averages of $\sigma$ and $\sigma_i^{\mathrm{M}}$ over all individuals after selection. The first term of equation C3 corresponds to the contribution from selfed offspring to the average after reproduction (each parent contributing to the pool of selfed offspring in proportion $\sigma/\overline{\sigma}'$), while the second term corresponds to the contribution from outcrossed offspring. Using equation

2 in the main text, this becomes:

$$\Delta_{\mathrm{r}}\,\overline{\sigma_i^{\mathrm{M}}} = \mathrm{E}'\left[\sigma\,\frac{\zeta_i^{\mathrm{M}} + \zeta_i^{\mathrm{P}} - \nabla_i'}{2}\right] + \mathrm{E}'\left[(1-\sigma)\,\frac{\zeta_i^{\mathrm{M}} + \zeta_i^{\mathrm{P}} - \nabla_i'}{2}\right] \tag{C4}$$

with $\nabla_i' = \overline{\sigma_i^{\mathrm{M}}}' - \overline{\sigma_i^{\mathrm{P}}}'$. Using the fact that $\mathrm{E}'\left[\zeta_i^{\mathrm{M}}\right] = \mathrm{E}'\left[\zeta_i^{\mathrm{P}}\right] = 0$, equation C4 yields:

$$\Delta_{\mathrm{r}}\,\overline{\sigma_i^{\mathrm{M}}} = -\frac{\nabla_i'}{2}. \tag{C5}$$

Similarly, the change in $\overline{\sigma_i^{\mathrm{P}}}$ is given by:

$$\Delta_{\mathrm{r}}\,\overline{\sigma_i^{\mathrm{P}}} = \overline{\sigma}'\,\mathrm{E}'\left[\frac{\sigma}{\overline{\sigma}'}\,\frac{\sigma_i^{\mathrm{M}} + \sigma_i^{\mathrm{P}}}{2}\right] + (1-\overline{\sigma}')\,\mathrm{E}'\left[\frac{1-\kappa\,\sigma}{1-\kappa\,\overline{\sigma}'}\,\frac{\sigma_i^{\mathrm{M}} + \sigma_i^{\mathrm{P}}}{2}\right] - \overline{\sigma_i^{\mathrm{P}}}' \tag{C6}$$

where again the first term corresponds to the contribution from selfed offspring while the second term corresponds to the contribution from outcrossed offspring, each parent contributing to the pollen pool in proportion $(1-\kappa\,\sigma)\,/\,(1-\kappa\,\overline{\sigma}')$. Equation C6 may be written as:

$$\Delta_{\mathrm{r}}\,\overline{\sigma_i^{\mathrm{P}}} = \mathrm{E}'\left[\sigma\,\frac{\zeta_i^{\mathrm{M}} + \zeta_i^{\mathrm{P}} + \nabla_i'}{2}\right] + \frac{1-\overline{\sigma}'}{1-\kappa\,\overline{\sigma}'}\,\mathrm{E}'\left[(1-\kappa\,\sigma)\,\frac{\zeta_i^{\mathrm{M}} + \zeta_i^{\mathrm{P}} + \nabla_i'}{2}\right]. \tag{C7}$$

Using equations C1 and equation 2 in the main text, the selfing rate of an individual is given by:

$$\sigma = \overline{\sigma}' + \sum_i \left(\zeta_i^{\mathrm{M}} + \zeta_i^{\mathrm{P}}\right). \tag{C8}$$

Replacing $\sigma$ by equation C8, using equations 4 and 5 in the main text, and neglecting associations between different loci affecting the selfing rate, equation C7 becomes:

$$\Delta_{\mathrm{r}}\,\overline{\sigma_i^{\mathrm{P}}} = \frac{1-\kappa}{1-\kappa\,\overline{\sigma}'}\left(\tilde{D}_{ii}' + D_{i,i}'\right) + \frac{\nabla_i'}{2}. \tag{C9}$$

From equations C1, C2, C5 and C9, the change in mean selfing rate during reproduction is:

$$\Delta_{\mathrm{r}}\,\overline{\sigma} = \frac{1-\kappa}{1-\kappa\,\overline{\sigma}'}\,\frac{V_\sigma'}{2} \tag{C10}$$

where $V'_\sigma$ is the variance in selfing rate after selection. $\Delta_{\mathrm{r}}\,\overline{\sigma}$ is positive and represents the effect of the automatic transmission advantage associated with selfing, which vanishes under complete pollen discounting ($\kappa = 1$). Because we assume that the variance in selfing rate in the population is small, and because the change in $\overline{\sigma}$ during selection should be proportional to $V_\sigma$, $\overline{\sigma}'$ can be approximated by $\overline{\sigma}$ in the equation above, yielding:

$$\Delta_{\mathrm{r}}\,\overline{\sigma} \approx \frac{1-\kappa}{1-\kappa\,\overline{\sigma}}\frac{V'_\sigma}{2}. \tag{C11}$$

The change in mean selfing rate during selection is given by:

$$\begin{aligned}\Delta_{\mathrm{s}}\,\overline{\sigma} &= \mathrm{E}\left[\frac{W}{\overline{W}}\sum_i \left(\sigma_i^{\mathrm{M}} + \sigma_i^{\mathrm{P}}\right)\right] - \overline{\sigma} \\ &= \mathrm{E}\left[\frac{W}{\overline{W}}\sum_i \left(\zeta_i^{\mathrm{M}} + \zeta_i^{\mathrm{P}}\right)\right].\end{aligned} \tag{C12}$$

Ignoring epistatic interactions between loci affecting fitness, equation 8 in the main text simplifies to:

$$\frac{W}{\overline{W}} = 1 + \sum_j \left[a_j\left(\zeta_{j,\emptyset} + \zeta_{\emptyset,j}\right) + a_{j,j}\left(\zeta_{j,j} - D_{j,j}\right)\right]. \tag{C13}$$

Using equation C13 and equations 4 and 5 in the main text, this yields:

$$\Delta_{\mathrm{s}}\,\overline{\sigma} = 2\sum_{i,j} a_j\left(\tilde{D}_{ij} + \tilde{D}_{i,j}\right) + 2\sum_{i,j} a_{j,j}\,\tilde{D}_{ij,j} \tag{C14}$$

where the sums are over all loci $i$ affecting the selfing rate of individuals and all loci $j$ affecting fitness. The first term of equation C14 depends on directional selection at loci affecting fitness ($a_j$) and on associations between alleles increasing selfing and alleles 1 at fitness loci, either on the same or on different haplotypes ($\tilde{D}_{ij}$, $\tilde{D}_{i,j}$). As we will see, this term is positive, reflecting the fact that selfing increases the efficiency of selection by increasing homozygosity at fitness loci, so that the favored alleles at these loci tend to be more frequent in genomes coding for higher selfing rates. The second term of equation C14 depends on dominance at fitness loci ($a_{j,j}$) and on associations $\tilde{D}_{ij,j}$ between loci affecting selfing and fitness loci. We will see that these associations are positive, reflecting the fact that alleles coding for higher selfing rates tend to be found in more homozygous backgrounds. The coefficients $a_{j,j}$ are negative when deleterious alleles are partially recessive, in which case the second term of equation C14 represents the effect of inbreeding depression disfavoring selfing.

**Associations generated by selfing.** The genetic associations that appear in equation C14 in turn depend on other associations between pairs of loci. Some of these associations are generated by partial selfing, even in the absence of selection ($W$ constant) and in the absence of genetic variation for the selfing rate. In particular, a recursion for the within-locus association $D_{i,i}$ (under neutrality and in the absence of variation for selfing) is given by:

$$D''_{i,i} = \frac{\overline{\sigma}}{2} \left( \tilde{D}_{ii} + D_{i,i} \right). \tag{C15}$$

Indeed, the covariance between the effects of the two homologous alleles at locus $i$ in an individual equals zero if this individual has been produced by random mating, while if the individual has been produced by selfing (probability $\overline{\sigma}$) its two alleles are the copies of the same parental allele with probability 1/2, and come from the two parental alleles with probability 1/2. At equilibrium, equation C15 yields:

$$D_{i,i} = F\, \tilde{D}_{ii}, \quad \text{with} \quad F = \frac{\overline{\sigma}}{2 - \overline{\sigma}}. \tag{C16}$$

Similarly, $D_{j,j} = F\, \tilde{D}_{jj}$ at equilibrium; however, $\tilde{D}_{jj} = p_j q_j$ in the case of biallelic loci (where $p_j$ and $q_j$ are the frequencies of alleles 1 and 0 at locus $j$), so that:

$$D_{j,j} = F\, p_j q_j. \tag{C17}$$

The association $D_{ij,ij}$ is also generated by selfing, even when loci $i$ and $j$ are neutral. In this case, $D_{ij,ij}$ at the next generation is given by:

$$\begin{aligned} D''_{ij,ij} = \frac{\overline{\sigma}}{2}\Big[&(1-\rho_{ij})^2\left(\tilde{D}_{iijj}+D_{ij,ij}\right) + 2\rho_{ij}\left(1-\rho_{ij}\right)\left(\tilde{D}_{iij,j}+\tilde{D}_{ijj,i}\right) \\ &+ {\rho_{ij}}^2\left(\tilde{D}_{iijj}+D_{ij,ij}\right)\Big] \end{aligned} \tag{C18}$$

where $\rho_{ij}$ is the recombination rate between loci $i$ and $j$. From the fact that locus $j$ is biallelic, repeated $j$ indices that appear in the associations above can be eliminated using the relation (e.g., Barton and Turelli, 1991; Kirkpatrick et al., 2002):

$$D_{\mathbb{S}jj} = p_j q_j D_{\mathbb{S}} + \left(1-2p_j\right) D_{\mathbb{S}j} \tag{C19}$$

where $\mathbb{S}$ is any set of loci either on the same or on different haplotypes from the same individual. Furthermore, associations involving a single $j$ index equal zero at equilibrium when locus $j$ is neutral (since alleles 0 and 1 are then equivalent), so that $\tilde{D}_{iijj} = \tilde{D}_{ii}\, p_j q_j$ and $\tilde{D}_{ijj,i} = D_{i,i}\, p_j q_j$. Finally, one can show using the same reasoning as above that $\tilde{D}_{ijj,i} = F\,\tilde{D}_{ii}\, p_j q_j$ at equilibrium, leading to the following expression for $D_{ij,ij}$ at equilibrium:

$$D_{ij,ij} = \phi_{ij}\,\tilde{D}_{ii}\, p_j q_j, \quad \text{with} \quad \phi_{ij} = \frac{\overline{\sigma}}{2-\overline{\sigma}} \frac{2-\overline{\sigma}-2\left(2-3\overline{\sigma}\right)\rho_{ij}\left(1-\rho_{ij}\right)}{2-\overline{\sigma}\left[1-2\rho_{ij}\left(1-\rho_{ij}\right)\right]} \tag{C20}$$

(e.g., Abu Awad and Roze, 2018), $\phi_{ij}$ representing the joint probability of identity-by-descent at loci $i$ and $j$.

**Associations $\tilde{D}_{ij,j}$.** The association $\tilde{D}_{ij,j}$ is generated by the effect of locus $i$ on the selfing rate, even in the absence of selection at locus $j$. Its equilibrium value can be computed as follows. We have:

$$D''_{ij,j} = \mathrm{E}''\left[\left(\sigma_i^{\mathrm{M}} - \overline{\sigma_i^{\mathrm{M}}}''\right)\left(X_j^{\mathrm{M}} - p_j^{\mathrm{M}\prime\prime}\right)\left(X_j^{\mathrm{P}} - p_j^{\mathrm{P}\prime\prime}\right)\right] \tag{C21}$$

where $\mathrm{E}''$ stands for the average among offpsring (after reproduction), $\overline{\sigma_i^{\mathrm{M}}}''$ is the average of $\sigma_i^{\mathrm{M}}$ among offspring, and $p_j^{\mathrm{M}}{}''$, $p_j^{\mathrm{P}}{}''$ the frequencies of alleles 1 at locus $j$ in maternally and paternally derived genomes of offspring. In the absence of selection ($W$ constant) we have $\Delta_{\mathrm{r}}\,\overline{\sigma_i^{\mathrm{M}}} = \overline{\sigma_i^{\mathrm{M}}}'' - \overline{\sigma_i^{\mathrm{M}}}$, $\Delta_{\mathrm{r}}\,p_j^{\mathrm{M}} = p_j^{\mathrm{M}}{}'' - p_j^{\mathrm{M}}$ and $\Delta_{\mathrm{r}}\,p_j^{\mathrm{P}} = p_j^{\mathrm{P}}{}'' - p_j^{\mathrm{P}}$, and equation C21 can thus be written:

$$D''_{ij,j} = \mathrm{E}''\left[\left(\sigma_i^{\mathrm{M}} - \overline{\sigma_i^{\mathrm{M}}} - \Delta_{\mathrm{r}}\,\overline{\sigma_i^{\mathrm{M}}}\right)\left(X_j^{\mathrm{M}} - p_j^{\mathrm{M}} - \Delta_{\mathrm{r}}\,p_j^{\mathrm{M}}\right)\left(X_j^{\mathrm{P}} - p_j^{\mathrm{P}} - \Delta_{\mathrm{r}}\,p_j^{\mathrm{P}}\right)\right]. \tag{C22}$$

Noting that $\mathrm{E}''\left[\sigma_i^{\mathrm{M}} - \overline{\sigma_i^{\mathrm{M}}}\right] = \Delta_{\mathrm{r}}\,\overline{\sigma_i^{\mathrm{M}}}$, while $\mathrm{E}''\left[X_j^{\mathrm{M}} - p_j^{\mathrm{M}}\right] = \Delta_{\mathrm{r}}\,p_j^{\mathrm{M}}$ and $\mathrm{E}''\left[X_j^{\mathrm{P}} - p_j^{\mathrm{P}}\right] = \Delta_{\mathrm{r}}\,p_j^{\mathrm{P}}$, expanding equation C22 yields:

$$\begin{aligned} D''_{ij,j} = {} & D^{\mathrm{rep}}_{ij,j} - \left(\Delta_{\mathrm{r}}\,\overline{\sigma_i^{\mathrm{M}}}\right) D^{\mathrm{rep}}_{j,j} - \left(\Delta_{\mathrm{r}}\,p_j^{\mathrm{M}}\right) D^{\mathrm{rep}}_{i,j} - \left(\Delta_{\mathrm{r}}\,p_j^{\mathrm{P}}\right) D^{\mathrm{rep}}_{ij,\emptyset} \\ & + 2\left(\Delta_{\mathrm{r}}\,\overline{\sigma_i^{\mathrm{M}}}\right)\left(\Delta_{\mathrm{r}}\,p_j^{\mathrm{M}}\right)\left(\Delta_{\mathrm{r}}\,p_j^{\mathrm{P}}\right) \end{aligned} \tag{C23}$$

where $D^{\mathrm{rep}}_{\mathbb{U},\mathbb{V}}$ stands for associations measured after reproduction, but using "reference values" (the values of $\overline{\sigma_i^{\mathrm{M}}}$, $p_j^{\mathrm{M}}$ and $p_j^{\mathrm{P}}$ that appear in the associations) measured before reproduction: for example, $D^{\mathrm{rep}}_{ij,j} = \mathrm{E}''\left[\left(\sigma_i^{\mathrm{M}} - \overline{\sigma_i^{\mathrm{M}}}\right)\left(X_j^{\mathrm{M}} - p_j^{\mathrm{M}}\right)\left(X_j^{\mathrm{P}} - p_j^{\mathrm{P}}\right)\right]$. Equation C23 can be simplified by noting that $\Delta_{\mathrm{r}}\,p_j^{\mathrm{M}} = \Delta_{\mathrm{r}}\,p_j^{\mathrm{P}} = 0$ when allele $j$ is neutral, yielding:

$$D''_{ij,j} = D^{\mathrm{rep}}_{ij,j} - \left(\Delta_{\mathrm{r}}\,\overline{\sigma_i^{\mathrm{M}}}\right) D^{\mathrm{rep}}_{j,j}. \tag{C24}$$

Furthermore, because $\Delta_{\mathrm{r}}\,\overline{\sigma_i^{\mathrm{M}}}$ is proportional to the amount of genetic variation for selfing (that we suppose small), $D^{\mathrm{rep}}_{j,j}$ can be replaced by its expression in the absence of variation for selfing, given by $\frac{\overline{\sigma}}{2}\left(p_j q_j + D_{j,j}\right)$ (see above). Finally, $D^{\text{rep}}_{ij,j}$ is given by:

$$
\begin{aligned}
D^{\text{rep}}_{ij,j} = \overline{\sigma}\ \mathrm{E}\Bigg[\frac{\sigma}{\overline{\sigma}}\Bigg[&\frac{1-\rho_{ij}}{4}\left(\zeta_{ijj,\emptyset} + \zeta_{ij,j} + \zeta_{j,ij} - \nabla_i\,\zeta_{j,j} + \zeta_{\emptyset,ijj} - \nabla_i\,\zeta_{\emptyset,jj}\right) \\
&+ \frac{\rho_{ij}}{4}\big(\zeta_{ij,j} + \zeta_{i,jj} + \zeta_{jj,i} - \nabla_i\,\zeta_{jj,\emptyset} + \zeta_{j,ij} - \nabla_i\,\zeta_{j,j}\big)\Bigg]\Bigg] \\
+ (1-\overline{\sigma})\ \mathrm{E}\Bigg[&\frac{1-\sigma}{1-\overline{\sigma}}\Bigg[\frac{1-\rho_{ij}}{2}\left(\zeta_{ij,\emptyset} + \zeta_{\emptyset,ij} - \nabla_i\,\zeta_{\emptyset,j}\right) + \frac{\rho_{ij}}{2}\left(\zeta_{i,j} + \zeta_{j,i} - \nabla_i\,\zeta_{j,\emptyset}\right)\Bigg]\Bigg] \\
&\times \mathrm{E}\left[\frac{1-\kappa\,\sigma}{1-\kappa\,\overline{\sigma}}\,\frac{\zeta_{j,\emptyset} + \zeta_{\emptyset,j}}{2}\right].
\end{aligned}
\tag{C25}
$$

The first term of equation C25 corresponds to the contribution of individuals produced by selfing to $D^{\text{rep}}_{ij,j}$. In this case, the three genes may have been present in different configurations in the parent with different probabilities (corresponding to the different $\zeta_{\mathbb{U},\mathbb{V}}$ terms). The $\nabla_i$ terms stem from the fact that $\overline{\sigma^{\text{M}}_i} \neq \overline{\sigma^{\text{P}}_i}$; note that $p^{\text{M}}_j = p^{\text{P}}_j$ under the assumption that locus $j$ is neutral, so that $\nabla_j$ terms do not arise. The second term of equation C25 corresponds to the contribution of individuals produced by outcrossing: the first average is over all maternal individuals (transmitting genes at loci $i$ and $j$), while the second average is over all paternal individuals (transmitting one gene at locus $j$). Using equation C8, neglecting genetic associations between different loci affecting selfing and assuming that the genetic variance in selfing rate is small (so that products between associations involving a single $i$ index may be neglected), one arrives at:

$$
\begin{aligned}
D^{\text{rep}}_{ij,j} \approx \frac{\overline{\sigma}}{2}&\left[(1-\rho_{ij})\,\tilde{D}_{ijj} + \rho_{ij}\,\tilde{D}_{i,jj} + \tilde{D}_{ij,j} - \frac{\nabla_i}{2}\left(p_j q_j + D_{j,j}\right)\right] \\
&+ \frac{1}{2}\left[(1-\rho_{ij})\,\tilde{D}_{iijj} + \rho_{ij}\,\tilde{D}_{ii,jj} + \tilde{D}_{ijj,i} + \tilde{D}_{iij,j} + D_{ij,ij}\right]
\end{aligned}
\tag{C26}
$$

(note that the second term of equation C25 cancels under the assumption that locus $j$ is neutral). Using equation C19 and the fact that associations with a single $j$ index equal zero when locus $j$ is neutral (in particular, $\tilde{D}_{ij} = \tilde{D}_{i,j} = 0$), we have $\tilde{D}_{ijj} = \tilde{D}_{i,jj} = 0$ while $\tilde{D}_{iijj} = \tilde{D}_{ii,jj} = \tilde{D}_{ii}\, p_j q_j$. Using equations C5, C16, C17, C20 and C24, one finally obtains:

$$D''_{ij,j} \approx \frac{\overline{\sigma}}{2}\, \tilde{D}_{ij,j} + \frac{1}{2}\left[(1+F)^2 + G_{ij}\right] \tilde{D}_{ii}\, p_j q_j \tag{C27}$$

where $G_{ij} = \phi_{ij} - F^2$ is the identity disequilibrium between loci $i$ and $j$ (e.g., Weir and Cockerham, 1973).

The recursion for $D_{j,ij}$ is obtained using the same general method. We have in particular:

$$D''_{j,ij} = D^{\text{rep}}_{j,ij} - \left(\Delta_{\text{r}}\, \overline{\sigma^{\text{P}}_i}\right) D^{\text{rep}}_{j,j}, \tag{C28}$$

while:

$$D^{\text{rep}}_{j,ij} \approx \frac{\overline{\sigma}}{2}\left[\tilde{D}_{ij,j} + \frac{\nabla_i}{2}\left(p_j q_j + D_{j,j}\right)\right] + \frac{1}{2}\left[(1+F)^2 + G_{ij}\right] \tilde{D}_{ii}\, p_j q_j. \tag{C29}$$

Equations C9, C28 and C29 yield:

$$\begin{aligned} D''_{j,ij} \approx\ & \frac{\overline{\sigma}}{2}\, \tilde{D}_{ij,j} + \frac{1}{2}\left[(1+F)^2 + G_{ij}\right] \tilde{D}_{ii}\, p_j q_j \\ & - \frac{\overline{\sigma}}{2}\, \frac{1-\kappa}{1-\kappa\,\overline{\sigma}}\, (1+F)^2\, \tilde{D}_{ii}\, p_j q_j. \end{aligned} \tag{C30}$$

Finally, equations C27 and C30 yield the following expression for $\tilde{D}_{ij,j}$ at QLE:

$$\tilde{D}_{ij,j} \approx \frac{1}{2-\overline{\sigma}}\left[(1+F)^2\left(1 - \frac{\overline{\sigma}}{2}\, \frac{1-\kappa}{1-\kappa\,\overline{\sigma}}\right) + G_{ij}\right] \tilde{D}_{ii}\, p_j q_j. \tag{C31}$$

Equation C31 is equivalent to the result obtained by Epinat and Lenormand (2009) under strong discounting ($\kappa \approx 1$, their equation A5). Under complete outcrossing, it simplifies to $\tilde{D}_{ij,j} \approx (1/2)\, \tilde{D}_{ii}\, p_j q_j$.

**Associations $\tilde{D}_{ij}$ and $\tilde{D}_{i,j}$.** The associations $\tilde{D}_{ij}$, $\tilde{D}_{i,j}$ that appear in the first term of equation C14 are generated by selection acting at locus $j$ and by the effect of locus

$i$ on the selfing rate, and can be computed as follows. Using the same reasoning as for the derivation of equation C24, one obtains:

$$D''_{ij,\emptyset} = D^{\text{rep}}_{ij,\emptyset} - \left(\Delta_{\text{r}}\, \overline{\sigma^{\text{M}}_i}\right)\left(\Delta_{\text{r}}\, p^{\text{M}}_j\right) \approx D^{\text{rep}}_{ij,\emptyset}. \tag{C32}$$

Indeed, $\Delta_{\text{r}}\, \overline{\sigma^{\text{M}}_i}$ and $\Delta_{\text{r}}\, p^{\text{M}}_j$ are both proportional to the variance in selfing rate, and their product can thus be neglected. $D^{\text{rep}}_{ij,\emptyset}$ is then given by:

$$D^{\text{rep}}_{ij,\emptyset} = (1-\rho_{ij})\, \tilde{D}'_{ij} + \rho_{ij}\, \tilde{D}'_{i,j}. \tag{C33}$$

Similarly,

$$D''_{\emptyset,ij} \approx D^{\text{rep}}_{\emptyset,ij} \tag{C34}$$

while $D^{\text{rep}}_{\emptyset,ij}$ is given by:

$$\begin{aligned} D^{\text{rep}}_{\emptyset,ij} = \overline{\sigma}'\, \text{E}\Bigg[\frac{\sigma}{\overline{\sigma}'}\Bigg[&\frac{1-\rho_{ij}}{2}\left(\zeta_{ij,\emptyset} + \nabla'_i\, \zeta_{j,\emptyset} + \nabla'_j\, \zeta_{i,\emptyset} + \nabla'_i \nabla'_j + \zeta_{\emptyset,ij}\right) \\ &+ \frac{\rho_{ij}}{2}\left(\zeta_{i,j} + \nabla'_i\, \zeta_{\emptyset,j} + \zeta_{j,i} + \nabla'_j\, \zeta_{\emptyset,i}\right)\Bigg]\Bigg] \\ + (1-\overline{\sigma}')\, \text{E}\Bigg[\frac{1-\kappa\,\sigma}{1-\kappa\,\overline{\sigma}'}\Bigg[&\frac{1-\rho_{ij}}{2}\left(\zeta_{ij,\emptyset} + \nabla'_i\, \zeta_{j,\emptyset} + \nabla'_j\, \zeta_{i,\emptyset} + \nabla'_i \nabla'_j + \zeta_{\emptyset,ij}\right) \\ &+ \frac{\rho_{ij}}{2}\left(\zeta_{i,j} + \nabla'_i\, \zeta_{\emptyset,j} + \zeta_{j,i} + \nabla'_j\, \zeta_{\emptyset,i}\right)\Bigg]\Bigg] \end{aligned} \tag{C35}$$

where the first term is the contribution of individuals produced by selfing while the second term is the contribution of individuals produced by outcrossing, and $\nabla'_j = p^{\text{M}\prime}_j - p^{\text{P}\prime}_j$. The terms in $\nabla'_i$, $\nabla'_j$ are proportional to the variance in selfing rate and generate terms that are proportional to the square of the variance, and are thus neglected. One obtains:

$$\begin{aligned} D^{\text{rep}}_{\emptyset,ij} = &(1-\rho_{ij})\, \tilde{D}'_{ij} + \rho_{ij}\, \tilde{D}'_{i,j} \\ &+ \frac{1-\kappa}{1-\kappa\,\overline{\sigma}}\left[(1-\rho_{ij})\, \tilde{D}'_{iij} + \rho_{ij}\, \tilde{D}'_{ii,j} + \tilde{D}'_{ij,i}\right]. \end{aligned} \tag{C36}$$

It is possible to show that $\tilde{D}'_{iij}$ and $\tilde{D}'_{ii,j}$ are negligible compared to $\tilde{D}'_{ij,i}$ (using the same method as for deriving $\tilde{D}_{ij,i}$ below), and equations C32, C33, C34 and C36 thus yield:

$$\tilde{D}''_{ij} = (1 - \rho_{ij})\, \tilde{D}'_{ij} + \rho_{ij}\, \tilde{D}'_{i,j} + \frac{1-\kappa}{2\,(1-\kappa\,\overline{\sigma})}\, \tilde{D}'_{ij,i}. \tag{C37}$$

Similarly, one arrives at:

$$\tilde{D}''_{i,j} = \frac{\overline{\sigma}}{2} \left( \tilde{D}'_{ij} + \tilde{D}'_{i,j} \right) + \tilde{D}'_{ij,i}. \tag{C38}$$

Using the same reasoning as for deriving equation C32, $D'_{ij,\emptyset}$ is given by:

$$D'_{ij,\emptyset} = D^{\mathrm{sel}}_{ij,\emptyset} - \left( \Delta_{\mathrm{s}}\, \overline{\sigma_i^{\mathrm{M}}} \right) \left( \Delta_{\mathrm{s}}\, p_j^{\mathrm{M}} \right) \tag{C39}$$

where $D^{\mathrm{sel}}_{\mathbb{U},\mathbb{V}}$ refers to associations measured after selection, but using as reference values $\overline{\sigma_i^{\mathrm{M}}}$, $\overline{\sigma_i^{\mathrm{P}}}$, $p_j^{\mathrm{M}}$ and $p_j^{\mathrm{P}}$ before selection — in particular, $D^{\mathrm{sel}}_{ij,\emptyset} = \mathrm{E}' \left[ \left( \sigma_i^{\mathrm{M}} - \overline{\sigma_i^{\mathrm{M}}} \right) \left( X_j^{\mathrm{M}} - p_j^{\mathrm{M}} \right) \right]$ — and where $\Delta_{\mathrm{s}}\, \overline{\sigma_i^{\mathrm{M}}}$ and $\Delta_{\mathrm{s}}\, p_j^{\mathrm{M}}$ are the changes in $\overline{\sigma_i^{\mathrm{M}}}$ and in $p_j^{\mathrm{M}}$ due to selection. In the following, we compute expressions to the first order in the strength of selection (measured as $\epsilon$, and representing the largest of $a_{\mathbb{U},\mathbb{V}}$ coefficients in absolute value), and thus neglect the last product of equation C39 (which is of order $\epsilon^2$). We thus have:

$$D'_{ij,\emptyset} \approx D^{\mathrm{sel}}_{ij,\emptyset} = \mathrm{E}\left[ \frac{W}{\overline{W}}\, \zeta_{ij,\emptyset} \right] \tag{C40}$$

where the last average is over all individuals before selection. Using equation C13, and neglecting associations involving different fitness loci, this yields:

$$D'_{ij,\emptyset} \approx D_{ij,\emptyset} + a_j \left( D_{ijj,\emptyset} + D_{ij,j} \right) + a_{j,j} \left( D_{ijj,j} - D_{ij,\emptyset} D_{j,j} \right). \tag{C41}$$

Eliminating repeated $j$ indices from associations, we have $D_{ijj,\emptyset} = (1 - 2p_j)\, D_{ij,\emptyset}$, and $D_{ijj,j} = p_j q_j\, D_{i,j} + (1 - 2p_j)\, D_{ij,j}$. Because $D_{ij,\emptyset}$ and $D_{i,j}$ are of order $\epsilon$, equation C41 simplifies to:

$$D'_{ij,\emptyset} \approx D_{ij,\emptyset} + \left[ a_j + a_{j,j}\, (1 - 2p_j) \right] D_{ij,j} + o(\epsilon). \tag{C42}$$

However, using equation C42 (and the equivalent expression for $D'_{i,j}$) yields QLE expressions that diverge when $\rho_{ij}\,(1-\overline{\sigma})$ tends to zero (that is, either when $\rho_{ij}$ tends to 0 or when $\overline{\sigma}$ tends to 1). Indeed, $D_{ij,\emptyset}$ and $D_{i,j}$ are of order 1 when $\rho_{ij}\,(1-\overline{\sigma})$ is of order $\epsilon$, and terms such as $a_j\,D_{ij,\emptyset}$, $a_{j,j}\,D_{i,j}$ become of order $\epsilon$, and must be retained in order to prevent divergence at low effective recombination (e.g., Charlesworth, 1990; Roze, 2014; Gervais and Roze, 2017) — note that this implies that allele frequencies at fitness loci are at an equilibrium, in order for the QLE approximation to hold. In that case, and in order to simplify expressions, we will assume that polymorphism stays low at fitness loci and derive expressions to the first order in $p_j q_j$ ("rare alleles approximation"). Given that associations involving one or several $j$ indices are proportional to $p_j q_j$, this yields:

$$D'_{ij,\emptyset} \approx \left[1 + a_j\,(1-2p_j)\right] D_{ij,\emptyset} + \left[a_j + a_{j,j}\,(1-2p_j)\right] D_{ij,j} + o(\epsilon)\,. \tag{C43}$$

Similarly, one obtains:

$$D'_{i,j} \approx \left[1 + a_j\,(1-2p_j)\right] D_{i,j} + \left[a_j + a_{j,j}\,(1-2p_j)\right] D_{ij,j} + o(\epsilon) \tag{C44}$$

and finally

$$\tilde{D}'_{ij} \approx \left[1 + a_j\,(1-2p_j)\right] \tilde{D}_{ij} + \left[a_j + a_{j,j}\,(1-2p_j)\right] \tilde{D}_{ij,j} + o(\epsilon) \tag{C45}$$

$$\tilde{D}'_{i,j} \approx \left[1 + a_j\,(1-2p_j)\right] \tilde{D}_{i,j} + \left[a_j + a_{j,j}\,(1-2p_j)\right] \tilde{D}_{ij,j} + o(\epsilon)\,. \tag{C46}$$

Equations C37, C38, C45 and C46 finally yield:

$$\tilde{D}_{ij} + \tilde{D}_{i,j} \approx \frac{\left[a_j + a_{j,j}\,(1-2p_j)\right](1+2\rho_{ij}\overline{\sigma})\,\tilde{D}_{ij,j} + X\,\tilde{D}'_{ij,i}}{\rho_{ij}\,(1-\overline{\sigma}) - a_j\,(1-2p_j)\left[1-\frac{\overline{\sigma}}{2}-\rho_{ij}\,(1-2\overline{\sigma})\right]} \tag{C47}$$

with:

$$X = \frac{1-\kappa}{2\,(1-\kappa\,\overline{\sigma})} + 2\rho_{ij} - a_j\,(1-2p_j)\,(1-2\rho_{ij})\,. \tag{C48}$$

Associations $\tilde{D}_{ij}$ and $\tilde{D}_{i,j}$ thus depend on $\tilde{D}_{ij,j}$ (given by equation C31) and on $\tilde{D}'_{ij,i}$, whose expression at QLE must be computed to the first order in the strength of selection $\epsilon$. The effect of genetic variance for the selfing rate can be ignored when computing the effect of reproduction on $\tilde{D}_{ij,i}$, as it would generate second-order terms in $V_\sigma$. This yields:

$$\tilde{D}''_{ij,i} = \tilde{D}^{\mathrm{rep}}_{ij,i} = \frac{\overline{\sigma}}{2}\left[(1-\rho_{ij})\,\tilde{D}'_{iij} + \rho_{ij}\,\tilde{D}'_{ii,j} + \tilde{D}'_{ij,i}\right]. \tag{C49}$$

As mentioned above, $\tilde{D}'_{iij}$ and $\tilde{D}'_{ii,j}$ are found to equal zero when computed under the same assumptions as for $\tilde{D}_{ij,i}$, and are thus neglected. We then have:

$$\tilde{D}'_{ij,i} = \tilde{D}^{\mathrm{sel}}_{ij,i} - (\Delta_{\mathrm{s}}\, p_j)\, D_{i,i} + o(\epsilon) \tag{C50}$$

with (ignoring associations between different loci affecting fitness):

$$\Delta_{\mathrm{s}}\, p_j = a_j\, p_j q_j + [a_j + a_{j,j}\,(1-2p_j)]\, D_{j,j} + o(\epsilon) \tag{C51}$$

$$\tilde{D}^{\mathrm{sel}}_{ij,i} = \tilde{D}_{ij,i} + a_j\, D_{i,i}\, p_j q_j + [a_j + a_{j,j}\,(1-2p_j)]\, D_{ij,ij} + o(\epsilon)\,. \tag{C52}$$

Therefore:

$$\tilde{D}'_{ij,i} = \tilde{D}_{ij,i} + [a_j + a_{j,j}\,(1-2p_j)]\, G_{ij}\, \tilde{D}_{ii}\, p_j q_j + o(\epsilon) \tag{C53}$$

where again $G_{ij} = \phi_{ij} - F^2$ is the identity disequilibrium between loci $i$ and $j$. Equations C49 and C53 yield, at QLE:

$$\tilde{D}_{ij,i} = [a_j + a_{j,j}\,(1-2p_j)]\, F\, G_{ij}\, \tilde{D}_{ii}\, p_j q_j + o(\epsilon) \tag{C54}$$

which is equivalent to equation A37 in Roze (2015). Using equation C53, we thus have:

$$\tilde{D}'_{ij,i} = [a_j + a_{j,j}\,(1-2p_j)]\,(1+F)\, G_{ij}\, \tilde{D}_{ii}\, p_j q_j + o(\epsilon)\,. \tag{C55}$$

Equation C47 shows that associations $\tilde{D}_{ij}$, $\tilde{D}_{i,j}$ are generated by two different effects. The first involves the association $\tilde{D}_{ij,j}$, representing the fact that alleles increasing selfing at locus $i$ tend to be associated with more homozygous backgrounds at locus $j$. As a consequence, selection at locus $j$ is more efficient among individuals carrying alleles that increase selfing, causing an increased frequency of the best allele at locus $j$ in these individuals. The second effect involves the association $\tilde{D}'_{ij,i}$ and is generated by the identity disequilibrium $G_{ij}$ between loci $i$ and $j$ (equation C55). Identity disequilibrium results from partial selfing, and represents the fact that homozygotes at locus $i$ tend to be also homozygous at locus $j$. A consequence of this identity disequilibrium is that, because selection at locus $j$ is more efficient among homozygotes at this locus, the frequency of the best allele at locus $j$ tends to be higher among homozygotes at locus $i$ than among heterozygotes, which is represented by the association $\tilde{D}_{ij,i}$. In the presence of direct selection at locus $i$ (that is, when $\kappa < 1$), the fact that selection at locus $i$ is more efficient among homozygotes than among heterozygotes, and the fact that homozygotes at locus $i$ tend to be associated with the best allele at locus $j$ generate a positive association between the alleles at both loci that are favored by direct selection. This corresponds to the previously described result that partial selfing generates positive associations between selected loci, even in the absence of epistasis among those loci (Roze and Lenormand, 2005; Kamran-Disfani and Agrawal, 2014). However, equation C48 indicates that the effect of $\tilde{D}_{ij,i}$ persists even in the absence of direct selection at locus $i$ ($\kappa = 1$). This effect can be understood as follows. For simplicity, we will consider the case where only two alleles segregate at locus $i$, an allele $S$ coding for more selfing, and an allele $O$ coding for more outcrossing. Furthermore, we will call $A$ the favored allele at locus $j$, and $a$ the deleterious allele. $\tilde{D}_{ij,i}$ indicates that $A$ tends to be more frequent in $SS$ and $OO$ individuals, while $a$ tends to be more frequent in $OS$ individuals. The population thus contains an excess of $SA/S\cdot$, $OA/O\cdot$, $Sa/O\cdot$ and $Oa/S\cdot$ genotypes, where the dot $\cdot$ stands for any allele at locus $j$, and where the slash symbol separates maternal and paternal haplotypes. The association between loci present on the same haplotype ($\tilde{D}_{ij}$) is not affected by whether individuals have been produced by selfing or outcrossing. However, the association between genes on different haplotypes ($\tilde{D}_{i,j}$) depends on how offspring are produced: in particular, selfing from $SA/S\cdot$ individuals maintains an association between $S$ and $A$ on different haplotypes, while this association is lost when offspring are produced by outcrossing. Similarly, selfing from $OA/O\cdot$ individuals maintains an association between $O$ and $A$ on different haplotypes. This effect is stronger in the case of $SA/S\cdot$ individuals, however, due to their higher selfing rate: as a consequence, the net effect of this process is to generate an association between $S$ and $A$ on different haplotypes ($\tilde{D}_{i,j}$), which can then be converted into an association between genes on the same haplotype ($\tilde{D}_{ij}$) by recombination.

Because $\tilde{D}'_{ij,i} = 0$ while $\tilde{D}_{ij,j} = (1/2)\,\tilde{D}_{ii}\,p_j q_j$ in the absence of selfing ($\overline{\sigma} = 0$), the expressions for $\tilde{D}_{ij}$ and $\tilde{D}_{i,j}$ simplify to:

$$\tilde{D}_{ij} \approx \frac{1}{2}\frac{a_j + a_{j,j}\,(1-2p_j)}{\rho_{ij} - a_j\,(1-2p_j)\,(1-\rho_{ij})}\tilde{D}_{ii}\,p_j q_j, \quad \tilde{D}_{i,j} = 0. \tag{C56}$$

**Variance in selfing rate after selection.** In order to complete the derivations, we can note that equation C11 involves the variance in selfing rate measured after selection $V'_\sigma = 2\left(\tilde{D}'_{ii} + D'_{i,i}\right)$, which must be expressed in terms of $V_\sigma$ before selection. From the QLE expressions given above, the term in $a_{j,j}\,\tilde{D}_{ij,j}$ of equation C14 is proportional to $a_{j,j}$, while the term in $a_j\left(\tilde{D}_{ij}+\tilde{D}_{i,j}\right)$ is proportional to ${a_j}^2$. For consistency, we thus express $V'_\sigma$ to the first order in $a_{j,j}$, and to the second order in $a_j$: this can be achieved by supposing that $a_j$ is of order $\eta$ (where $\eta$ is a small term) while $a_{j,j}$ is of order $\eta^2$. We have:

$$\tilde{D}'_{ii}=\tilde{D}^{\mathrm{sel}}_{ii}-\frac{1}{2}\left[\left(\Delta_{\mathrm{s}}\overline{\sigma_i^{\mathrm{M}}}\right)^2+\left(\Delta_{\mathrm{s}}\overline{\sigma_i^{\mathrm{P}}}\right)^2\right] \tag{C57}$$

$$\tilde{D}'_{i,i}=\tilde{D}^{\mathrm{sel}}_{i,i}-\left(\Delta_{\mathrm{s}}\overline{\sigma_i^{\mathrm{M}}}\right)\left(\Delta_{\mathrm{s}}\overline{\sigma_i^{\mathrm{P}}}\right). \tag{C58}$$

Because $\Delta_{\mathrm{s}}\overline{\sigma_i^{\mathrm{M}}}$ and $\Delta_{\mathrm{s}}\overline{\sigma_i^{\mathrm{P}}}$ are proportional to the genetic variance in selfing rate, their products can be neglected, yielding:

$$\tilde{D}'_{ii}\approx\tilde{D}^{\mathrm{sel}}_{ii},\quad \tilde{D}'_{i,i}\approx\tilde{D}^{\mathrm{sel}}_{i,i}. \tag{C59}$$

We then have (from equation C13):

$$\tilde{D}^{\mathrm{sel}}_{ii}=\tilde{D}_{ii}+\sum_j\left[a_j\left(\tilde{D}_{iij}+\tilde{D}_{ii,j}\right)+a_{j,j}\left(\tilde{D}_{iij,j}-\tilde{D}_{ii}\,D_{j,j}\right)\right]. \tag{C60}$$

As mentioned above, the associations $\tilde{D}_{iij}$ and $\tilde{D}_{ii,j}$ equal zero at QLE, to the first order in selection coefficients, while $\tilde{D}_{iij,j}$ and $\tilde{D}_{ii}\,D_{j,j}$ both equal $F\,\tilde{D}_{ii}\,p_jq_j$ in the absence of selection; therefore:

$$\tilde{D}'_{ii}=\tilde{D}_{ii}+o\left(\eta^2\right). \tag{C61}$$

Then,

$$\tilde{D}^{\mathrm{sel}}_{i,i}=\tilde{D}_{i,i}+\sum_j\left[2a_j\,\tilde{D}_{ij,i}+a_{j,j}\left(D_{ij,ij}-D_{i,i}\,D_{j,j}\right)\right] \tag{C62}$$

finally leading to:

$$\tilde{D}'_{ii}+D'_{i,i}\approx\tilde{D}_{ii}+D_{i,i}+\sum_j\left[\left(2a_j^2\,F+a_{j,j}\right)G_{ij}\,p_jq_j\right]\tilde{D}_{ii} \tag{C63}$$

and thus:

$$V'_\sigma \approx V_\sigma + 2 \sum_{i,j} \left[ \left( 2{a_j}^2\, F + a_{j,j} \right) G_{ij}\, p_j q_j \right] \tilde{D}_{ii}. \tag{C64}$$

Equation C64 stems from the fact that homozygosity at loci coding for the selfing rate is affected by identity disequilibria between these loci and loci affecting fitness; when $a_j$ and $a_{j,j}$ are of the same order of magnitude, the term in ${a_j}^2$ in equation C64 becomes negligible, and the result becomes equivalent to equation 5 in Roze (2015). Because $F = G_{ij} = 0$ when $\overline{\sigma} = 0$, equation C64 simplifies to $V'_\sigma \approx V_\sigma$ when the mean selfing rate of the population approaches zero.

**Expressing $\Delta_{\text{purge}}\overline{\sigma}$ in terms of the increase in mean fitness following a single generation of selfing.** From the results given above, the contribution of purging to the strength of selection for selfing in a randomly mating population ($\Delta_{\text{purge}}\overline{\sigma} = 2 \sum_{i,j} a_j\, \tilde{D}_{ij}$) may be expressed in terms of the effect of a single generation of selfing on the mean fitness of offspring. Imagine a parental, non-inbred population from which a pool of selfed offspring and a pool of outcrossed offspring are produced. Inbreeding depression represents the difference in mean fitness between those two pools of offspring. Imagine now that these individuals are let to reproduce in proportion to their fitness, by random mating within each pool. We will call $\overline{W}_{\text{F2, self}}$ and $\overline{W}_{\text{F2, out}}$ the mean fitnesses of the offspring of selfed and outcrossed individuals, respectively. Neglecting epistasis, the fitness of an individual relative to the mean fitness of the parental population can be expressed as (from equation A1 in Supplementary File S1):

$$\frac{W}{\overline{W}} = 1 + \sum_j a_j \left( \zeta_{j,\emptyset} + \zeta_{\emptyset,j} \right) + \sum_j a_{j,j}\, \zeta_{j,j} \tag{C65}$$

where the allele frequencies $p_j$ within $\zeta_{j,\emptyset}$, $\zeta_{\emptyset,j}$ and $\zeta_{j,j}$ are allele frequencies in the parental population. Averaging $\zeta_{j,\emptyset} + \zeta_{\emptyset,j}$ over the offspring of selfed individuals yields $2\Delta_{\text{self}} p_j = 2\left(p_j^{\text{F2, self}} - p_j\right)$, where $p_j^{\text{F2, self}}$ is the frequency of allele 1 at locus $j$ among the offspring of selfed individuals. Given that reproduction among selfed individuals occurs by random mating, one can show that the average of $\zeta_{i,i}$ among their offspring is given by $(\Delta_{\text{self}} p_j)^2$. Neglecting terms in $(\Delta_{\text{self}} p_j)^2$, we thus have:

$$\frac{\overline{W}_{\text{F2, self}}}{\overline{W}} \approx 1 + 2 \sum_j a_j \, \Delta_{\text{self}} p_j \tag{C66}$$

and similarly

$$\frac{\overline{W}_{\text{F2, out}}}{\overline{W}} \approx 1 + 2 \sum_j a_j \, \Delta_{\text{out}} p_j. \tag{C67}$$

The gain in mean fitness caused by one generation of purging through selfing can thus be expressed as:

$$P = \frac{\overline{W}_{\text{F2, self}}}{\overline{W}} - \frac{\overline{W}_{\text{F2, out}}}{\overline{W}} \approx 2 \sum_j a_j \left(\Delta_{\text{self}} p_j - \Delta_{\text{out}} p_j\right). \tag{C68}$$

The change in $p_j$ due to selection among selfed individuals is given by:

$$\Delta_{\text{self}} p_j = \text{E}\left[\frac{W}{\overline{W}_{\text{self}}} \frac{\zeta_{j,\emptyset} + \zeta_{\emptyset,j}}{2}\right] \tag{C69}$$

where the average is over all selfed individuals, and where $\overline{W}_{\text{self}}$ is the mean fitness of these individuals. From equation C65, we have (given that $D_{j,j}$ among selfed offspring equals $p_j q_j / 2$):

$$\frac{\overline{W}_{\text{self}}}{\overline{W}} = 1 + \frac{1}{2} \sum_j a_{j,j} \, p_j q_j \tag{C70}$$

and thus:

$$\frac{W}{\overline{W}_{\text{self}}} = \frac{W/\overline{W}}{\overline{W}_{\text{self}}/\overline{W}} = 1 + \sum_j a_j \left(\zeta_{j,\emptyset} + \zeta_{\emptyset,j}\right) + \sum_j a_{j,j} \left(\zeta_{j,j} - \frac{1}{2} p_j q_j\right) \tag{C71}$$

to the first order in $a_{\mathbb{U},\mathbb{V}}$ coefficients. From equations C69 and C71, and neglecting associations between loci, one obtains:

$$\Delta_{\text{self}} p_j \approx \frac{1}{2} \left[3a_j + a_{j,j} \left(1 - 2p_j\right)\right] p_j q_j. \tag{C72}$$

Similarly, we have:

$$\Delta_{\text{out}} p_j = \mathrm{E}\left[\frac{W}{\overline{\overline{W}}}\,\frac{\zeta_{j,\emptyset}+\zeta_{\emptyset,j}}{2}\right] \approx a_j\,p_j q_j \tag{C73}$$

and equation C68 thus leads to:

$$P \approx \sum_j a_j\left[a_j + a_{j,j}\left(1-2p_j\right)\right] p_j q_j. \tag{C74}$$
