## Supplementary File S4 for "Epistasis, inbreeding depression and the evolution of self-fertilization"

### FILE S4: QLE EXPRESSIONS, INCLUDING THE EFFECTS OF EPISTASIS

Derivations including the effects of epistasis between pairs of selected loci can be found in a *Mathematica* notebook available as supplementary material. Although it is possible to obtain general expressions valid for arbitrary selfing rate, these expressions quickly become cumbersome when $\overline{\sigma} > 0$ and we thus only present results relative to the initial spread of selfing in an outcrossing population ($\overline{\sigma} \approx 0$). In that case, the change in mean selfing rate per generation can still be decomposed in a change during selection $\Delta_{\mathrm{s}}\overline{\sigma}$ and a change during reproduction $\Delta_{\mathrm{r}}\overline{\sigma}$, with:

$$\Delta_{\mathrm{r}}\,\overline{\sigma} \approx \frac{1-\kappa}{2}\,V_{\sigma} \tag{D1}$$

representing direct selection for selfing (automatic transmission advantage), while (from equation A1 in Supplementary File S1):

$$\begin{aligned}\Delta_{\mathrm{s}}\,\overline{\sigma} = {} & 2\sum_{i,j} a_j\left(\tilde{D}_{ij}+\tilde{D}_{i,j}\right) + 2\sum_{i,j} a_{j,j}\,\tilde{D}_{ij,j} \\ & + 2\sum_{i,j<k} a_{jk}\left(\tilde{D}_{ijk}+\tilde{D}_{i,jk}\right) + 2\sum_{i,j<k} a_{j,k}\left(\tilde{D}_{ij,k}+\tilde{D}_{ik,j}\right) \\ & + 2\sum_{i,j<k} a_{jk,j}\left(\tilde{D}_{ijk,j}+\tilde{D}_{ij,jk}\right) + 2\sum_{i,j<k} a_{jk,k}\left(\tilde{D}_{ijk,k}+\tilde{D}_{ik,jk}\right) \\ & + 2\sum_{i,j<k} a_{jk,jk}\,\tilde{D}_{ijk,jk}\,.\end{aligned} \tag{D2}$$

Leading-order expressions fort the different genetic associations that appear in equation D2 can be obtained using the same methods as in Supplementary File S3 (see also *Mathematica* notebook). The association $\tilde{D}_{ijk,jk}$ is generated by the effect of locus $i$ on the selfing rate, even in the absence of selection at loci $j$ and $k$ (representing the fact that alleles increasing the selfing rate at locus $i$ tend to be more often present in individuals which are homozygous at loci $j$ and $k$). To leading order, it is given by:

$$\tilde{D}_{ijk,jk} \approx \frac{1}{2}\left[1 - 2\rho_{jk}\left(1 - \rho_{jk}\right)\right] \tilde{D}_{ii}\, pq_{jk} \tag{D3}$$

where $pq_{jk} = p_j q_j p_k q_k$. Associations $\tilde{D}_{ijk,j}$ and $\tilde{D}_{ij,jk}$ are given by:

$$\tilde{D}_{ijk,j} \approx \tilde{D}_{ij,jk} \approx \frac{1}{2}\left(1 - 2p_j\right)\left[\left(1 - \rho_{jk}\right)\tilde{D}'_{jk} + \rho_{jk}\,\tilde{D}'_{j,k}\right]\tilde{D}_{ii} \tag{D4}$$

where $\tilde{D}'_{jk}$ and $\tilde{D}'_{j,k}$ are the associations $\tilde{D}_{jk}$ and $\tilde{D}_{j,k}$ measured after selection. To the first order in $a_{\mathbb{U},\mathbb{V}}$ coefficients, $\tilde{D}_{i,jk} = 0$ at QLE while $\tilde{D}_{ij,j}$ and $\tilde{D}_{ij,k}$ are given by:

$$\tilde{D}_{ij,j} \approx \frac{1}{2}\, p_j q_j\, \tilde{D}_{ii}, \quad \tilde{D}_{ij,k} \approx \frac{1}{2}\left(\tilde{D}'_{jk} + \tilde{D}'_{j,k}\right)\tilde{D}_{ii}. \tag{D5}$$

Summing these expressions over all loci affecting the selfing rate, and using $V_\sigma = 2\sum_i \tilde{D}_{ii}$ and the expression for inbreeding depression measured after selection ($\delta'$) given by equation B9 in Supplementary File S2, equation D2 becomes:

$$\Delta_{\mathrm{s}}\,\overline{\sigma} = -\delta'\, V_\sigma + 2\sum_{i,j} a_j\left(\tilde{D}_{ij} + \tilde{D}_{i,j}\right) + 2\sum_{i,j<k} a_{jk}\,\tilde{D}_{ijk}\,. \tag{D6}$$

To the first order in $a_{\mathbb{U},\mathbb{V}}$ coefficients, $\tilde{D}_{ijk}$ at QLE is given by:

$$\begin{aligned}
\tilde{D}_{ijk} \approx\; & \frac{1}{\rho_{ijk} - \left(1 - \rho_{ijk}\right)\left[a_j\left(1 - 2p_j\right) + a_k\left(1 - 2p_k\right) + a_{jk}\left(1 - 2p_j\right)\left(1 - 2p_k\right)\right]} \\
& \times \Bigg[\Big[\left(1 - \rho_{jk}\right) a_{jk} + \rho_{jk}\, a_{j,k} + a_{jk,j}\left(1 - 2p_j\right) + a_{jk,k}\left(1 - 2p_k\right) \\
& \qquad + a_{jk,jk}\left(1 - 2p_j\right)\left(1 - 2p_k\right)\Big]\tilde{D}_{ijk,jk} \\
& + \Big[\left(1 - \rho_{jk}\right) a_{j,k} + \rho_{jk}\, a_{jk} + a_{jk,j}\left(1 - 2p_j\right)\Big] p_k q_k\, \tilde{D}_{ij,j} \\
& + \Big[\left(1 - \rho_{jk}\right) a_{j,k} + \rho_{jk}\, a_{jk} + a_{jk,k}\left(1 - 2p_k\right)\Big] p_j q_j\, \tilde{D}_{ik,k} \\
& + \frac{1}{2}\left(a_{jk} + 2\rho_{jk}\, a_{j,k}\right)\tilde{D}_{ii}\, pq_{jk}\Bigg]
\end{aligned} \tag{D7}$$

where $\rho_{ijk}$ is the probability that at least one recombination event occurs between the three loci $i$, $j$, $k$ during meiosis, and where $\tilde{D}_{ijk,jk}$, $\tilde{D}_{ij,j}$ and $\tilde{D}_{ik,k}$ are given by equations D3 and D5. Using equations A12 – A14 in Supplementary File S1, equation D7 yields equation 45 in the main text for the case of uniformly deleterious alleles with fixed epistasis (assuming that $p_j$, $p_k$ are small). In the case of stabilizing selection on quantitative traits, the terms in $a_{jk,j}$, $a_{jk,k}$ and $a_{jk,jk}$ in equation D7 all generate terms in $(1-2p_j)(1-2p_k)$ (see equations A25 – A26 and A54 – A55 in Supplementary File S1), that should vanish when summed over a large enough number of loci coding for the traits (due to the fact that two symmetric equilibria exist for $p_j$, one with $p_j < 1/2$ and the other with $p_j > 1/2$). This yields (using the fact that $a_{jk} = a_{j,k}$, and assuming that recombination rates are large relative to selection coefficients):

$$\sum_{j,k} a_{jk}\,\tilde{D}_{ijk} \approx \sum_{j,k} \frac{2+{\rho_{jk}}^2}{\rho_{ijk}}\,{a_{jk}}^2 pq_{jk}\,\tilde{D}_{ii}\,. \tag{D8}$$

Using the fact that $\sum_{j,k} {a_{jk}}^2 pq_{jk} \approx 4U^2/n$ independently of the parameter $Q$ describing the shape of the fitness peak (see equation A49 in Supplementary File S1), equation D8 yields equation 50 in the main text.

Finally, we have $\tilde{D}_{i,j} \approx 0$ when $\overline{\sigma} \approx 0$, while $\tilde{D}_{ij}$ is given by (to the first order in $a_{\mathbb{U},\mathbb{V}}$ coefficients):

$$\begin{aligned}\tilde{D}_{ij} \approx \frac{1}{\rho_{ij} - (1-\rho_{ij})\,a_j\,(1-2p_j)} \Bigg[ & [a_j + a_{j,j}\,(1-2p_j)]\,\tilde{D}_{ij,j} \\ & + \sum_k a_{jk,k}\,p_j q_j\,\tilde{D}_{ik,k} + \sum_k [a_{jk,k} + a_{jk,jk}\,(1-2p_j)]\,\tilde{D}_{ijk,jk}\Bigg].\end{aligned} \tag{D9}$$

Using equations D3 and D5, equation D9 yields equation 44 in the main text.
