## Supplementary File S5 for "Epistasis, inbreeding depression and the evolution of self-fertilization"

FILE S5: INDIVIDUAL-BASED SIMULATIONS

**Fixed epistatic effects.** The simulation program used to represent uniformly deleterious mutations with fixed epistatic effects is similar to those used in previous works (*i.e.*, Roze, 2009, 2015, 2016). The program represents a diploid population of constant size $N$ with discrete generations. The genome of each individual consists in two copies of a linear chromosome. Every generation, the number of new mutations per chromosome is drawn from a Poisson distribution with parameter $U$ (the deleterious mutation rate per haploid genome) and the position of each mutation along the chromosome is drawn from a uniform distribution (effectively allowing for a very large number of bi-allelic loci on each chromosome). A modifier locus affecting the selfing rate of individuals is located at the mid-point of the chromosome, with an infinite number of possible alleles coding for selfing rates between 0 and 1. Mutations occur at this locus at rate $U_{\text{self}}$ per generation, and the selfing rate coded by the new allele is drawn from a Gaussian distribution centered on the value of the allele before mutation, with standard deviation $\sigma_{\text{self}}$ (the new value is set to zero if it is negative, and to 1 if it is greater than 1). The fitness of each individual is computed using equation 9 in the main text, while the selfing rate $\sigma$ of the individual is given by the average of its two alleles at the modifier locus. The next generation is formed by first choosing a maternal parent. For each offspring, this is done by drawing a parent at random and retaining it as the maternal parent only if its fitness $W$ is greater than a random number drawn from a uniform distribution between 0 and 1 (otherwise another individual is sampled, until the test is satisfied). With a probability $\sigma$ the individual self-fertilizes and the maternal parent is also retained as the paternal parent. With a probability $1 - \sigma$ a second parent is selected as the paternal parent using the same procedure as for the maternal parent, except that the random number is compared with $W\,(1-\kappa\,\sigma)$ (where $W$ is the fitness of the individual and $\sigma$ its selfing rate) to take the effect of pollen discounting into account. Once the parents are sampled, meiosis takes place in both parents, the number of cross-overs being sampled from a Poisson distribution with parameter $R$ (genome map length), and the position of each cross-over along the chromosome being sampled from a uniform distribution. Inbreeding depression is estimated by comparing the mean fitnesses of 100 selfed offspring and 100 offspring produced by random mating (inbreeding depression after selection is measured by sampling parents according to their fitness as described above). At the start of the simulations individuals are free from deleterious mutation; during the first 20,000 generations the allele coding for $\sigma = 0$ is fixed at the modifier locus, then mutation is introduced at the modifier locus and the population is allowed to evolve during $2 \times 10^5$ generations, the mean selfing rate being recorded every 100 generations.

**Stabilizing selection.** Our simulation program is similar to the program used in Abu Awad and Roze (2018), representing $n$ polygenic traits under stabilizing selection. The genome of each individual consists of two copies of a linear chromosome carrying $\ell$ equidistant loci, represented by sequences of $\ell$ bits (0 or 1). Each locus has a probability $u = U/\ell$ of mutating (either from 0 to 1 or from 1 to 0) per generation. A table holds the phenotypic effects $r_{\alpha j}$ of changing the allelic state from 0 to 1 at each locus $j$, and is filled at the start of the simulation (each value is sampled independently from a centered Gaussian distribution with variance $a^2$). Although the program allows us to tune the degree of pleiotropy of mutations (number of traits $m$ affected by a given locus), in the present article we only considered full pleiotropy (every locus affects all $n$ traits). Every generation, the trait values $g_\alpha$ of each individual are computed according to its genotype (equation 13 in the main text); we assume no environmental effect on phenotypic traits (including environmental variance would decrease the effective strength of selection $V_s$, e.g., Lande, 1976). Individual fitnesses are then computed using equation 14 in the main text, $V_s$ being fixed to 10 in all simulation runs. The variance of mutational effects $a^2$ is adjusted to achieve a given mean deleterious effect of heterozygous mutations $\bar{\varsigma}$, using $\bar{\varsigma} = na^2/(2V_s)$ (Abu Awad and Roze, 2018). The selfing rate of individuals is determined by a set of $\ell_\sigma$ (fixed to 10) multi-allelic loci with additive effects. As in the program with fixed epistasis, the overall mutation rate at these loci is denoted $U_{\text{self}}$, while $\sigma_{\text{self}}$ is the standard deviation of mutational effects on selfing. The selfing rate of an individual is given by the sum of allelic values at the $\ell_\sigma$ loci (if the sum is above 1 or below 0, the selfing rate is truncated at 1 or 0, respectively). Selection and recombination are implemented as in the program with fixed epistasis. At the start of the simulation, allele 0 is fixed at each locus affecting phenotypic traits (so that all individuals are at the phenotypic optimum, $g_\alpha = 0$), while loci affecting the selfing rate are also fixed for allele 0 (random mating). The selfing rate stays at 0 during the first 20,000 generations, and is then allowed to evolve during the next 50,000 generations (the mean selfing rate and inbreeding depression being measured every 100 generations).
