## Supplementary Figures for "Epistasis, inbreeding depression and the evolution of self-fertilization"

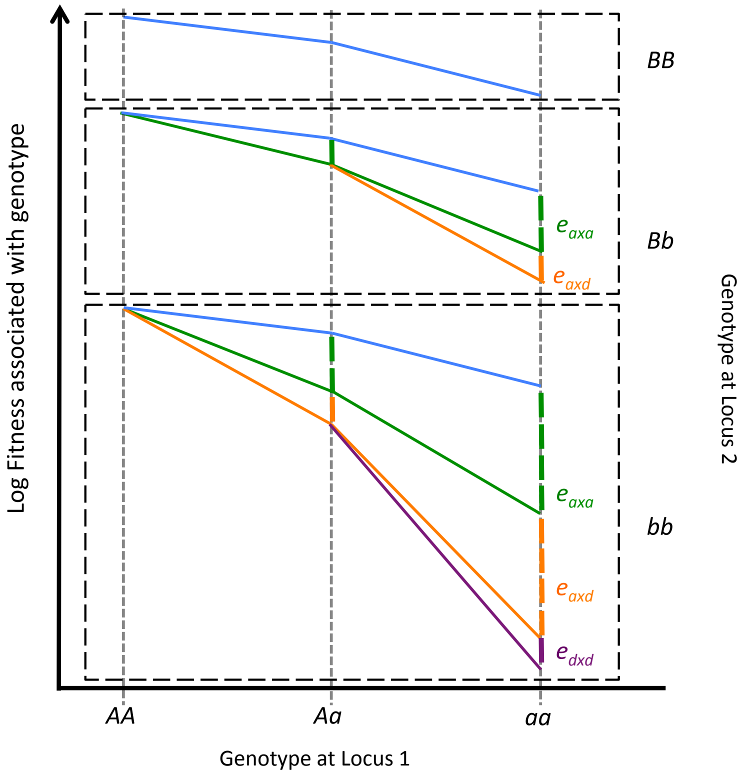


**Figure S1.** Graphical interpretation of the different components of epistasis in diploids. The $y$-axis shows the log-fitness of the different genotypes at two loci, where $a$ and $b$ are the deleterious alleles. Additive-by-additive epistasis ($e_{\mathrm{axa}}$) measures the effect of the interaction between two deleterious alleles at the two loci, additive-by-dominance epistasis ($e_{\mathrm{axd}}$) the extra effect of the interaction between three deleterious allele, and dominance-by-dominance epistasis ($e_{\mathrm{dxd}}$) the extra effect of the interaction between four deleterious alleles. Here these three forms of epistasis are negative.

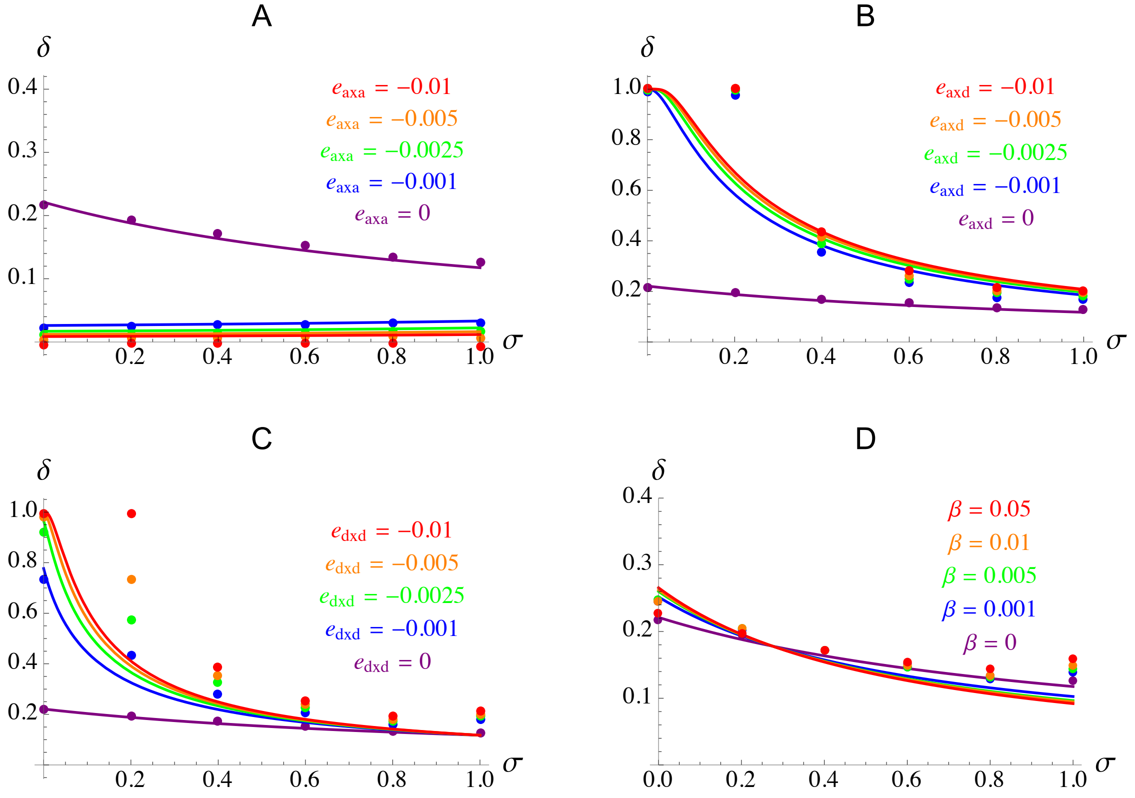


**Figure S2.** Same as Figure 1 in the main text, with $s = 0.01$.

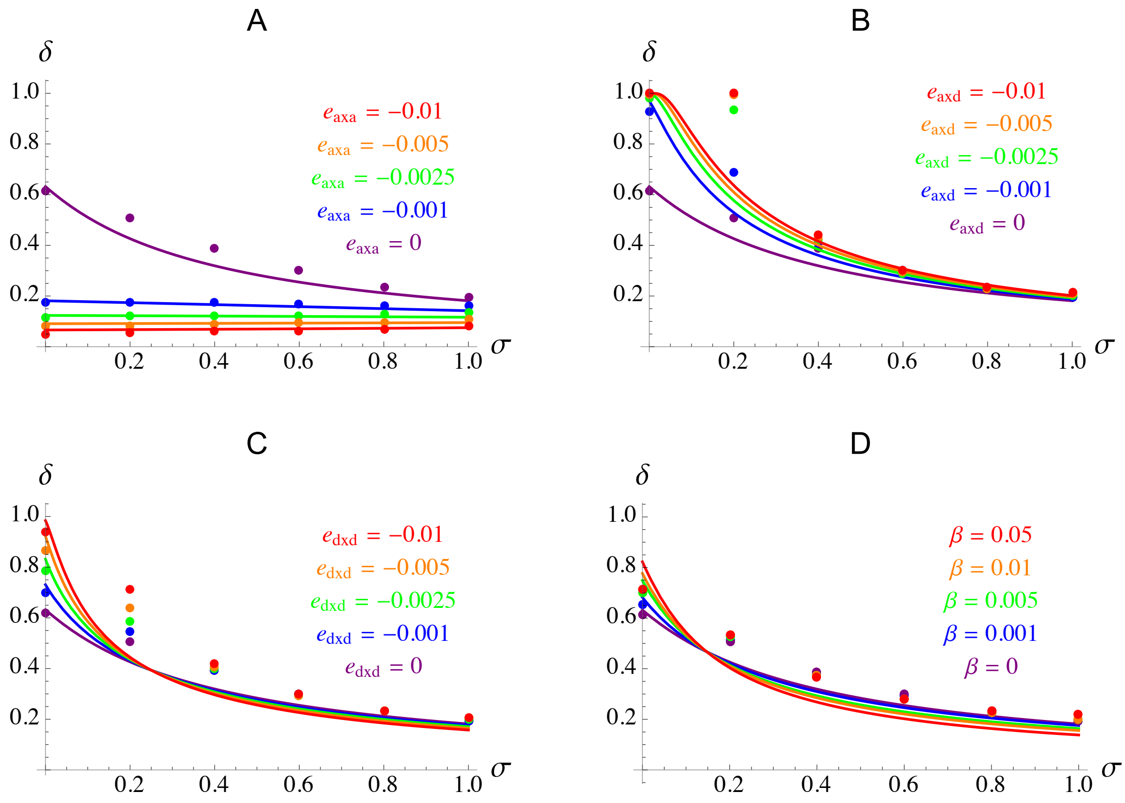


**Figure S3.** Same as Figure 1 in the main text, with $h = 0.1$.

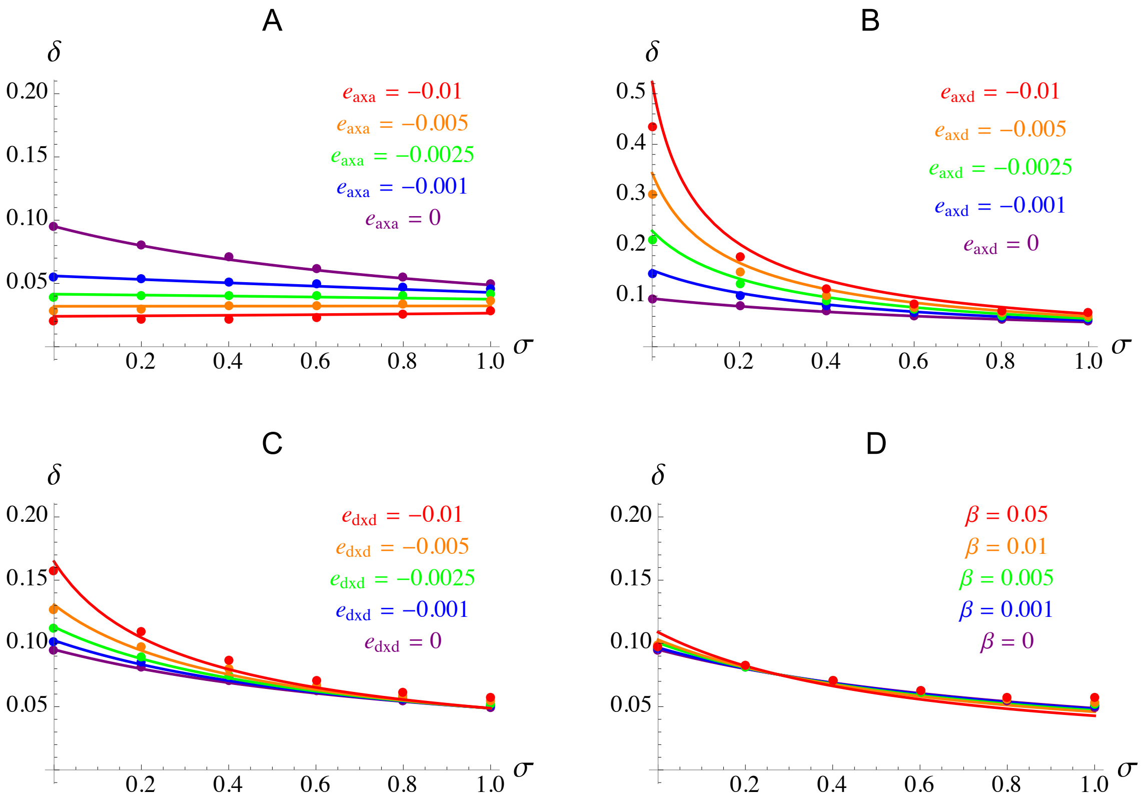


**Figure S4.** Same as Figure 1 in the main text, with $U = 0.1$.

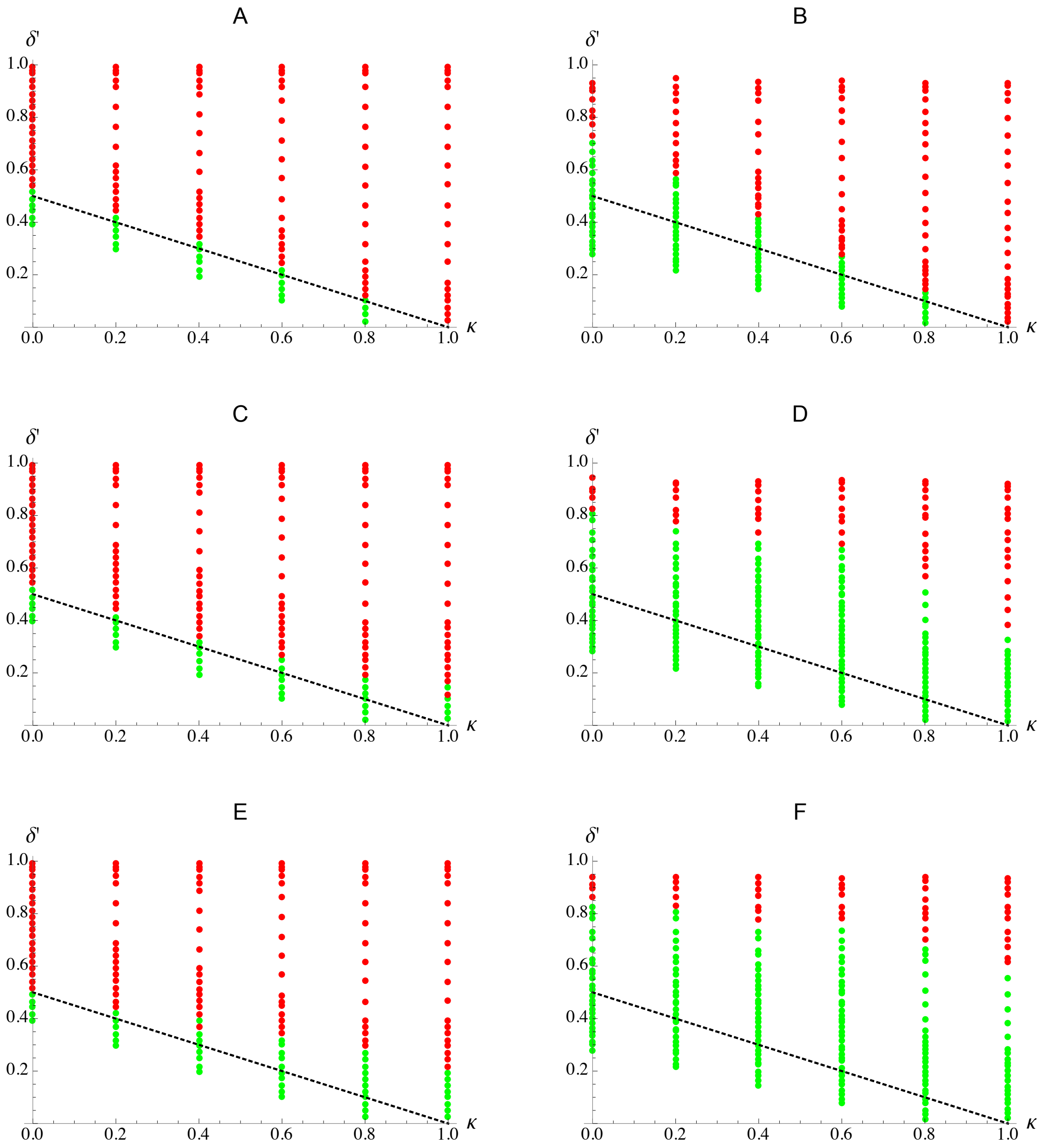


**Figure S5.** A, C, E: same as Figure 4A in the main text (no epistasis), with $\sigma_{\text{self}} = 0.03$ (A), $\sigma_{\text{self}} = 0.3$ (C), and mutation only between $\sigma = 0$ and $\sigma = 1$ in E. B, D, F: same as Figure 4E in the main text ($e_{\text{axa}}$, $e_{\text{axd}}$, $e_{\text{dxd}} < 0$), with $\sigma_{\text{self}} = 0.03$ (B), $\sigma_{\text{self}} = 0.3$ (D), and mutation only between $\sigma = 0$ and $\sigma = 1$ in F.

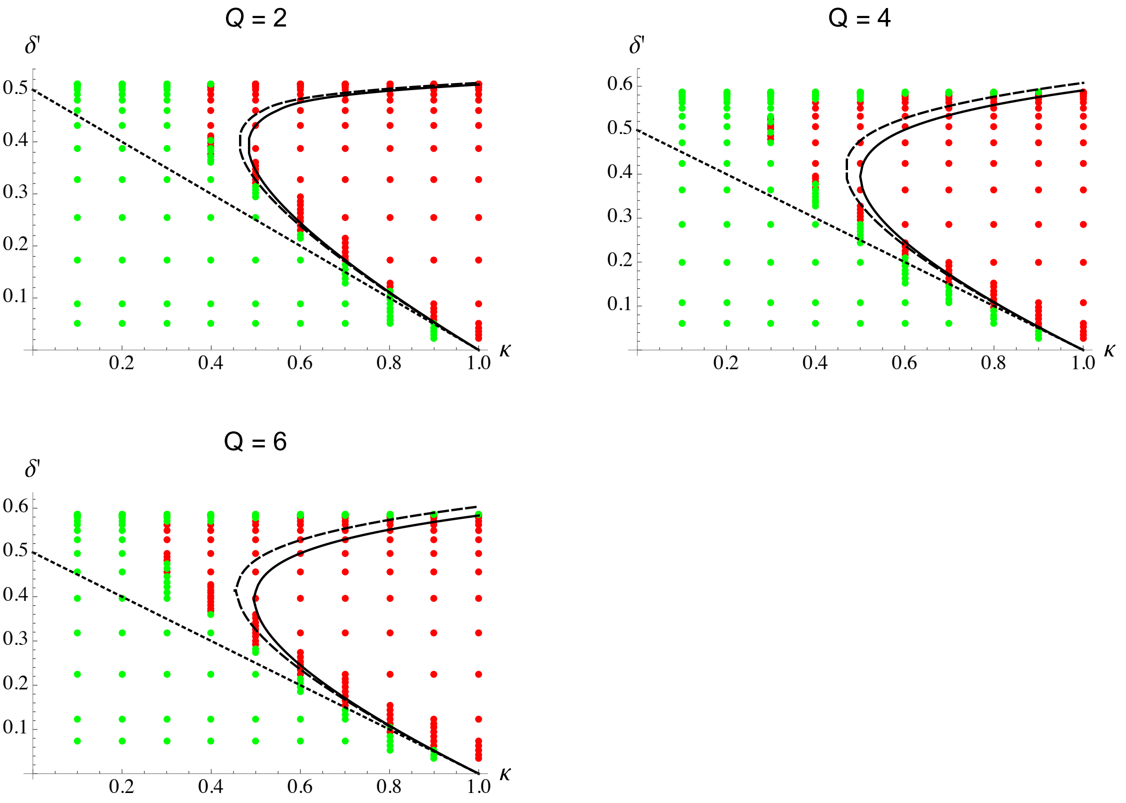


**Figure S6.** Same as Figure 5 in the main text with $n = 15$, for different values of the parameter $Q$ governing the shape of the fitness peak (equation 15 in the main text). Parameter values are the same as in Figure 5, the variance of mutational effects $a^2$ being set to $1/75$ (so that $\bar{\varsigma} = 0.01$ when $Q = 2$) in all cases.

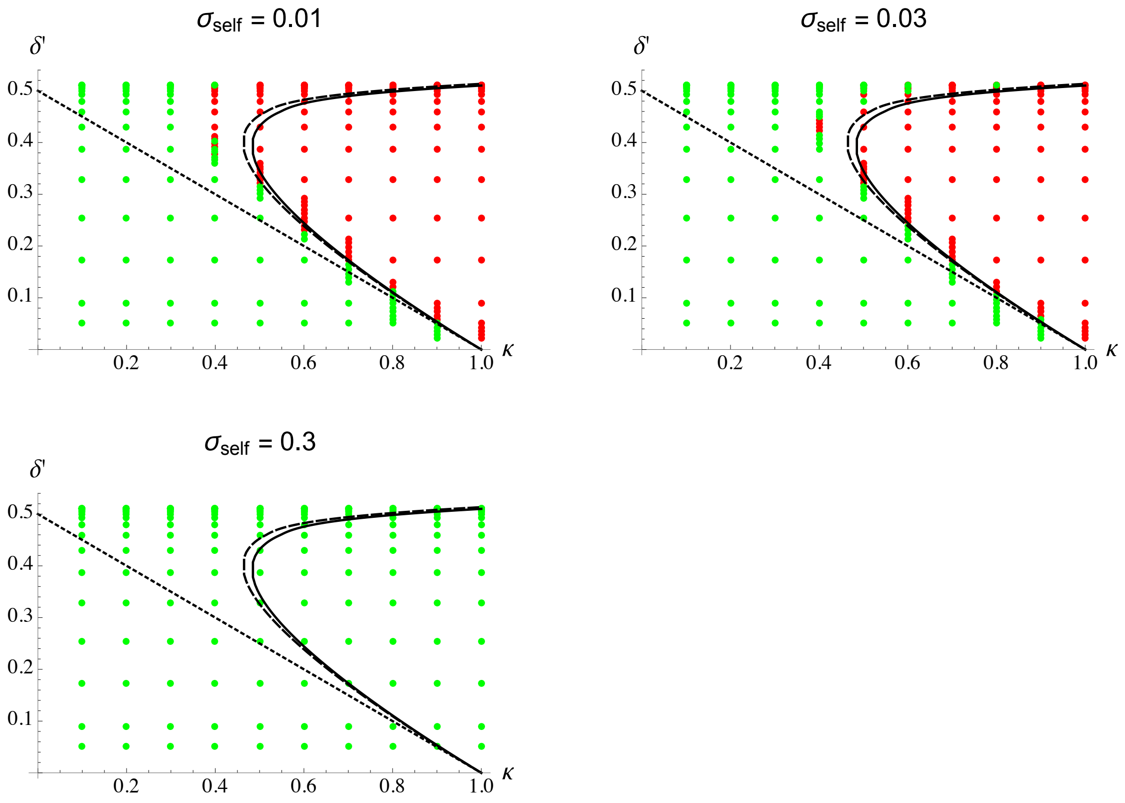


**Figure S7.** Same as Figure 5 in the main text with $n = 15$, for different values of the standard deviation of mutational effects at selfing modifier loci $\sigma_{\text{self}}$. Parameter values are the same as in Figure 5.
